# The extracellular matrix drives guanylate production and protects pancreatic cancer cells from oxaliplatin-induced DNA damage

**DOI:** 10.1101/2025.01.10.631286

**Authors:** Georgios Efthymiou, Coralie Compare, Eugénie Lohmann, Paraskevi Kousteridou, Pierre Bertrand, Heather S. Carr, Issam Ben-Sahra, Ivan Nemazanyy, Sarah Simha Tubiana, Zainab Hussain, Pascal Finetti, François Bertucci, Ghislain Bidaut, Christophe Lachaud, Fabienne Guillaumond, Richard Tomasini, Sophie Vasseur

## Abstract

The excessive production of extracellular matrix (ECM) and the metabolic adaptations in pancreatic ductal adenocarcinoma (PDAC) contribute individually to enhanced chemoresistance, dramatic tumor progression and dismal patient survival. However, ECM-driven metabolic alterations that promote chemoresistance in PDAC are so far unexplored. Here, we use *in-vitro*-generated ECM bio-scaffolds that recapitulate cell-ECM interactions and induce broad metabolic alterations in PDAC cells. High-throughput integration of multi-omics datasets coupled with metabolic tracing showed that the ECM enhances the generation of guanylates in PDAC cells, the accumulation of which alleviates oxaliplatin-induced DNA damage, and boosts PDAC cell proliferation. These events are guided by the guanosine monophosphate (GMP)-producing enzymes Impdh and Gmps, the expression of which correlated with that of matrisomal and DNA repair genes in PDAC patient samples. We propose that targeting ECM-driven metabolic processes, like the enhanced Impdh activity, may be an effective therapeutic approach for PDAC patients that bypasses the negative side effects of direct targeting of the ECM itself.

## Introduction

Pancreatic ductal adenocarcinoma (PDAC) is an inherently aggressive disease characterized by strong resistance to current therapeutic schemes. The extracellular matrix (ECM) greatly contributes to drug resistance and PDAC progression resulting in a dismal 5-year relative survival of 12.8% (*1*). Upon activation by tumor-cell-derived signals, myofibroblastic-like cancer-associated fibroblasts (myCAFs) become the prominent ECM producers within the tumor generating excessive amounts of collagen, fibronectin, laminin, and hyaluronic acid (*2–5*). This process, known as desmoplasia (*2*, *6*), plays a central role in the formation of a dense and rigid “activated” stromal compartment around the tumor cells (*6*) that accounts for up to 90% of the tumor mass (*7*) and is associated with poor prognosis (*8*). The rigidified ECM forms a barrier around the tumor cells, causes blood vessel collapse, and results in a gradient of oxygen concentration and nutrient availability to the tumor cells (*9*, *10*). Under these unfavorable conditions, PDAC cells deploy metabolic reprogramming that maintains the levels of metabolic intermediates needed for proliferation. Nucleotides are essential components for anabolic processes that support the aggressive behavior of rapidly proliferating PDAC cells (*11*, *12*). Indeed, low-differentiated quasi-mesenchymal PDAC cells preferentially promote aerobic glycolysis to provide the pentose phosphate pathway with the necessary carbon for nucleotide synthesis (*13*, *14*). Similarly, addiction to glutamine provides ample nitrogen for the de novo generation of nucleotides and is correlated with poor prognosis (*13*).

Over the past decade, few studies have focused on the ECM as a driver of cancer cell metabolism. Extracellular protein catabolism has been demonstrated to provide PDAC cells with numerous nutrients under nutrient-limited conditions (*15*, *16*). Interestingly, our team has shown that PDAC cells degrade collagen from the ECM and use collagen-derived proline to feed the TCA cycle, thus promoting tumor cell survival and PDAC growth (*17*). Similarly, hyaluronic acid, an abundant component of PDAC ECM (*18*), has been described as a fuel for the hexosamine biosynthetic pathway supporting PDAC growth (*19*). Apart from a nutrient pool, the ECM presents a biochemical scaffold, the remodeling of which may drive multiple cellular processes. Indeed, remodeling of the hyaluronic acid network was shown to enable the interaction between cell receptors and ECM-bound factors, inducing an acute glycolytic response in PDAC cells lines (*20*). Whether this resulted in enhanced PDAC tumorigenicity is not known. Cell-ECM interactions and ECM-relayed cues are numerous, highly complex and depend on the spatial properties of the ECM topology and the residing cells (*21–23*). Exploration of these interactions will help us understand their effect in PDAC physiology, progression, and chemoresistance.

Oxaliplatin (OX) is a major antineoplastic drug used routinely for the treatment of unresectable PDAC patients in combination with fluorouracil (5-FU) and irinotecan (in a drug cocktail known as FOLFIRINOX). Tumor resistance to OX is a major challenge in PDAC patients who often develop rapid tumor recurrence with extensive ECM remodeling, in association with significant decrease in survival (*24–28*). This is attributed, at least in part, to the chemotherapy-induced changes in the function of CAFs and their role in the production and remodeling of the local ECM (*24*). Neo-adjuvant oxaliplatin-containing chemotherapy induces a dramatic alteration in the distribution of CAF phenotypes in colorectal cancer (accumulation of ECM-producing myofibroblasts, decreased presence of ECM remodeling CAFs) resulting in a pronounced fibrotic response (*29*). Similar results were observed in pancreatic cancer where chemotherapy increased the frequency of sub-tumor microenvironments (subTMEs) characterized by thin, spindle-shaped fibroblasts and mature ECM fibers with chemoprotective properties (*30*). Despite the established role of the ECM as a conductor of tumorigenicity, and of metabolism as a cancer hallmark (*31*, *32*), ECM-driven metabolism and its effect in PDAC chemoresistance is an uncharted territory. Indeed, standard cell culture studies often use non-physiologically relevant or variable ECM formulations (coatings, soluble) that may mask the full range of ECM-relayed cues [reviewed in (*33*)].

To address this issue, we implemented a straightforward, highly robust *in vitro* model consisting of the culture of PDAC cells on CAF-derived ECM scaffolds that mimic the ECM compartment of PDAC tumors. We combined this biologically relevant system with high throughput bioinformatic integration of RNA sequencing (RNAseq), metabolomics, and steady state metabolic tracing in PDAC cells. We show that the ECM favors the production and accumulation of guanylates in PDAC cells, and this is closely linked with their enhanced capacity to repair oxaliplatin-induced DNA lesions and maintain a high rate of proliferation. We identified Impdh and Gmps as the enzyme(s) responsible for the accumulation of guanylates and for the capacity of cancer cells to repair OX-induced DNA lesions. This study puts forth the presence of an ECM – metabolism – DNA repair axis, each part of which may be specifically targeted in combination with standard chemotherapy regimens, paving the way for the development of tailored therapeutic strategies against PDAC.

## Results

### CAF ECM induces purine imbalance in PDAC cells

To understand ECM-modulated metabolism in PDAC, we implemented a cell-on-matrix *in vitro* system where murine PDAC cells (hereafter PK4A) were cultivated on the ECM produced by PDAC-patient-derived CAFs (Fig. 1A). This CAF-derived matrix (CDM) represents a bio-scaffold with structural and topological features that mimic PDAC ECM (*34*) and recapitulates cell-ECM interactions [(*35*, *36*) and references therein]. Within this setting, we utilized human plasma-like medium (HPLM), which unlike synthetic formulations, contains physiologically relevant concentrations of nutrients (*37*) allowing us to achieve a more accurate metabolite make-up of PDAC. For the generation of the CDM, we tested two patient-derived CAF populations that corresponded to the activated resident myo-fibroblast-like CAF subtype, the major ECM producer in PDAC (*6*). Upon hTERT (human telomerase reverse transcriptase)-immortalization (CAFi), both populations expressed the typical CAF markers integrin beta 1 (ITGB1), platelet-derived growth factor receptor B (PDGFRB), fibroblast activation protein (FAP) (Fig. S1A). CAFi 2, however, deposited an elaborate provisional fibronectin (FN) network with thin long fibrils and coarse short ones (Fig. S1B right), that correspond to FN deposition and fibril maturation respectively. In contrast, CAFi 1 displayed mostly coarse FN fibers with high intensity nodes (Fig. S1B left), suggesting a fragmented ECM. This is in line with the lower expression of alpha-smooth muscle actin [αSMA, a marker of CAF activation (*4*, *6*)] (Fig. S1C) in CAFi 2, and the lower collagen gel contraction (Fig. S1D), a proxy of ECM remodeling and turnover (*38*) in these cells. Finally, unlike CAFi 1, the number of viable CAFi 2 cells significantly increased over 3 days of culture (Fig. S1E), and they were therefore selected for the generation of the CDM.To explore the ECM-driven metabolic adaptations of PDAC cells, we used two batches of PK4A cells, each isolated from the PDAC of a KIC (Pdx-Cre; LSL-Kras^G12D/+^; Ink4a/Arf^flox/flox^) mouse. We performed RNAseq and semi-targeted metabolomics in cells cultivated on CDM and on CDM-free tissue culture plastic plates (TCP), and we integrated the comparative datasets into a joint-pathway analysis (Fig. 1A). Among the most significantly deregulated metabolic pathways common in both PK4A cell batches (Fig. 1B, S1F, Tables S1, 2), purine metabolism had the highest cumulative impact and p-value (Fig. S1G). Interestingly, guanine (Gua) and guanosine (Guo) were significantly upregulated in CDM compared to TCP (Fig. 1C, D top), while no difference was observed for adenine (Ade) or adenosine (Ado) (Fig. 1C, Table S3). Inosine monophosphate (IMP) and inosine (Ino) were also upregulated in PK4A cells on CDM (Fig. 1C). IMP is converted to guanosine monophosphate (GMP) and adenosine monophosphate (AMP) via the *de novo* purine biosynthesis, while Ino may give rise to AMP or GMP via multiple steps that include the salvage pathway (Fig. 1D bottom). This simultaneous increase of IMP, Ino, Guo and Gua in cancer cells grown on CDM with no concomitant rise of Ado or Ade indicates an imbalance in cellular purine pools and suggests a shift of purine production towards GMP. Interestingly, half of the significantly deregulated purine metabolism genes (Table S4) are involved in the production and turnover of guanylates, while only a few are associated with adenylates or both (Fig. 1E). Among these, guanine deaminase (*Gda*) downregulation may impede guanylate degradation in CDM-grown PK4A cells, while the upregulation of inosine monophosphate dehydrogenase 1 (*Impdh1*) with the simultaneous downregulation of guanosine monophosphate reductase (*Gmpr*) may promote GMP production (Fig. 1D bottom, E). Furthermore, adenine phosphoribosyltransferase (*Aprt*) and adenosine monophosphate deaminase (*Ampd3*) overexpression in cells cultivated on CDM may deplete adenylate pools for IMP re-generation (Fig. 1D bottom, E). Of note, the expression of genes involved in pyrimidine metabolism remained unaltered between cells grown on CDM and TCP (Fig. S1H, Table S5), and no concentration imbalance was observed between uracil– and cytosine-containing compounds in cells grown on CDM (Fig. S1I, Table S6). Finally, the substrate rigidity perceived by the cells grown on TCP and CDM was similar, indicated by the similar nuclear localization of the mechanosensor YAP1 (Yes-associated protein 1) in both conditions (Fig. S1H). Together, these data suggest that the CDM modulates a mechanosensing-independent metabolic alteration in PDAC cells, where the cellular concentration of guanine-containing compounds is increased.

**Fig. 1.**
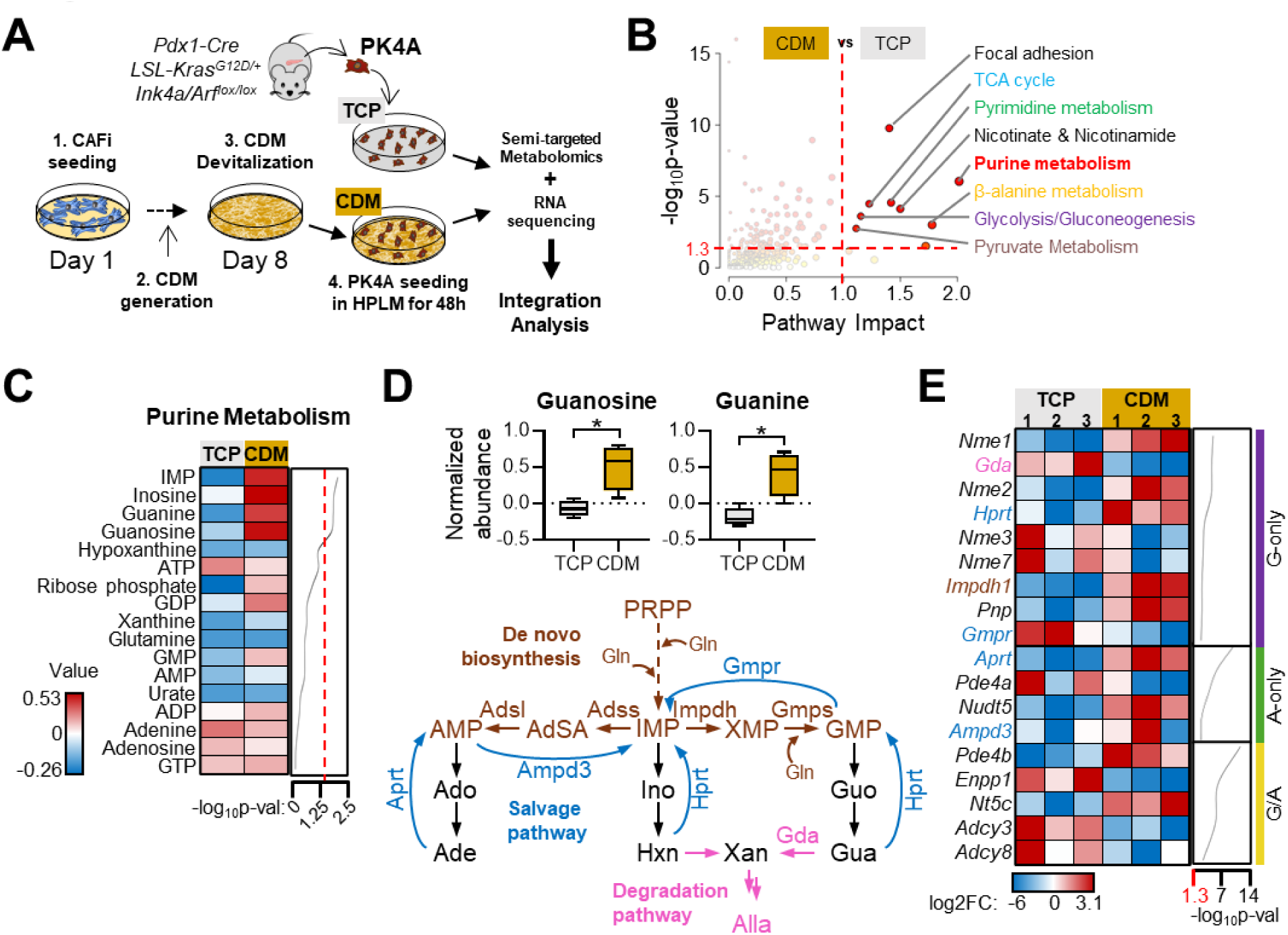
CAF-derived ECM drives purine imbalance in PDAC cells. **(A)** Schematic representation of the in vitro model used in the study. Cancer associated fibroblasts (CAF), isolated from PDAC patient biopsies and immortalized by hTERT overexpression (CAFi) were cultivated for 8 days in the presence of L-ascorbic acid to generate CAF-derived matrices (CDM). PDAC cells (PK4A) from a KIC mouse model were seeded on decellularized CDM or on CDM-free tissue culture plates (TCP) for 48h in human plasma-like medium (HPLM) for the generation of RNAseq and semi-targeted metabolomics datasets (CDM vs TCP). **(B)** Integration of the matched RNAseq and metabolomics datasets presented as a dot plot. Each dot represents a KEGG metabolic pathway, and the dot size reflects the number of identified features (genes/metabolites) within that pathway. A threshold of 1.3 was applied for –log_10_p val (red horizontal dotted line), and 1.0 for pathway impact (red vertical dotted line). **(C)** Heatmap representation of Purine Metabolism Compounds in PK4A cells cultivated on TCP and on CDM for 48h in HPLM. Cumulative data from two independent experiments in two batches of PK4A cells. Compounds are ranked by significance with the –log_10_p value shown on the right. The dotted line delineates the threshold of significance (-log_10_p ≥ 1.3). **(D)** Top: Normalized abundance of guanosine and Guanine n PK4A cells cultivated for 48h in HPLM on top of CDMs or on TCP. Cumulative data from two independent experiments in two batches of PK4A cells. Unpaired parametric *t* test. * p≤0.05. Bottom: Simplified scheme of purine metabolism. The *de novo* biosynthetic pathway is shown in brown, the salvage in blue, and the degradation pathway in pink. PRPP, phosphoribosylpyrophosphate; Gln, glutamine; IMP, inosine monophosphate; Ino, inosine; Hxn, hypoxanthine; XMP, xanthosine monophosphate; Xan, xanthine; GMP, guanosine monophosphate; Guo, guanosine; Gua, guanine; Alla, allantoin; AdSA, adenylosuccininc acid; AMP, adenosine monophosphate; Ado, adenosine; Ade, adenine. Impdh, inosine-5’-monophosphate dehydrogenase; Gmps, guanosine monophosphate synthase; Adss, adenylosuccinate synthase; Adsl, adenylosuccinate lyase; Hprt, hypoxanthine-guanine phosphoribosylatransferase; Gmpr, guanosine monophosphate reductase; Aprt, adenine phosphoribosyltransferase; Ampd3, adenosine monophosphate deaminase 3; Gda, guanine deaminase. **(E)** Heatmap representation of significantly deregulated purine metabolism genes (-log_10_p ≥ 1.3) identified and extracted from the integration analysis in (D). The genes are divided into three groups: involved in the metabolism of guanylates only (purple, top), of adenylates only (green, middle) or both (yellow, bottom), and ranked by decreasing –log_10_p value shown on the right.

### CDM drives guanylate accumulation and oxaliplatin resistance in PDAC cells

Guanylate accumulation has previously been associated with the repair of radiation-induced DNA damages in cancer cells thus promoting chemoresistance and cell survival (*39*). Thus, we next addressed whether the CDM potentiates the metabolic flux of the *de novo* purine biosynthetic pathway towards an enhanced production of guanylates to promote chemoprotection of PDAC cells against oxaliplatin (OX) treatment. To that end, we traced the incorporation of glutamine-amide-^15^N (^15^N-Gln) in the purines of OX-treated PK4A cells grown on CDM and on TCP (Fig. 2A). As expected, the CDM significantly enhanced the incorporation of Gln-derived ^15^N in GMP (GMP M+3) upon OX treatment in PDAC cells (Fig. 2B left), but it did not restore the OX-mediated decrease in Gln-derived nitrogen incorporation into AMP (AMP M+5) (Fig. 2B right). This indicates that the overall contribution of the *de novo* biosynthetic pathway was enhanced towards the production of GMP but not of AMP in OX-challenged PK4A cells cultivated on CDM. Similarly, we determined the contribution of the salvage pathway to the production of GMP and AMP by using U-^13^C-hypoxanthine (U-^13^C-Hxn) (Fig. 2A). We observed that Hxn-derived carbon incorporation in IMP (IMP M+5) (Fig. 2C left) and AMP (AMP M+5) (Fig. 2C center) was considerably lower in OX-treated PDAC cells grown on CDM. Intriguingly, the M+5 fraction of GMP remained considerably high in these cells (Fig. 2C left). These data suggest that despite the reduced HPRT (hypoxanthine-guanine phosphoribosyltransferase)-mediated purine salvage, GMP production remains intact in OX-treated PK4A cells grown on CDM. Taken together, these results suggest that the CDM enhances the rate of IMP-to-GMP conversion in OX-treated PDAC cells, a process mediated by Impdh and Gmps. Consistent with the tracing results, the mRNA levels of both isoforms of Impdh (Impdh1 and Impdh2), the enzyme that catalyzes the conversion of IMP to xanthosine monophosphate (XMP) (Fig. 2A), were significantly upregulated in OX-treated PK4A cells in a CDM-dependent manner (Fig. 2D). Conversely, the expression of adenylosuccinate synthase (*Adss*) and adenosylsuccinate lyase (*Adsl*), involved in the AMP-producing arm of the *de novo* purine biosynthesis, remained unchanged in all conditions tested (Fig. S2A). These data demonstrate that the ECM strongly favors the activity of the GMP-generating arm of the *de novo* purine biosynthetic pathway in OX-challenged PK4A cells grown on CDM, thus contributing to the overrepresentation of guanylates in these cells. Interestingly, while the addition of OX resulted, as expected, in a significant decrease in the number of viable cells grown on TCP, this effect was completely reversed by the presence of the CDM in both batches of PK4A cells (Fig. 2E, F and Fig. S2B, C). This protective feature of the CDM against OX was lost when PK4A cells were cultured in the presence of a homogenized CDM (hCDM) preparation (Fig. 2E and Fig. S2B dotted bars). These results suggest that the observed CDM-driven chemoprotection relies on the spatial distribution and the high-order assembly of the ECM components that trigger pro-tumoral cellular processes, including the shifts in purine production and chemoresistance.

**Fig. 2.**
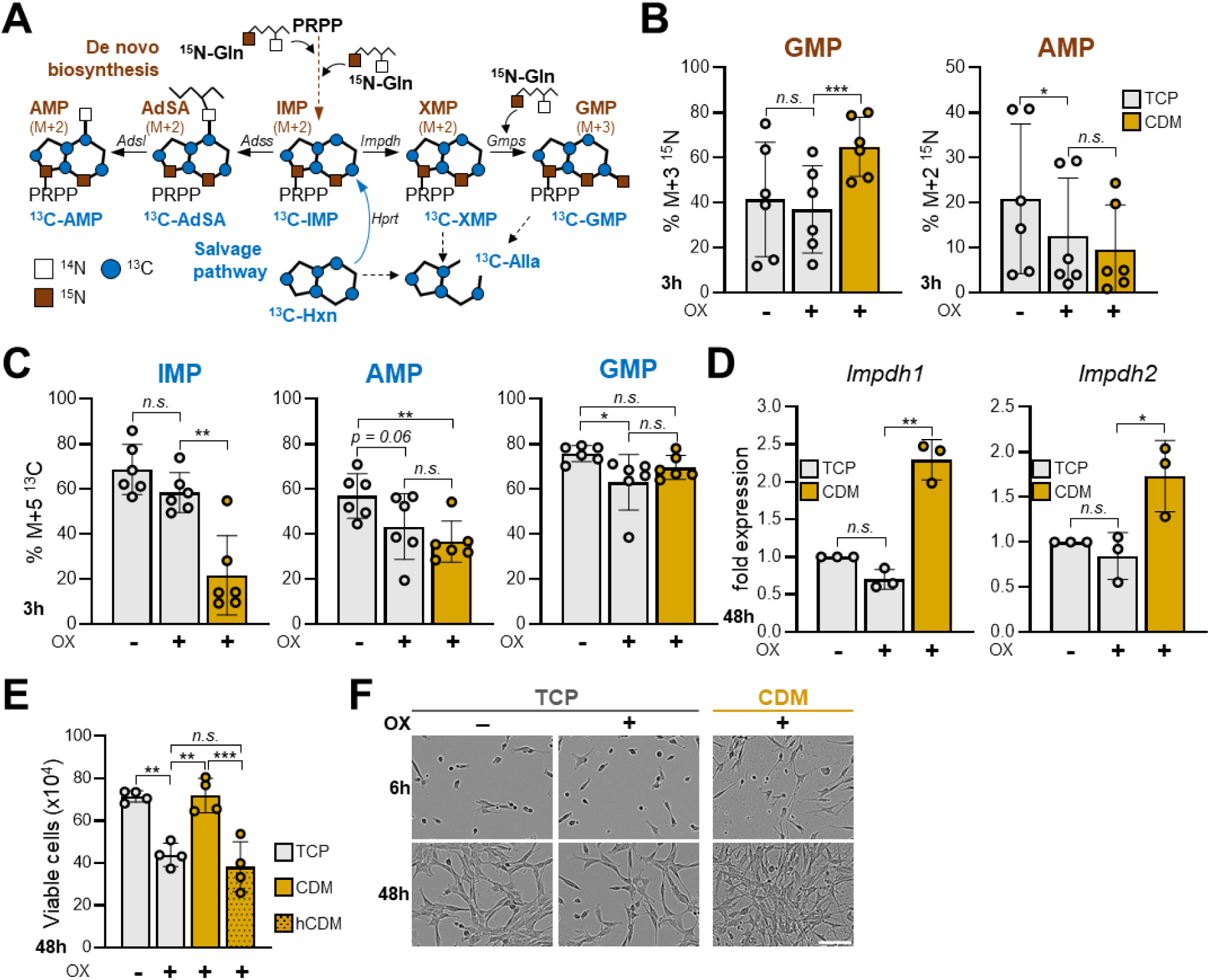
CDM drives GMP production and oxaliplatin resistance in PDAC cells. **(A)** Schematic representation of glutamine-derived nitrogen (brown squares) incorporation into purines via the de novo biosynthetic pathway (brown), and of hypoxanthine-derived carbon (blue circles) via the salvage pathway (blue). **(B)** Graph demonstrating the incorporation of three Gln-derived ^15^N in GMP (M+3), or two Gln-derived ^15^N in AMP (M+2) in PK4A cells cultivated on CDM and on TCP for 3h in HPLM containing 4mM of amido-^15^N-glutamine (^15^N-Gln) and 0.5μM of OX. Data are represented as mean ±SD from three independent experiments in two batches of PK4A cells. Repeated measures one-way ANOVA with Tuckey’s multiple comparisons test, with a single pooled variance. *** p≤0.001; * p≤0.05; not significant. **(C)** Graph demonstrating the incorporation of Hxn-derived ^13^C (M+5) in IMP, GMP, and AMP in PK4A cells cultivated on CDM and on TCP for 3h in HPLM containing 100μM of ^13^C-hypoxanthine (^13^C-Hxn) and 0.5μM of OX. Data are represented as mean ±SD from three independent experiments in two batches of PK4A cells. Repeated measures one-way ANOVA with Tuckey’s multiple comparisons test, with a single pooled variance. ** p≤0.01; * p≤0.05; n.s., not significant. **(D)** Expression of *Impdh1* and *Impdh2* mRNA in PK4A cells grown on CDM compared to TCP in HPLM in presence of 0.5μM of OX for 48h. Data are represented as fold change relative to TCP (TCP+OX vs TCP; CDM+OX vs TCP+OX) from three independent experiments. Ordinary one-way ANOVA. ** p≤0.01. **(E)** Bar graph displaying the number of PK4A cells grown on TCP or on CDM after 48h in HPLM with addition of 0.5μM of OX. hCDM signifies a homogenized CDM preparation presented to the cells concurrently with cell seeding and OX addition. Data are presented as mean ±SD of four independent experiments. Ordinary one-way ANOVA, with Tukey’s multiple comparisons test, with a single pooled variance. *** p≤0.001; ** p≤0.01; n.s., not significant. **(F)** Phase contrast images of PK4A cells cultivated on TCP and on CDM in HPLM for 6h (top), or 48h (bottom) in the presence of 0.5μM of oxaliplatin (OX). Scale bar, 50μm.

To further examine the role of the CDM-driven production and accumulation of guanylates in PK4A cell proliferation, we utilized mizoribine, a potent inhibitor of Impdh and Gmps (GMP synthase) (Fig. 1D bottom). As expected, the addition of mizoribine to PK4A cells upon seeding dramatically decreased the number of viable cells both on CDM and on TCP substrates (5-fold and 3-fold decrease compared to corresponding non-treated cells, respectively) (Fig. 3A). The addition of GMP significantly reversed the effect of mizoribine independently of the substrate, while the addition of Guo restored the number of viable cells only when they were cultivated on CDM (Fig. 3A). Next, we cultivated PK4A cells on CDM and on TCP for 24h to allow for guanylates to accumulate in cells grown on CDM, and then we inhibited the GMP-generating branch of the *de novo* biosynthetic pathway for another 24h (Fig. 3B). As expected, the addition of mizoribine resulted in approximately 50% decrease in the number of viable PK4A cells grown on TCP, while PK4A cells grown on CDM were less sensitive to Impdh/Gmps inhibition and reached a number comparable to that of control TCP (Fig. 3B). These data suggest that during the first 24h the CDM supported the accumulation of guanylates, which significantly counteracted the inhibitory effect of mizoribine, and maintained the proliferative capacity of these cells. The CDM-mediated guanylate accumulation is further supported by the significantly lower Gda mRNA (Fig. 1E) and protein levels (Fig. 3C). Gda converts guanine to xanthine for degradation, which was also considerably lower in OX-treated cells grown on CDM compared to those cultivated on TCP. This was assessed by the Hxn-derived ^13^C incorporation in allantoin (Alla), the final product of purine degradation in the mouse (Fig. 2A, 3D).

**Fig. 3.**
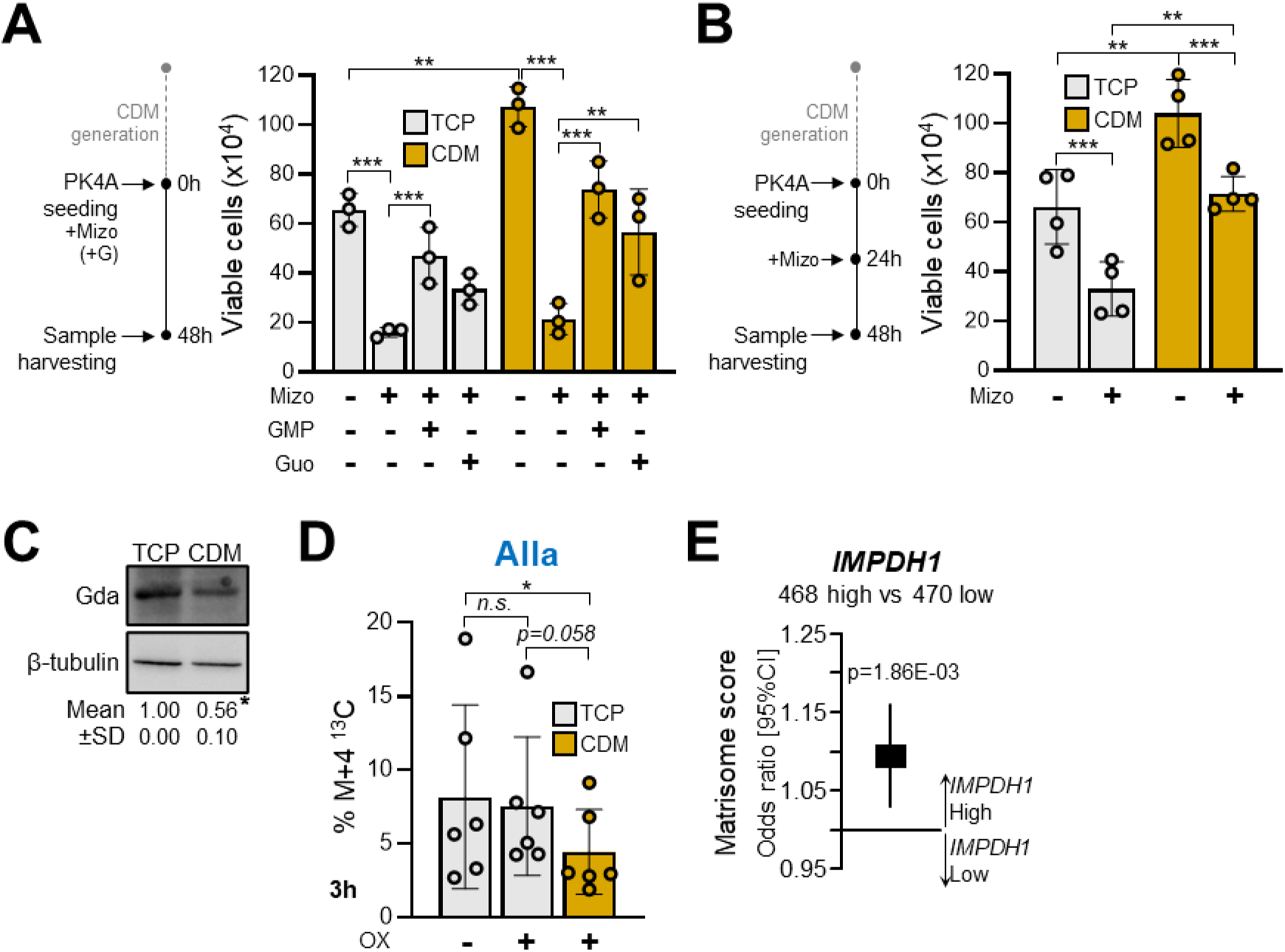
CDM promotes guanylate production and accumulation in PDAC cells. **(A)** Experimental design (left) and bar graph representation (right) of the number of PK4A cells grown on TCP or on CDM after 48h in HPLM in the absence or presence of 2.5μM of mizoribine (Mizo), a specific inhibitor of Impdh and Gmps. Mizoribine was added concurrently with cell seeding, as were GMP and Guo as means of rescuing mizoribine-mediated Impdh/Gmps inhibition. Data are presented as mean ±SD from three independent experiments. Ordinary two-way ANOVA, with Tukey’s multiple comparisons test, with individual variances computed for each comparison. **** p≤0.0001; *** p≤0.001; ** p≤0.01; * p≤0.05; not significant comparisons were omitted for clarity. **(B)** Experimental design (left) and bar graph representation (right) of the number of PK4A cells grown on TCP or on CDM after an initial incubation for 24h, addition of 2.5μM of mizoribine 48h and an additional incubation for another 24h. Data are presented as mean ±SD of four independent experiments. Ordinary two-way ANOVA, with Tukey’s multiple comparisons test, with individual variances computed for each comparison. ** p≤0.01; * p≤0.05; not significant comparisons were omitted for clarity. **(C)** Western blot showing the decreased expression of deaminase (Gda) in PK4A cells grown on CDM compared to TCP. β-tubulin was used as loading control. The numbers beneath the blots express the mean fold decrease ±SD from 3 independent experiments. Student’s *t* test. * p≤0.05. **(D)** Ratio of the ^13^C-containing fractions of allantoin in PK4A cells cultivated on CDM and on TCP for 3h in HPLM containing 100μM of ^13^C-hypoxanthine (^13^C-Hxn) and 0.5μM of OX. Data are represented as mean ±SD from three independent experiments in two batches of PK4A cells. Repeated measures one-way ANOVA with Tuckey’s multiple comparisons test, with a single pooled variance. * p≤0.05; n.s., not significant. **(E)** Forest plot representation of the correlation between the matrisome metagene and *IMPDH1* expression classes in the 938 PDAC patient samples. Odds ratio (box) are indicated with confidence interval (line). P-value of the logistic regression was estimated by specifying a binomial family for model with a logit function.

Taken together, these results demonstrate that the increased cellular guanylate pools in PDAC cells are due in part to the ECM-controlled enhancement of the GMP-generating arm of the *de novo* biosynthetic pathway, through the activity of Impdh and/or Gmps, together with the decrease of guanine degradation. Interestingly, analysis of gene expression data from 938 primary PDAC samples showed a higher expression of matrisomal genes in *IMPDH1*-high than in *IMPDH1*-low tumors (Fig. 3E). This suggests that IMPDH-mediated guanylate production may be associated with specific features of PDAC ECM.

### CDM promotes the repair of OX-induced DNA lesions in an Impdh/Gmps-dependent manner

We showed that the CDM promotes the production and accumulation of guanine-containing compounds in PDAC cells while protecting them from OX treatment. OX is a platinum-based compound that binds and forms adducts in the DNA of rapidly proliferating cells causing extensive DNA damage which in turn results in polymerase stalling, replication stress, and ultimately, to apoptotic signaling and cell death [reviewed in (*25*, *40*)]. To assess the capacity of PK4A cells to repair OX-induced DNA damage when cultivated on CDM, we quantified γH2AX, a phosphorylated form of H2A Histone family member X, which is an indicator of DNA double strand breaks (DSB) (*41*). As expected, addition of OX in PK4A cells resulted in a significant increase in γH2AX intensity compared to non-treated cells, which was partly but significantly alleviated in OX-treated PK4A cells seeded on CDM (Fig. 4A). Differential gene expression analysis revealed that 42% of a DNA repair gene set (*42*) was significantly upregulated in PK4A cells grown on CDM compared to those cultivated on TCP (Fig. 4B and Table S7). Interestingly, these genes are involved in nucleotide excision repair (NER), as revealed by gene set enrichment analysis (Fig. 4C and Table S8). NER is a mechanism implemented by cancer cells to eliminate DNA lesions induced by chemotherapeutic agents, including platinum salts (*43*, *44*). Key genes involved in NER were significantly upregulated in OX-treated PK4A cells specifically grown on CDM, including the endonuclease component *Ercc1*, the transcription factor *Gtf2h1*, the lesion recognition component *Rad23*, and the polymerases *Pold4* and *Polb* (Fig. 4D). In addition, the NER negative regulator *Parp9* was downregulated in these cells (Fig. 4D). These results are in line with previous works reporting that cancer cells acquire resistance to OX by promoting NER [reviewed in (*26*)]. Since OX alone did not alter the expression of any of these genes in cells grown on TCP (Fig. 4D), our findings suggest the critical role of the ECM as a driver of OX resistance in PDAC.

**Fig. 4.**
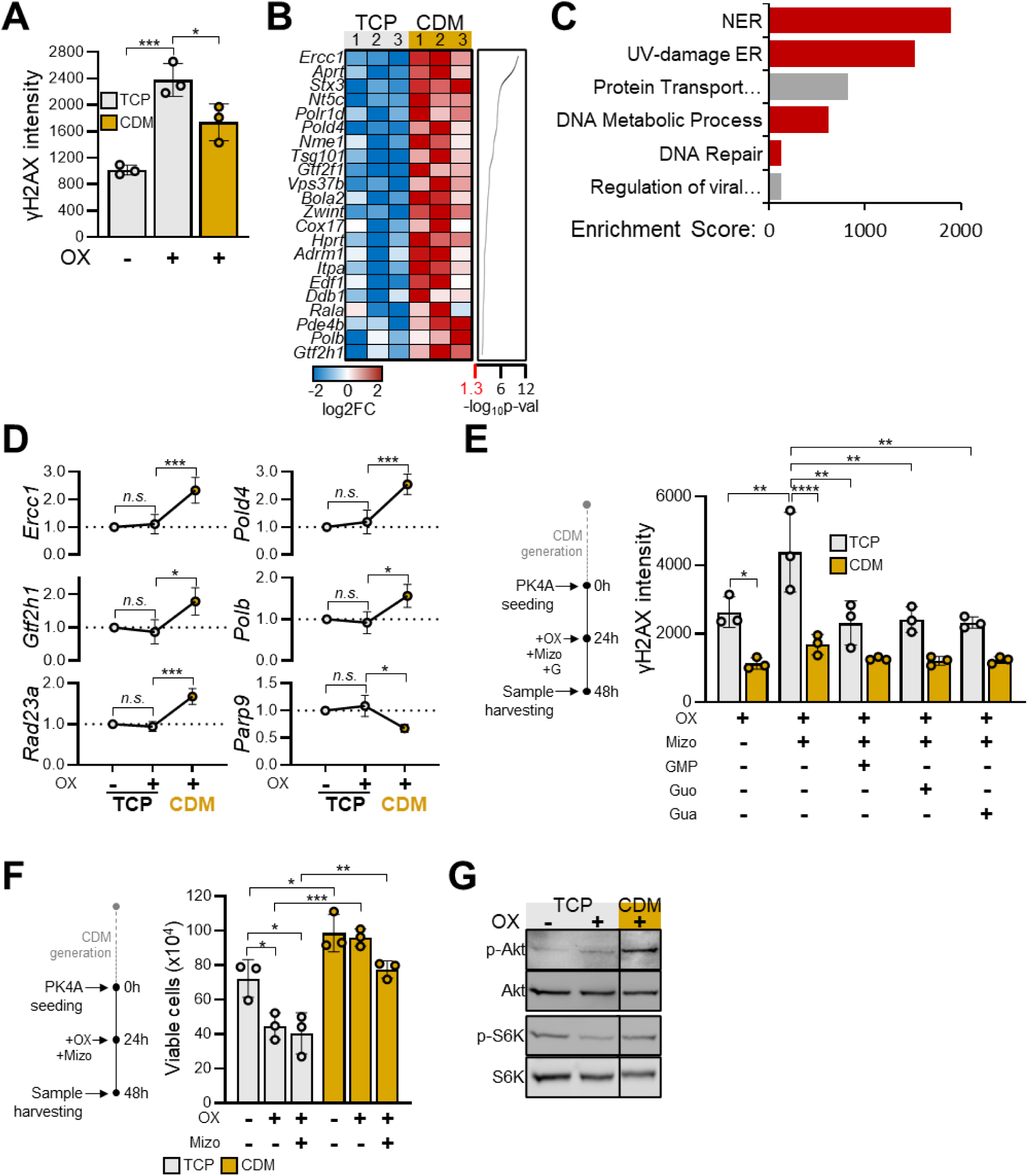
CDM drives OX-induced DNA lesion repair in an IMPDH/GMPS-dependent manner. **(A)** Mean ±SD of γH2AX intensity in PK4A cells grown for 48h on TCP or on CDM in HPLM in the presence of 0.5μM of OX added where indicated concurrently with cell seeding. Ordinary one-way ANOVA, with Tukey’s multiple comparisons test, with a single pooled variance. *** p≤0.001; * p≤0.05. **(B)** Heatmap representation of significantly deregulated DNA repair genes in PK4A cells grown on CDM or on TCP ranked by their –log_10_p value shown on the right. **(C)** Bar graph representation of the pathway enrichment score determined by enrichment analysis using the genes shown in A. Relevant pathways are shown in red. See Table S8 for full list of GO Terms and names. **(D)** mRNA expression levels of selected nucleotide excision repair (NER) genes in PK4A cells grown on CDM compared to TCP in the presence of 0.5μM of oxaliplatin added upon cell seeding. Data are demonstrated as mean fold change ±SD relative to TCP (indicated by the dotted line: TCP+OX vs TCP; CDM+OX vs TCP+OX) from three independent experiments. Ordinary one-way ANOVA. *** p≤0.001; ** p≤0.01; * p≤0.05; n.s., not significant. **(E)** Experimental design (left) and bar graph representation (right) of the γH2AX intensity determined by flow cytometry in PK4A cells grown on TCP or on CDM in HPLM under the simultaneous influence of mizoribine (2.5μM) and oxaliplatin (2μΜ) for 24h, after a 24h drug-free incubation. GMP, Guo, or Gua were individually added as an attempt to rescue the basal phenotype. Data are presented as mean ±SD of three independent experiments. Ordinary two-way ANOVA, with Tukey’s multiple comparisons test, with individual variances computed for each comparison. **** p≤0.0001; *** p≤0.001; ** p≤0.01; * p≤0.05. Not significant comparisons are omitted for clarity. **(F)** Experimental design (left) and bar graph representation of the number of PK4A cells grown on TCP or on CDM in HPLM under the simultaneous influence of mizoribine (2.5μM) and oxaliplatin (2μΜ) for 24h, after a 24h drug-free incubation. Data are presented as mean ±SD of three independent experiments. Ordinary two-way ANOVA, with Tukey’s multiple comparisons test, with individual variances computed for each comparison. *** p≤0.001; ** p≤0.01; * p≤0.05; n.s., not significant. **(G)** Western analysis of the phosphorylation levels of p70 s6K and AKT in cells grown on TCP or CDM for 48h in HPLM with addition of 0.5μM of OX at the time of cell seeding.

To address the association between Impdh/Gmps activity (and the subsequent guanylate production) with the capacity of PDAC cells to repair OX-induced DNA lesions, we measured γH2AX intensity in PK4A cells cultivated on TCP or on CDM in the presence of both OX and mizoribine over the course of 48h. Simultaneous addition of the drugs upon PK4A seeding resulted in a similar and dramatic increase in γH2AX intensity 24h later (Fig. S3A), highlighting the crucial role of a functional Impdh/Gmps duo and of guanylate pools for inherent DNA repair in these cells. To evaluate whether the CDM-driven accumulation of guanylates can alleviate OX-induced DNA damage, PK4A cells were cultivated overnight on TCP or on CDM, and then simultaneously treated with OX and mizoribine (Figure 4E). γH2AX intensity was significantly increased in cells grown on TCP upon treatment with mizoribine/OX compared to OX alone, and the addition of any guanine-containing compound fully restored basal γH2AX levels (Figure 4E), underlining the prominent role of guanylates in DNA repair. In contrast, γH2AX intensity was considerably lower in OX-treated PK4A cells cultivated on CDM compared to TCP irrespective of the presence of mizoribine or the exogenous addition of guanylates (Fig. 4E). Consistently, the number of viable PK4A cells grown on CDM was unaffected by the combined treatment with mizoribine and OX, contrary to those cultivated on TCP which displayed an almost 50% decreased proliferative capacity after OX alone or combined to mizoribine (Fig. 4F). Finally, despite the presence of OX, the phosphorylation levels of p70 S6 kinase, and of Akt were increased in cells grown on CDM compared to TCP (Fig. 4G), suggesting active proliferative signaling. Altogether, these results demonstrate that the accumulation of guanylates in PK4A cells cultivated on CDM limits OX-induced DNA damage, counteracts the mizoribine-mediated inhibition of GMP production, and supports their proliferative potential.

### Guanylate metabolism couples ECM remodeling with DNA repair in PDAC

To address the influence of desmoplasia on purine metabolism and DNA repair at early/preneoplastic, intermediate, and late stages of tumor development, we performed differential gene expression analysis on the entire PDAC from 4-, 6-, and 9-week-old KIC mice compared to the pancreases from age-matched healthy littermates (Fig. 5A). We found that a considerable number of matrisomal genes were significantly deregulated in KIC pancreases compared to healthy ones as early as 4 weeks of age (91 genes up, 4 down, Fig. 5B, Table S9), suggesting that the transformation of the pancreatic tissue occurs before the appearance of anatomopathological structures associated with tumoral features (*45*). At intermediate-stage tumors (6 weeks) we observed a similar number of significantly deregulated genes (98 up, 2 down) as at 4 weeks. In advanced disease stage (9 weeks), however, the number of differentially expressed genes in PDAC compared to healthy pancreas, and the amplitude of deregulation (shown by the range of the log fold change – logFC) increased considerably with 140 up– and 16 down-regulated matrisomal genes (Fig. 5B, Table S9), in line with the prominent ECM deposition and desmoplastic reaction at that stage (*17*). A similar transcriptomic deregulation was observed for purine metabolism genes, with the total number of differentially expressed genes increasing from 20 (16 up, 4 down) in preneoplastic pancreas to 61 (43 up, 18 down) in advanced-stage PDAC (Fig. 5C, Table S10). Finally, the number of significantly deregulated DNA repair genes was highly increased in late stage PDAC compared to intermediate tumor and preneoplastic tissue with a total of 40 genes being up– and 18 genes downregulated (Fig. 5D, Table S11). Gene set enrichment analysis using the deferentially expressed DNA repair genes in advanced PDAC revealed NER among the most transcriptionally deregulated processes (Fig. 5E, Table S12), involving genes like *Ddb1*, *Ercc1*, and *Pold4* that were also modulated by the desmoplasia-mimicking CDM (Fig. 4B, D). These data further support our hypothesis that a desmoplastic environment may trigger alterations in purine metabolism that result in nucleotide imbalance which in turn may tune tumorigenic properties in PDAC cells. Finally, gene expression analysis on 938 primary PDAC patient samples showed a strong correlation between the expression of *IMPDH1*, *IMPDH2*, and *GMPS* and that of DNA repair genes (Fig. 5G). These results suggest that concomitantly with ECM production in the PDAC TME, tumor cells adapt purine metabolism and enhance NER to promote DNA repair, thus supporting PDAC tumorigenic properties throughout disease progression.

**Fig. 5.**
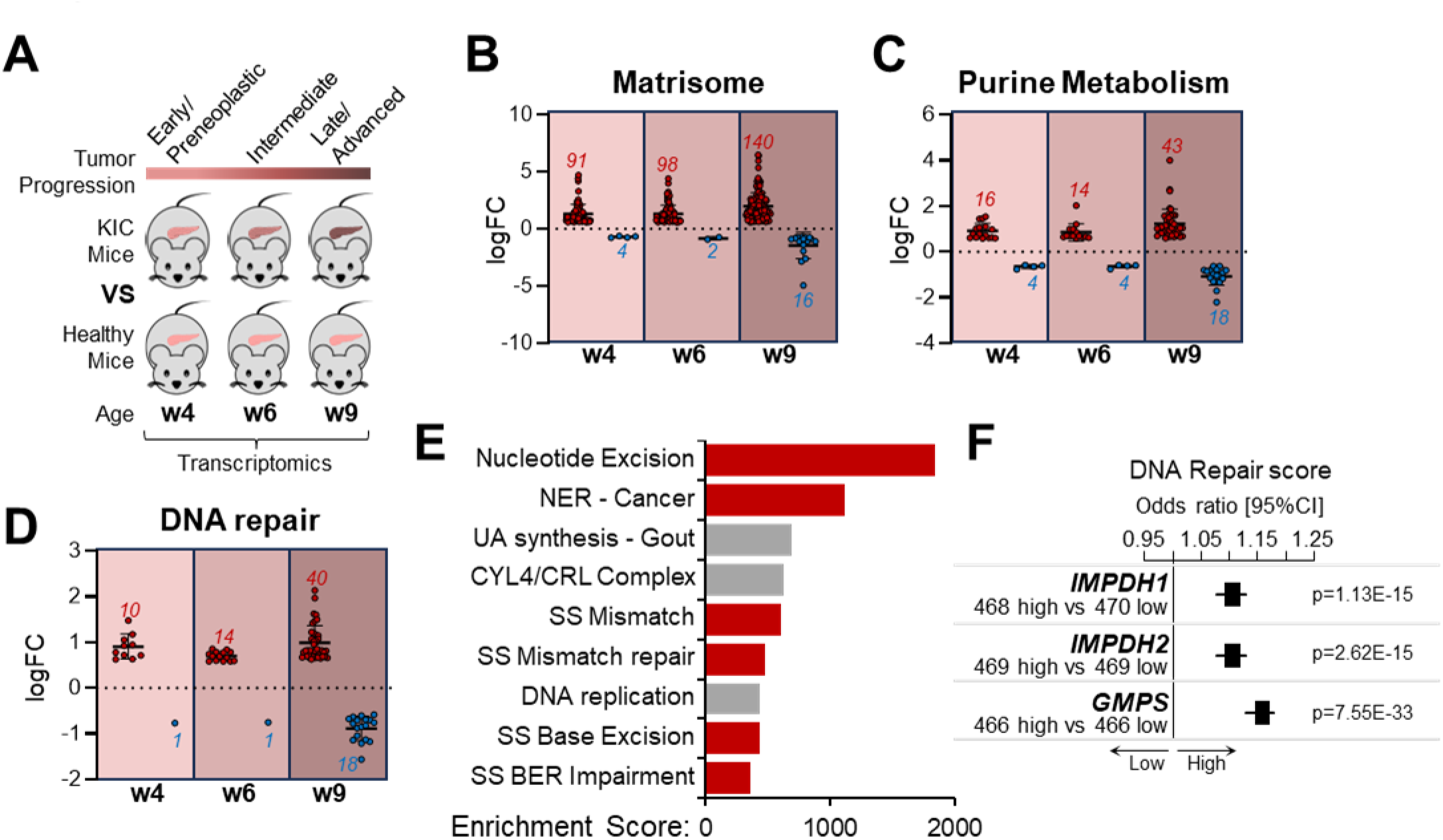
Guanylate metabolism couples ECM remodeling with DNA repair in PDAC. **(A)** Schematic representation of the experimental plan followed to assess the transcriptomic alterations that take place in KIC mice during PDAC progression (at 4, 6, and weeks of age) compared to age-matched healthy animals. **(B)** Graph showing the logFC of differentially expressed matrisomal genes in KIC vs healthy mice at 4, 6, and 9 weeks of PDAC progression. Each dot represents a differentially expressed gene. Red, upregulated genes; blue, downregulated genes. **(C)** Graph showing the logFC of differentially expressed purine metabolism genes in KIC vs healthy mice at 4, 6, and 9 weeks of PDAC progression. Each dot represents a differentially expressed gene. Red, upregulated genes; blue, downregulated genes. **(D)** Graph showing the logFC of differentially expressed DNA repair genes in KIC vs healthy mice at 4, 6, and 9 weeks of PDAC progression. Each dot represents a differentially expressed gene. Red, upregulated genes; blue, downregulated genes. **(E)** Bar graph representation of the pathway enrichment score determined by enrichment analysis using the genes extracted from w9 shown in (E). Relevant pathways are shown in red. See Table S12 for full list of GO Terms and names. **(F)** Forest plot representation of the correlation between the DNA repair metagene and *IMPDH1*, *IMPDH2*, and *GMPS* expression classes in 938 PDAC patient samples as in Fig. 3E.

## Discussion

In this study, we established a robust and highly customizable *in vitro* model using the ECM produced by PDAC-patient-derived CAF. This approach involves the devitalization of CAF monolayers and the use of unabridged ECM substrates that maintain their 3D structure, fiber topology, and secreted factor bioavailability. This system highlights the role of the ECM composition/structure on cellular responses that may go undetected in experimental settings that overlook the topological features of native ECM (e.g. soluble ECM components, gels, coatings) (*33*). For the generation of the CDM we used immortalized patient-derived CAF that express markers of activated myofibroblasts to simulate the ECM generated by myCAF, the major matrix producers/remodelers in PDAC (*2–5*). A recent study revealed a CAF-subtype-dependent ECM heterogeneity that influences patient prognosis in gastric cancer (*46*). Considering the rich ECM observed in PDAC, the study of CAF-specific ECM with distinct tumorigenic or antitumoral features and their impact on patient survival is a field worthy of further research.

The importance of the ECM in PDAC aggressiveness has previously been emphasized by the chemoprotection the desmoplasia confers to tumor cells by forming a protective shield around them, preventing drug penetrance (*47*, *48*). However, whether and how the ECM biochemically contributes to genotoxic stress resistance remains poorly understood. Tumor cell adhesion to fibronectin, a major ECM component with multiple isoforms in solid tumors (*49*), alleviated radiation-induced genotoxic injury in squamous cancer cells (*50*), but the underlying mechanism is unknown. Collagen, the most abundant component of the non-cellular compartment of the TME has a dual role. On the one hand, PDAC cells enhance their proliferative capacity by feeding the TCA cycle with collagen-derived proline acquired from the TME under nutrient stress (*17*). On the other hand, collagen has been shown to confer chemoresistance to 2D– and 3D-cultured PDAC cells by reducing apoptosis (*51*, *52*) and by enhancing the DNA damage response (*53*). The biomechanical and biochemical action of PDAC ECM in chemoresistance was recently highlighted when PDAC cell lines were less sensitive to FOLFIRINOX treatment when cultivated in PDAC-patient-derived decellularized tissue (*54*). Although the underlying mechanisms er not yet elucidated, studies like this stress the functional role of the ECM as a driver of pro-tumoral processes (e.g. chemoresistance) that in turn contribute to PDAC aggressiveness. Here, we demonstrate that ECM-modulated metabolism blunts drug sensitivity in PDAC. More specifically, we demonstrate that PDAC ECM induces an adaptation in purine metabolism in PDAC cells, that fosters PDAC cell survival despite the presence of OX. This is mediated via two routes: i) a metabolic shift in purine metabolism favoring the GMP-generating arm of the *de novo* purine biosynthetic pathway at the expense of AMP production, and ii) the downregulation of guanylate degradation. Together, these routes result in the accumulation of guanosine and guanine. The treatment of PDAC cells with OX in conjunction with simultaneous or deferred addition of the Impdh/Gmps inhibitor mizoribine revealed a strong dependency of PDAC cells on guanylate production. The sensitization of tumor cells to genotoxic stress under guanylate-limited conditions has so far only been described in glioblastoma. In that setting, glioblastoma cells acquired resistance to DNA-damage-inducing radiotherapy by driving the generation of guanylates that in turn promoted DNA repair mechanisms and proliferative signaling (*39*, *55*). Similar to our study, Impdh-inhibition-mediated depletion of guanylates resulted in pronounced increase of γH2AX in cells treated with ionizing radiation *in vitro,* and in a xenograft GBM model (*39*). In the same study, the authors found a decrease in the survival probability in GBM patients with high expression of *IMPDH1* (*39*). In our study, we demonstrate that mizoribine-mediated guanylate depletion increases the amount of OX-induced DNA damage in PDAC cells. The presence of the CDM enhances the removal of OX-induced DNA lesions by the simultaneous upregulation of NER gene expression and the maintenance of cellular guanylate pools. Combined with the activation of proliferative signaling, removal of OX-induced DNA lesions fosters PDAC cell survival. In accordance with our results, a previous study highlighted the role of guanylates in the activation of Akt and mTORC1 in HeLa cells (*56*). In that study, long term guanylate depletion resulted in a considerable decrease in mTORC1 signaling activation (*56*). Although we did not assess the phosphorylation of Akt or p70-S6K upon mizoribine treatment, we demonstrate a high level of pAkt and p-p70-S6K 48h after seeding of PK4A cells on CDM, when guanylates are abundant.

The practical significance of our study lies in the discovery that an ECM – metabolism – DNA repair axis is involved in PDAC cell resistance to OX. More specifically, our analysis showed that the ECM-induced guanylate accumulation in OX-challenged PDAC cells is essential for their ability to moderate DNA damage. Interestingly, despite its strong cytotoxic effects, a major factor that contributes to the failure of FOLFIRINOX treatment is the acquired resistance of PDAC cells via the upregulation of DNA damage response mechanisms (*57*, *58*). Our *in vitro* results are supported by our *in vivo* transcriptomics which demonstrate a synchronous deregulation of matrisomal, purine metabolism and DNA repair genes as the tumor progresses towards an advanced stage. This was also corroborated by the strong correlation between the expression of *IMPDH1*, *IMPDH2*, and *GMPS* with that of DNA repair genes in PDAC patient samples. A study using the GMP-depleting compound MMA (mycophenolic acid), or its derivative MMF (mycophenolate mofetil), showed a moderate decrease in the size of PDAC patient-derived xenografts (PDX), but no significant anti-tumoral effects were observed when administered in a cohort of 6 PDAC patients (*59*). Although clinically approved for the prevention of rejection in renal transplantation (*60*), mizoribine has not yet been tested in PDAC mouse models or in PDAC clinical trials. We propose that the repurposing of mizoribine as a pre-treatment to deplete PDAC cells from accumulated guanylates and sensitize them prior to the administration of platinum-based chemotherapy may be an appealing approach worthy of further investigation. Given the recently proposed use of purine metabolites as markers of disease progression for certain cancers (*61*), purine, and most specifically, guanylate metabolism and its modulation by desmoplasia present a major scientific interest with a potentially high clinical impact.

## Materials and Methods

### Materials and Reagents

All materials and reagents were from Sigma-Merck (St Louis, MO) and Euromedex (France) unless otherwise specified.

### Cell culture, Live Cell Imaging, Proliferation and Cell counting

Immortalized PDAC-patient-derived CAFs (CAFi) were generated as described before (*62*) and cultivated in Dulbecco’s Modified Eagle Medium F12 (DMEM F12) (Thermo Fisher Scientific, Waltham, MA), supplemented with 10% fetal bovine serum (Biosera, France), 100units/ml of penicillin, 100μg/ml of streptomycin, and 0.25μg/ml of amphotericin B (Thermo Fisher Scientific). CAF-derived matrices (CDM) were generated as described before (*63*). Briefly, 72000 cells/cm^2^ were seeded in tissue culture treated vessels and cultivated in culture medium supplemented with 50μg/ml of L-ascorbic acid. Culture medium was replaced every 2 days for a total of 8 days. Matrices were subsequently devitalized (decellularized) with 20mM NH4OH, 0.1% Triton X-100 in PBS, residual DNA was digested with 15μg/ml of DNAse I (Roche, Switzerland) in PBS Ca^2+^Mg^2+^ 1mM, and they were subsequently equilibrated in tumor cell medium (see below). Matrix homogenization was performed using a cell scraper to detach the devitalized, DNA-free matrix which was subsequently passed through a 31G insulin syringe. For CAF proliferation curves, CAFi 1 and CAFi 2 were seeded at an initial density of 50000 cells/well in 24-well plates and enumerated with a Vi-CELL XR Cell Viability Analyzer (Beckman Coulter, Brea, CA) every day for 5 days.

Two batches of tumor cells (hereafter PK4A) were isolated from two male animals of a Pdx-Cre;LSL-Kras^G12D/+^;Ink4a/Arf^flox/flox^ (KIC) mouse model as described before (*10*). Cells were routinely cultivated in glutamine-free Dulbecco’s Modified Eagle Medium (DMEM) (Thermo Fisher Scientific), supplemented with 10% fetal bovine serum, 2mM of glutamine (Thermo Fisher Scientific), 100units/ml of penicillin, 100μg/ml of streptomycin, and 0.25μg/ml of amphotericin B.

All cells were routinely tested for mycoplasma contamination with the MycoAlert® Mycoplasma Detection Kit (Lonza, Switzerland) according to the manufacturer’s instructions.

Prior to all experiments, PK4A cells were cultivated in Human Plasma-Like Medium (HPLM, Thermo Fisher Scientific), supplemented with 10% fetal bovine serum, 100units/ml of penicillin, 100μg/ml of streptomycin, and 0.25μg/ml of amphotericin B, for 2 to 5 days. Then, cells were seeded on CDM-coated or CDM-free 12-well plates (10^5^ cells/well) in HPLM supplemented with 10% fetal bovine serum, 100units/ml of penicillin, 100μg/ml of streptomycin, and 0.25μg/ml of amphotericin B for 48h, unless otherwise stated.

Live cell imaging and video microscopy were performed with an Incucyte® S3 Live-Cell Imaging and Analysis System (Sartorius AG, Germany), while endpoint cell counting was performed with Vi-CELL XR Cell Viability Analyzer.

### Collagen-based gel contraction assay

Collagen contraction assay was performed as described before (*64*) with slight modifications. Briefly, CAFi 1 and CAFi 2 were seeded (2.5X10^5^ cells/lattice) in a mixture of cell culture medium, 1 mg/ml rat tail collagen I (CORNING, Bedford, MA), and the appropriate volume of NaOH 2.5N. Mixtures were cast into 12-well plates, and CAFi culture medium was added on top of the lattices 30 min later. Lattices were detached from the well walls with a sterile spatula and were incubated for a total of 5 days. Plates were scanned every 24h and the lattice surface was quantified. Their contractile capacity was assessed by measuring the reduction of the area occupied by the gels.

### RNA extraction for RNAseq and transcriptomics analysis from in vitro cultures

For the transcriptomic analysis, PK4A cells were cultivated on CDM-coated or non-coated 12-well plates (10^5^ cells/well) in HPLM for 48h. RNA from three independent experiments was subsequently isolated using a RNeasy Plus Kit (Qiagen, Netherlands) and the RNA Integrity Number was determined with an RNA analysis kit (Agilent, Santa Clara, CA). mRNA library preparation was performed with 50ng of RNA per sample using an Illumina Stranded mRNA Prep (Illumina, San Diego, CA) following the manufacturer’s instructions. Samples were sequenced on a NovaSeq 6000 Sequencing System (Illumina) using a S1 Reagent Kit v1.5 (200 cycles) corresponding to 2×30 million reads per sample after demultiplexing. For data analysis, we applied a pipeline developed by the Cibi (CRCM Integrative Bioinformatics) platform. A FASTQ data pre-processing tool was used (Fastp version 0.23.2) for first nucleotide elimination (cutoff quality > 21) and adapter removal. Reads were then mapped to the mm10 reference genome using Subread v1.6.4 with an average mapping score >90% determined with the featureCounts function. Differential analysis was performed with DESeq2 v1.32.0. Only transcripts with more than 20 counts within the same condition were kept. Gene Set Enrichment Analysis (GSEA) was performed using clusterProfiler v4.0.5 and results were visualized using Ridgeplot and DotPlot. Heatmaps were generated with NMF v0.24.0. Other graphs were created with ggplot2 v3.3.5.

### Murine PDAC progression transcriptome analysis

Expression data from pancreases from 4-week-old KIC mice (n = 3 PDAC vs n = 3 control/healthy pancreas) were obtained and analyzed as described before (*45*). For expression data in 6-week-old and 9-week-old KIC mice we used the previously established datasets GSE127891 and GSE61412 respectively. Significantly deregulated genes were extracted from datasets by applying a fold change cutoff of 1.5 and a q value ≤ 0.5 in PDAC vs control in each time point (4, 6, 9 weeks).

### Semi-targeted metabolomics in cell lysates

For metabolomic analysis, PK4A cells were cultivated on CDM-coated or uncoated 12-well plates as described in the previous section in three independent experiments in technical duplicates: one for metabolite detection, the second to determine the cell number per condition and for subsequent normalization. Metabolites were extracted with a 5:3:1 solution of methanol, acetonitrile, and water by adjusting the volume to the cell number (1ml per 10^7^cells). The samples were vigorously mixed for 5 minutes at 4oC, then centrifuged at 16000g for 15 minutes at 4°C. The supernatants were subsequently analyzed by LC/MS with a QExactive Plus Orbitrap mass spectrometer equipped with an Ion Max source and a HESI II probe coupled to a Dionex UltiMate 3000 uHPLC system (Thermo Fisher Scientific). External mass calibration was performed using a standard calibration mixture every seven days, as recommended by the manufacturer. The 5µl samples were injected onto a ZIC-pHILIC column (150 mm × 2.1 mm; i.d. 5 µm) with a guard column (20 mm × 2.1 mm; i.d. 5 µm) (Millipore) for LC separation. Buffer A was 20 mM ammonium carbonate, 0.1% ammonium hydroxide (pH 9.2), and buffer B was acetonitrile. The chromatographic gradient was run at a flow rate of 0.2 µl min−1 as follows: 0–20 min, linear gradient from 80% to 20% of buffer B; 20–20.5 min, linear gradient from 20% to 80% of buffer B; 20.5–28 min, 80% buffer B. The mass spectrometer was operated in full scan, polarity switching mode with the spray voltage set to 2.5 kV and the heated capillary held at 320 °C. The sheath gas flow was set to 20 units, the auxiliary gas flow to 5 units and the sweep gas flow to 0 units. The metabolites were detected across a mass range of 75–1000 m/z at a resolution of 35000 (at 200 m/z) with the automatic gain control target at 106 and the maximum injection time at 250ms. Lock masses were used to ensure mass accuracy below 5 ppm. Data were acquired with Xcalibur Data Acquisition and Intepretation Software (Thermo Fisher Scientific). The peak areas of metabolites were determined using TraceFinder Software (Thermo Fisher Scientific), identified by the exact mass of each singly charged ion and by the known retention time on the HPLC column.

### Metabolomics and Joint Pathway Analysis

For metabolomics analysis, the peak areas previously identified were uploaded in csv file format in MetaboAnalyst 5.0 in two groups (TCP and CDM). Values were normalized by sum and range scaled. Univariate analysis including fold change analysis and t-tests, were performed within the software.

Joint Pathway Analysis was performed with MetaboAnalyst 5.0 using the previously generated transcriptomics and metabolomics datasets. For joint pathway analysis, metabolites with FC≥1.5 and p value≤0.05, and genes with p value≤0.05 (0.1 for PK4A batch 2) in CDM vs TCP were selected. The integration method was performed with the application of a hypergeometric test for enrichment analysis using the KEGG database on both queries.

### Targeted metabolic flux analysis

For metabolic flux analysis of glutamine and hypoxanthine, a “custom-made” HPLM (cm-HPLM) was prepared by combining the method published by Cantor et al (*37*) with the official HPLM composition from Thermo Fisher Scientific. PK4A cells were seeded on CDM-coated or non-coated 10cm dishes (3×10^6^ cells/dish) in HPLM supplemented with 10% fetal bovine serum, 100units/ml of penicillin, 100μg/ml of streptomycin, and 0.25μg/ml of amphotericin B. Oxaliplatin (Hospira, Lake Forest, IL) was added to the cells at the time of seeding at a final concentration of 0.32μM, to condition the cells without significantly reducing their viability. Cells were then incubated overnight at 37°C in a humidified incubator supplemented with 5% CO2. For amido-^15^N-glutamine flux studies, cells were washed once with glutamine-free cm-HPLM and then incubated for 3 hours in cm-HPLM supplemented with 10% dialyzed fetal bovine serum (Thermo Fisher Scientific), 100units/ml of penicillin, 100μg/ml of streptomycin, and 0.25μg/ml of amphotericin B, containing 4mM of amido-^15^N-glutamine (Cambridge Isotope Laboratories, Tewksbury, MA) and 0.5μM of oxaliplatin. For ^13^C-hypoxanthine flux studies, cells were washed once with hypoxanthine-free cm-HPLM and then incubated for 3 hours in cm-HPLM supplemented with 10% fetal bovine serum, 100units/ml of penicillin, 100μg/ml of streptomycin, and 0.25μg/ml of amphotericin B, containing 100μM of ^13^C-hypoxanthine (Cambridge Isotope Laboratories), and 0.5μM of oxaliplatin. The medium was then removed, cells were incubated with 4ml of 80% methanol for 20 minutes at –80°C and were subsequently scraped off the culture dishes and transferred into 15ml conical tubes. Insoluble material was pelleted by centrifugation at 3000g for 5 minutes at –4°C, followed by two subsequent centrifugations of the insoluble pellet with an additional 0.5ml of 80% methanol and centrifugation at 21000g for 5 minutes at 4°C. The pooled supernatants were dried under nitrogen gas using an N-EVAP (Organomation Associates, Berlin, MA). Dried pellets were re-suspended in 20µL HPLC-grade water for mass spectrometry and 5µL were injected using a 5500 QTRAP hybrid triple quadrupole mass spectrometer (ABI/SCIEX, Framingham, MA) coupled to a Prominence UFLC HPLC system (Shimadzu Corporation, Japan) with Amide XBridge HILIC chromatography (Waters Corporation, Milford, MA) via selected reaction monitoring (SRM) (*65*) and polarity switching between positive and negative modes. The parameters for instrumental and software analysis were as described before (*66*). Peak areas from the total ion current for each metabolite SRM transition were integrated using MultiQuant v2.0 software (SCIEX, Framingham, MA). Custom SRMs were created for expected 15N or 13C incorporation in various forms for targeted LC-MS/MS.

### RNA extraction and qPCR amplification

Cells were seeded in 12-well plates as described above for 48h, then lysed and RNA was isolated with Trizol (Thermo Fisher Scientific) according to the manufacturer’s guidelines. Reconstituted RNA was quantified with a Fluostar Omega spectrophotometer (BMG Labtech, Germany). cDNA was generated with a GoScript cDNA generation kit (Promega, France) using random primers starting from 1μg of RNA. Each qPCR reaction was performed with 5ng of cDNA in triplicates using a GoTaq SYBR Green qPCR Mix (Promega) in a AriaMX qPCR System (Agilent) using the gene specific primers listed in Table S13. Raw fluorescence values were baseline-corrected with a passive reference dye (ROX), normalized with Tbp using the Pfaffle method for primer efficiency correction (*67*), and represented as fold change relative to the indicated control.

### Whole cell extracts and western blot

Cells were seeded in 12-well plates for 48h as described above, then rinsed with ice-cold PBS and lysed with 3x Laemmli lysis buffer (6% SDS, 30% glycerol, 187.5mM Tris-HCl pH 6.8). Lysates were subsequently sonicated with a Pulse 150 ultrasonic homogenizer (Benchmark Scientific, Sayreville, NJ), heated at 95°C for 5 minutes and quantified with a BCA Quantification Assay Kit (Thermo Fisher Scientific). Polyacrylamide gels were cast and 20-40μg of whole cell lysates were loaded per well for electrophoresis and transferred on PVDF membranes (Merck Millipore, Burlington, MA). Membranes were subsequently blocked with a 5% skim milk in TBS solution, followed by primary antibody incubation in 3% BSA in TBS-Tween 0.1% (see primary antibody list in Table S14). A secondary antibody coupled with horse radish peroxidase (Southern Biotech, Birmingham, AL) was used against the primary antibody isotype, followed by an enhanced chemiluminescence reaction (Merck Millipore). Protein bands were visualized in a PXi Multi-Application Gel Imagin System (Syngene, UK) and band densitometry analysis was performed with FIJI (*68*).

### Immunofluorescence and Imaging

For immunofluorescence-based protein detection and microscopy, PK4A cells were seeded on glass coverslips (Paul Marienfeld, Germany) as described above. Cells were then incubated with a fixing solution (4% paraformaldehyde, 3% sucrose in PBS) at 37oC for 20 minutes, washed with PBS, permeabilized with 0.2% Triton X-100 in PBS for 30 minutes, blocked with 3% BSA in PBS, incubated with the respective primary antibodies followed by secondary antibodies coupled to ALEXA fluorophores (see Table 14). Cells were counterstained with Hoechst 33342 (Thermo Fisher Scientific) and coverslips were finally mounted on glass slides with Prolong Gold anti-fade reagent (Thermo Fisher Scientific). Cell visualization and image acquisition were performed with a Zeiss Axio Imager microscopy system (Zeiss, Germany). Images were treated and analyzed with FIJI.

### Cytometry

For CAF phenotyping, cells were cultivated as described above. Cells were detached using StemPro accutase cell dissociation reagent (Thermo Fisher Scientific) and washed with PBS. Samples were resuspended in flow cytometry buffer (0.5% BSA, 2mM EDTA in PBS) and stained with extracellular fluorochrome-coupled antibodies (see Table 14) for 20 minutes at 4°C. Samples were then washed with PBS and one drop of viability marker Sytox green (Thermo Fisher Scientific) was added to each sample 5 minutes prior to analysis. Single-stained samples for compensation were made by staining UltraComp eBeads™ Compensation Beads (Thermo Fisher Scientific) with the same protocol as above. Samples were analyzed using a BD LSR-Fortessa-X20 Cell Analyzer (Becton Dickinson Biosciences, Franklin Lakes, NJ) and data were analyzed using FlowJo software (Becton Dickinson Biosciences).

For γH2AX staining, PK4A cells were seeded in 12-well plates as described before, with or without oxaliplatin and/or mizoribine and purine compounds (all from Sigma-Merck) at cell seeding unless otherwise specified for the indicated duration. Cells were then trypsinized, fixed with 3.7% paraformaldehyde in PBS, and incubated with a 1/10 dilution of a BD Pharmingen™ Alexa Fluor® 647 Mouse anti-H2AX, pS139 (Becton Dickinson Biosciences). Cytometry was performed in a BD LSRFortessa-X20 Cell Analyzer, data were analyzed with FlowJo v.10.10 and expressed as the mean cellular fluorescence intensity.

### PDAC Patient Dataset Analysis

Clinicopathological and gene expression data were collected from 17 publicly available mRNA expression datasets (Table S15) comprising a total of 938 primary PDAC samples. Patients’ informed consent to participate was obtained by the authors of each study, while ethical considerations were overseen by the respective institutional review board. Data pre-analytic processing was performed as described before (*69*). Batch effect correction on genes of interest was performed using z-score normalization with the mean and the standard deviation measured on primary cancer samples of each dataset. Tumor expression of purine genes was measured as discrete values after comparison with median expression in the 938 samples: high expression was defined by value > median and low expression by value ≤ median. Matrisomal and DNA repair genes (Table S16, Table S17) were similarly normalized and both metagenes were computed as the mean expression of all standardized genes from the respective list. All statistical tests were two-sided at the 5% level of significance. Statistical analysis was done using the survival package (version 3.5-5) in the R software (version 4.1.2; http://www.cran.r-project.org/).

### Statistics

Statistical analyses were performed with GraphPad Prism (GraphPad Software, Boston, MA). Appropriate statistical tests tailored to the specific conditions/analyses and the respective number of replicates are described in detail in the figure legends. Experimental versus control group differences were considered significant when p ≤ 0.05. Graphs and charts were created with Graph Pad Prism, or with Excel 365 (Microsoft Corporation, Redmond, WA). For the fold change analysis in the murine PDAC progression dataset, statistical differences were assessed with pairwise t test in log-transformed fold change data coupled with Bonferroni correction to adjust p values. (*70*)

## Acknowledgments

We wish to thank Dr Laura Brau for her technical assistance, as well as INSERM Cell Culture Platform (PCC, Marseille, France) and the CRCM bioinformatic platform (Cibi, Marseille, France) for their services. We extend our thanks to the BIDMC-Harvard Mass Spectrometry Facility and Asara Laboratory for their help with the steady-state metabolic tracing.

## Author contributions

Conceptualization: GE, SV

Data curation: GE, EL, PK, PB, PF, FB, GB, FG

Formal Analysis: GE, EL, IBS, PK, PB, PF, FB, FG

Funding Acquisition: GE, CL, RT, SV

Investigation: GE, CC, IN, SST, HSC, ZH, FG, SV

Methodology: GE, SV

Project Administration: GE, SV

Software: EL, PK, PB, GB

Resources: FB, RT, GB, SV

Supervision: RT, SV

Visualization: GE, EL, PK, SST, ZH, PF, FG, SV

Writing—original draft: GE, SV

Writing—review & editing: GE, FB, PF, RT, FG, SV

## Data and materials availability

The DNA microarray raw data from 4-week-old mice, and the RNA sequencing datasets from the two PK4A cell batches grown on CDM and on TCP have been deposited in Gene Expression Omnibus under GSE284155and GSE284085 respectively. Other data and materials can be made available upon request.

