## Supplemental Information for "The extracellular matrix drives guanylate production and protects pancreatic cancer cells from oxaliplatin-induced DNA damage"

Georgios Efthymiou *et al.*

*Georgios Efthymiou, Sophie Vasseur.

**This PDF file includes:**

Supplementary Text
Figs. S1 to S3
Tables S1 to S17

**Supplementary Text**

Integration of RNAseq and semi-targeted metabolomics datasets in PK4A cells cultivated on CDM vs TCP.

For the first batch of PK4A cells, differentially expressed genes on CDM vs TCP with adjusted p value ≤ 0.05, and metabolites with fold change ≥ 1.5 and a p value ≤ 0.05 on CDM vs TCP were used and Joint Pathway Analysis was performed with MetaboAnalyst 5.0 based on the mouse KEGG database. The results of the enrichment analysis are shown in Supplementary Table S1. Figure 1B was obtained after filtering out i) pathways with FDR ≤ 0.05, and ii) pathways with impact ≤ 1. For the second batch of PK4A cells, differentially expressed genes on CDM vs TCP with adjusted p value ≤ 0.1, and metabolites with fold change ≥ 1.5 and a p value ≤ 0.1 on CDM vs TCP were used and Joint Pathway Analysis was performed with MetaboAnalyst 5.0 based on the mouse KEGG database. The results of the enrichment analysis are shown in Supplementary Table S2. Figure S1F was obtained after filtering out i) pathways with FDR ≤ 0.05, and ii) pathways with impact ≤ 1.

Heatmaps on Purine and Pyrimidine Compounds from cells grown on CDM vs TCP.

Normalized values of purine and pyrimidine metabolism compounds were obtained from metabolomics data obtained from two independent experiments on two murine PDAC cell populations as described in Materials and Methods. The mean was obtained for each metabolite and Student's t test was used to calculate p values in CDM vs TCP, which was subsequently converted to -log10(p value). The heatmaps (Fig. 1C and Fig. S1I) were generated with the values shown on Table S3 and Table S6 respectively.

Heatmaps on Purine and Pyrimidine Gene Expression from cells grown on CDM vs TCP.

The heatmaps on purine and pyrimidine metabolism gene expression between CDM vs TCP (Fig. 1E and Fig. S1H) were generated using the values from the raw reads from the transcriptomics dataset on a batch of PK4A cells grown on CDM and TCP. The gene expression values, alongside the respective fold change and p value for purine and pyrimidine metabolism gene expression are shown on Table S4 and Table S5 respectively.

DNA Repair gene expression in PK4A cells cultivated on CDM vs TCP in the presence of oxaliplatin (OX).

The expression of the DNA repair genes in OX-treated PK4A cells grown on TCP is represented as fold change relative to PK4A cells grown on TCP in the absence of OX. Similarly, the expression of these genes in OX-treated PK4A cells grown on CDM is represented as fold change relative to OX-treated PK4A cells grown on TCP. Statistics were performed separately within these comparison groups, and the FC=1 in the CDM+OX vs TCP+OX comparison was removed from Fig. 4D to facilitate the visualization of the results.

Gene set enrichment analysis

For gene set enrichment analysis (GSEA), we used the online GSEA tool Enrichr (70). For GSEA on DNA repair gene expression in PK4A cells cultivated on CDM vs TCP, we uploaded the genes shown in Fig. 4B and we subsequently selected the "GO Biological Process 2023" superfamily composed of the elements shown on Table S8. Fig.4C was generated after filtering out Terms with an adjusted p value > 0.01. For GSEA on DNA repair gene expression in 9-week-old KIC mice relative to age-matched littermates, we uploaded the genes shown in Fig. 5D shown in Table S11, and we subsequently selected the "Elsevier Pathway Collection" superfamily with the GO Terms shown on Table S12. We kept Terms with an adjusted p value ≤ 0.01 to generate Fig. 5E.

#### Supplementary Figure 1

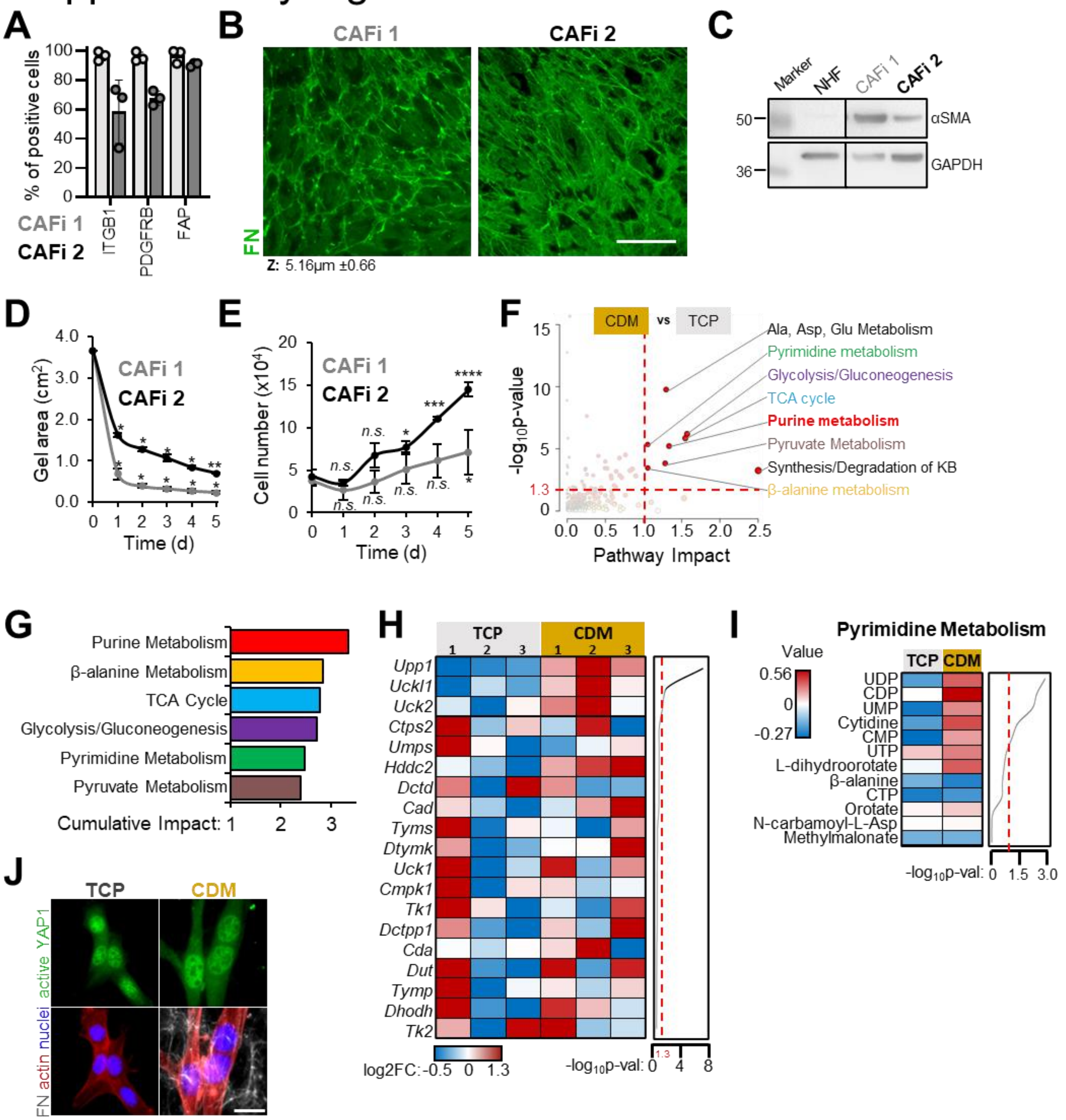


**Fig. S1. CAF-derived ECM drives purine imbalance in PDAC cells. (A)** Cytometry-based phenotypic analysis of two PDAC-patient-derived cancer associated fibroblast (CAF) populations. ITGB1, integrin β1; PDGFRB, platelet-derived growth factor receptor B; FAP, fibroblast activation protein alpha; Mean ±SD from three independent experiments. **(B)** Immunofluorescence of fibronectin (FN) fibrils deposited by CAFi 1 and 2 after 8 days in culture. The culture medium was replaced every two days and supplemented with 50µg/ml of L-ascorbic acid. The thickness of the FN matrix is shown beneath. Scale bar, 100µm. **(C)** Western analysis on whole cell lysates from the CAFi populations shown in (A) and comparison with normal human fibroblasts (NHF) for αSMA. GAPDH was used as loading control. **(D)** Collagen-based gel contraction assay in CAFi 1 and 2. The contractile capacity was assessed by measuring the reduction of the area occupied by the collagen lattices inside the wells of 12-well culture vessels. Data are represented as mean ±SD from two independent experiments. Statistical comparisons are displayed between each time point and t=0. Repeated measures two-way ANOVA with the Geisser-Greenhouse correction and Dunnett's multiple comparisons test, with individual variances computed for each comparison. ** $p \leq 0.01$; * $p \leq 0.05$. **(E)** Proliferation curves of CAFi 1 and 2. Data are represented as mean ±SD from two independent experiments. Statistical comparisons are displayed between each time point and t=0. Repeated measures two-way ANOVA with Dunnett's multiple comparisons test, with a single pooled variance. **** $p \leq 0.0001$; *** $p \leq 0.001$; * $p \leq 0.05$. n.s., not significant. **(F)** Integration analysis of matched transcriptomics and metabolomics datasets in a second batch of PK4A cells after a 48h culture in HPLM on CDM or on TCP. Each dot represents a KEGG metabolic pathway, and the dot size reflects the number of identified features (genes/metabolites) within that pathway. A threshold of 1.3 was applied for $-\log_{10}p$ val (red horizontal dotted line), and 1.0 for pathway impact (red vertical dotted line). **(G)** Bar graph representation of the cumulative impact of the common deregulated metabolic pathways between the batches of PK4A cells tested (Fig.1B and Fig. S1F). **(H)** Heatmap representation of Pyrimidine Metabolism genes identified and extracted from the integration analysis in Fig. 1B. The genes are ranked by their $-\log_{10}p$ value shown on the right and the threshold of significance ($-\log_{10}p \geq 1.3$) is indicated by the dotted line. **(I)** Heatmap representation of Pyrimidine Metabolism Compounds in PK4A cells cultivated on TCP or on CDM for 48h in HPLM. Cumulative data from two independent experiments in two batches of PK4A cells. Compounds are ranked by significance with the $-\log_{10}p$ value shown on the right. The dotted line delineates the threshold of significance ($-\log_{10}p \geq 1.3$). **(J)** Immunofluorescent staining of active YAP1 in PK4A cells after a 24-h culture on TCP or CDM in HPLM. FN was used to visualize the CDM and actin to delimit cell body, while nuclei were counterstained with Hoechst 33342. Scale bar, 25µm.

#### Supplementary Figure 2

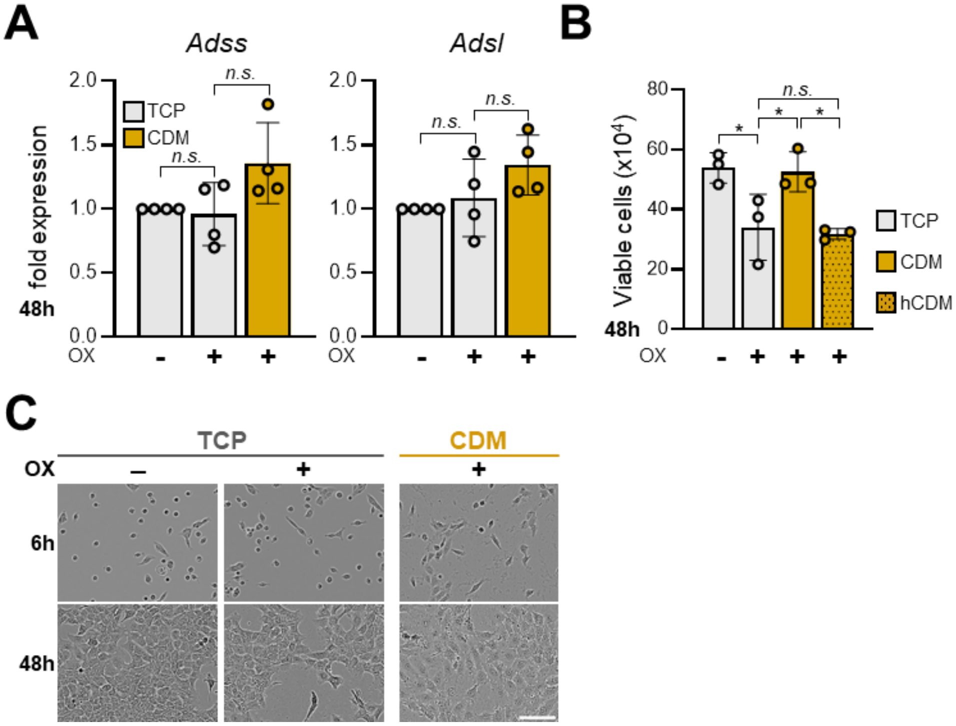


**Fig. S2. CDM drives GMP production and oxaliplatin resistance in PDAC cells. (A)** Expression of *Adss* and *Adsl* mRNA in PK4A cells grown on CDM compared to TCP in HPLM in presence of 0.5μM of OX for 48h. Data are represented as fold change relative to TCP (TCP+OX vs TCP; CDM+OX vs TCP+OX) from four independent experiments. Ordinary one-way ANOVA. n.s., not significant. **(B)** Bar graph displaying the number of a second batch of PK4A cells grown on TCP or on CDM after 48h in HPLM with addition of 0.5μM of OX. hCDM signifies a homogenized CDM preparation presented to the cells concurrently with cell seeding and OX addition. Data are presented as mean ±SD of three independent experiments. Ordinary one-way ANOVA, with Tukey's multiple comparisons test, with a single pooled variance. * p≤0.05; n.s., not significant. **(C)** Phase contrast images of PK4A cells cultivated on TCP or on CDM in HPLM for 6h (top), or 48h (bottom) in the presence of 0.5μM of OX. Scale bar, 50μm.

### Supplementary Figure 3

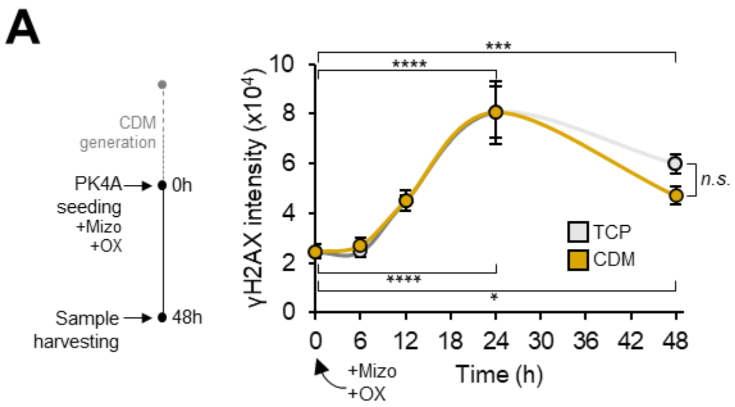


**Fig. S3. Guanylate depletion results in massive DNA damage. (A)** Kinetic of γH2AX intensity in PK4A cells cultivated on TCP or on CDM with simultaneous addition of 2.5μM of mizoribine and 0.5μM of oxaliplatin at the time of cell seeding. Individual time points represent mean ±SD from three independent experiments. Comparisons are between each time point and 0h of the respective condition. Repeated measures two-way ANOVA with the Geisser-Greenhouse correction and Dunnett's multiple comparisons test, with individual variances computed for each comparison. **** p≤0.0001; *** p≤0.001; n.s., not significant.

| Term | FDR | Impact |
|---|---|---|
| Purine metabolism | 3.67E-05 | 2.0147 |
| beta-Alanine metabolism | 0.0092773 | 1.7778 |
| JAK-STAT signaling pathway | 0.11432 | 1.7222 |
| Nicotinate and nicotinamide metabolism | 0.0011759 | 1.5 |
| Pyrimidine metabolism | 0.00050358 | 1.4186 |
| Focal adhesion | 1.83E-08 | 1.4032 |
| Phosphatidylinositol signaling system | 0.4645 | 1.2727 |
| Citrate cycle (TCA cycle) | 0.00057009 | 1.2273 |
| Glycolysis or Gluconeogenesis | 0.0026826 | 1.1538 |
| Pyruvate metabolism | 0.014292 | 1.1132 |
| Circadian rhythm | 0.18562 | 1.05 |
| Alanine, aspartate and glutamate metabolism | 2.20E-05 | 0.90164 |
| Arginine biosynthesis | 0.00048772 | 0.88571 |
| Glutathione metabolism | 0.012984 | 0.82609 |
| Long-term depression | 0.037967 | 0.78261 |
| Pentose phosphate pathway | 0.029943 | 0.77778 |
| Glycerophospholipid metabolism | 0.36758 | 0.7619 |
| Chemokine signaling pathway | 0.05326 | 0.75 |
| ECM-receptor interaction | 9.51E-08 | 0.72917 |
| Fc gamma R-mediated phagocytosis | 0.00039391 | 0.7037 |
| Regulation of actin cytoskeleton | 0.0022654 | 0.6988 |
| Inositol phosphate metabolism | 0.11796 | 0.67059 |
| Glycosphingolipid biosynthesis - lacto and neolacto series | 0.76359 | 0.65789 |
| Glutamatergic synapse | 0.12159 | 0.65714 |
| Pantothenate and CoA biosynthesis | 0.012984 | 0.64286 |
| Phospholipase D signaling pathway | 0.007648 | 0.62319 |
| Fructose and mannose metabolism | 0.11008 | 0.62162 |
| Platelet activation | 9.87E-05 | 0.61628 |
| cGMP-PKG signaling pathway | 0.012984 | 0.60976 |
| Arginine and proline metabolism | 0.00029585 | 0.60684 |
| Synthesis and degradation of ketone bodies | 0.3746 | 0.6 |
| Glycine, serine and threonine metabolism | 0.05667 | 0.59091 |
| Riboflavin metabolism | 0.05326 | 0.58333 |
| Mucin type O-glycan biosynthesis | 0.5459 | 0.58333 |
| Hippo signaling pathway - multiple species | 0.0022654 | 0.5814 |
| Circadian entrainment | 0.20619 | 0.55172 |
| Ether lipid metabolism | 0.99801 | 0.53488 |

| | | |
|---|---|---|
| Relaxin signaling pathway | 0.017166 | 0.52222 |
| Notch signaling pathway | 0.45431 | 0.51852 |
| Bacterial invasion of epithelial cells | 0.0022654 | 0.51163 |
| Vascular smooth muscle contraction | 0.025373 | 0.50704 |
| Leukocyte transendothelial migration | 0.015157 | 0.50549 |
| TGF-beta signaling pathway | 0.00085393 | 0.5 |
| Histidine metabolism | 0.088759 | 0.5 |
| Wnt signaling pathway | 0.1672 | 0.4878 |
| EGFR tyrosine kinase inhibitor resistance | 0.25355 | 0.48485 |
| Longevity regulating pathway | 0.15161 | 0.48148 |
| Fatty acid degradation | 0.39657 | 0.47899 |
| Prolactin signaling pathway | 0.15161 | 0.4697 |
| Glioma | 0.21973 | 0.46575 |
| AGE-RAGE signaling pathway in diabetic complications | 4.44E-05 | 0.46269 |
| Gap junction | 0.040362 | 0.46154 |
| Oxytocin signaling pathway | 0.11432 | 0.46053 |
| Fc epsilon RI signaling pathway | 0.52512 | 0.44 |
| Arachidonic acid metabolism | 1 | 0.4359 |
| Neurotrophin signaling pathway | 0.33843 | 0.42683 |
| Thyroid hormone signaling pathway | 0.23856 | 0.42453 |
| Tight junction | 0.18025 | 0.42222 |
| Hippo signaling pathway | 0.0022654 | 0.42157 |
| Renal cell carcinoma | 0.088759 | 0.42105 |
| Long-term potentiation | 0.47539 | 0.41935 |
| Chronic myeloid leukemia | 0.011025 | 0.41818 |
| Calcium signaling pathway | 0.015446 | 0.41791 |
| PI3K-Akt signaling pathway | 0.00029585 | 0.41584 |
| Longevity regulating pathway - multiple species | 0.3746 | 0.41509 |
| Autophagy - animal | 0.23272 | 0.4 |
| Growth hormone synthesis, secretion and action | 0.42183 | 0.39683 |
| Sphingolipid metabolism | 0.68583 | 0.39344 |
| Retinol metabolism | 1 | 0.38983 |
| Tyrosine metabolism | 0.5459 | 0.3871 |
| Epstein-Barr virus infection | 0.17613 | 0.38621 |
| Metabolism of xenobiotics by cytochrome P450 | 1 | 0.37912 |
| ErbB signaling pathway | 0.45066 | 0.37681 |
| D-Glutamine and D-glutamate metabolism | 0.14991 | 0.375 |
| B cell receptor signaling pathway | 0.61056 | 0.36735 |
| HIF-1 signaling pathway | 0.023362 | 0.36585 |
| Sphingolipid signaling pathway | 0.13069 | 0.36585 |

| | | |
|---|---|---|
| VEGF signaling pathway | 0.3746 | 0.36364 |
| Axon guidance | 0.0022654 | 0.36242 |
| Fluid shear stress and atherosclerosis | 0.12275 | 0.3619 |
| Rap1 signaling pathway | 0.05326 | 0.3617 |
| GnRH signaling pathway | 0.36198 | 0.3617 |
| Choline metabolism in cancer | 0.075347 | 0.35849 |
| Type II diabetes mellitus | 0.45431 | 0.35484 |
| Pathways in cancer | 0.0055333 | 0.33956 |
| Human cytomegalovirus infection | 0.22328 | 0.33742 |
| Pancreatic cancer | 0.12505 | 0.33333 |
| N-Glycan biosynthesis | 0.62809 | 0.32911 |
| Yersinia infection | 0.14222 | 0.32632 |
| Adrenergic signaling in cardiomyocytes | 0.59651 | 0.32394 |
| MAPK signaling pathway | 0.020819 | 0.32258 |
| Morphine addiction | 0.17931 | 0.32 |
| Apelin signaling pathway | 0.4645 | 0.32 |
| Dopaminergic synapse | 0.61957 | 0.31944 |
| NF-kappa B signaling pathway | 0.088759 | 0.31875 |
| Linoleic acid metabolism | 1 | 0.31707 |
| Osteoclast differentiation | 0.10142 | 0.31579 |
| Cholinergic synapse | 0.32491 | 0.31148 |
| Staphylococcus aureus infection | 0.99187 | 0.30769 |
| Proteoglycans in cancer | 0.0015016 | 0.305 |
| Parathyroid hormone synthesis, secretion and action | 0.17496 | 0.30488 |
| Platinum drug resistance | 0.075347 | 0.3 |
| Taste transduction | 0.12159 | 0.3 |
| Breast cancer | 0.49665 | 0.3 |
| Estrogen signaling pathway | 0.33009 | 0.2987 |
| Melanogenesis | 0.1985 | 0.29545 |
| Serotonergic synapse | 0.81849 | 0.29487 |
| Adherens junction | 0.45395 | 0.29333 |
| Phenylalanine, tyrosine and tryptophan biosynthesis | 0.33009 | 0.29268 |
| Apoptosis | 0.31937 | 0.28829 |
| Hepatocellular carcinoma | 0.027238 | 0.28814 |
| Signaling pathways regulating pluripotency of stem cells | 0.020375 | 0.28571 |
| Retrograde endocannabinoid signaling | 0.48128 | 0.28571 |
| Pertussis | 0.094791 | 0.28358 |
| Regulation of lipolysis in adipocytes | 0.084828 | 0.28333 |
| Cysteine and methionine metabolism | 0.096088 | 0.28155 |
| Inflammatory mediator regulation of TRP channels | 0.45431 | 0.2809 |
| Glycosphingolipid biosynthesis - ganglio series | 1 | 0.27586 |

| | | |
|---|---|---|
| mTOR signaling pathway | 0.027662 | 0.27381 |
| PPAR signaling pathway | 0.67231 | 0.27119 |
| Ras signaling pathway | 0.27916 | 0.27083 |
| Acute myeloid leukemia | 0.45395 | 0.27083 |
| Gastric acid secretion | 0.45066 | 0.26923 |
| Human papillomavirus infection | 9.28E-05 | 0.26875 |
| C-type lectin receptor signaling pathway | 0.077183 | 0.26842 |
| GABAergic synapse | 0.15161 | 0.26786 |
| Human T-cell leukemia virus 1 infection | 0.13024 | 0.26154 |
| Measles | 0.3455 | 0.25843 |
| Natural killer cell mediated cytotoxicity | 0.42874 | 0.25773 |
| Necroptosis | 0.3746 | 0.25532 |
| Th1 and Th2 cell differentiation | 0.46022 | 0.25352 |
| Cytokine-cytokine receptor interaction | 0.031484 | 0.25 |
| Endocrine resistance | 0.42874 | 0.24638 |
| Complement and coagulation cascades | 0.053354 | 0.24615 |
| cAMP signaling pathway | 0.038609 | 0.2459 |
| Leishmaniasis | 0.18562 | 0.24528 |
| Melanoma | 0.4645 | 0.24242 |
| Insulin signaling pathway | 0.75818 | 0.24242 |
| Taurine and hypotaurine metabolism | 0.0088821 | 0.24138 |
| Glyoxylate and dicarboxylate metabolism | 0.031518 | 0.23864 |
| Selenocompound metabolism | 0.84649 | 0.2381 |
| Chagas disease (American trypanosomiasis) | 0.016486 | 0.23611 |
| Drug metabolism - other enzymes | 0.65972 | 0.23333 |
| Gastric cancer | 0.29454 | 0.23301 |
| Non-small cell lung cancer | 0.46022 | 0.23214 |
| Colorectal cancer | 0.053354 | 0.22973 |
| Mannose type O-glycan biosynthesis | 0.64515 | 0.22917 |
| Herpes simplex virus 1 infection | 0.77152 | 0.22727 |
| Progesterone-mediated oocyte maturation | 0.6896 | 0.225 |
| Glycerolipid metabolism | 0.90397 | 0.22414 |
| Butanoate metabolism | 0.079378 | 0.21818 |
| Dilated cardiomyopathy (DCM) | 0.040362 | 0.21739 |
| Tuberculosis | 0.33125 | 0.21324 |
| Apoptosis - multiple species | 0.18562 | 0.21212 |
| Lysine degradation | 0.3746 | 0.20779 |
| Aminoacyl-tRNA biosynthesis | 0.00010123 | 0.20619 |
| Synaptic vesicle cycle | 0.0014296 | 0.20588 |
| Insulin resistance | 0.16033 | 0.20588 |
| Fatty acid elongation | 0.44048 | 0.20253 |

| | | |
|---|---|---|
| Basal cell carcinoma | 0.22769 | 0.2 |
| Th17 cell differentiation | 0.31937 | 0.2 |
| Valine, leucine and isoleucine degradation | 0.54224 | 0.19792 |
| Ubiquinone and other terpenoid-quinone biosynthesis | 1 | 0.19767 |
| T cell receptor signaling pathway | 0.62196 | 0.19718 |
| Prostate cancer | 0.47267 | 0.19672 |
| Hepatitis C | 0.74874 | 0.19588 |
| Propanoate metabolism | 0.23856 | 0.19444 |
| Thermogenesis | 0.3499 | 0.19277 |
| Valine, leucine and isoleucine biosynthesis | 0.01324 | 0.19231 |
| Phenylalanine metabolism | 0.25355 | 0.19178 |
| GnRH secretion | 0.17496 | 0.19149 |
| FoxO signaling pathway | 0.070436 | 0.19048 |
| Salmonella infection | 0.029764 | 0.1875 |
| PD-L1 expression and PD-1 checkpoint pathway in cancer | 1 | 0.1875 |
| Adipocytokine signaling pathway | 0.9137 | 0.18605 |
| Olfactory transduction | 1 | 0.18605 |
| Amyotrophic lateral sclerosis (ALS) | 0.21257 | 0.18519 |
| Mitophagy - animal | 0.56534 | 0.18367 |
| Amino sugar and nucleotide sugar metabolism | 0.84649 | 0.18012 |
| Starch and sucrose metabolism | 0.9931 | 0.17808 |
| Insulin secretion | 0.24659 | 0.17544 |
| Endometrial cancer | 0.75932 | 0.17391 |
| Toll-like receptor signaling pathway | 0.5459 | 0.16867 |
| Cell adhesion molecules (CAMs) | 0.13173 | 0.16858 |
| Kaposi sarcoma-associated herpesvirus infection | 0.7258 | 0.16794 |
| Cellular senescence | 0.65078 | 0.16667 |
| Neuroactive ligand-receptor interaction | 0.35455 | 0.16477 |
| Cushing syndrome | 0.29952 | 0.16379 |
| Inflammatory bowel disease (IBD) | 0.34112 | 0.16129 |
| Thyroid cancer | 0.75236 | 0.16 |
| AMPK signaling pathway | 0.25637 | 0.15957 |
| Aldosterone synthesis and secretion | 0.0090582 | 0.15942 |
| Toxoplasmosis | 0.14265 | 0.15873 |
| Small cell lung cancer | 0.075347 | 0.15686 |
| Human immunodeficiency virus 1 infection | 0.84649 | 0.15672 |
| Hepatitis B | 0.24144 | 0.15556 |
| TNF signaling pathway | 0.15161 | 0.15464 |
| Salivary secretion | 0.3554 | 0.15385 |
| Vitamin B6 metabolism | 0.75236 | 0.15385 |
| Cortisol synthesis and secretion | 0.23856 | 0.15254 |

| | | |
|---|---|---|
| Drug metabolism - cytochrome P450 | 1 | 0.15094 |
| Influenza A | 0.37605 | 0.15044 |
| Cocaine addiction | 0.7258 | 0.14815 |
| Folate biosynthesis | 1 | 0.14607 |
| Amoebiasis | 4.44E-05 | 0.14035 |
| Sulfur metabolism | 0.34112 | 0.14035 |
| Autophagy - other | 0.22443 | 0.13889 |
| Endocytosis | 1.00E-06 | 0.13869 |
| Biosynthesis of unsaturated fatty acids | 0.23856 | 0.13636 |
| SNARE interactions in vesicular transport | 0.6877 | 0.13158 |
| Oocyte meiosis | 0.90397 | 0.12941 |
| NOD-like receptor signaling pathway | 0.44789 | 0.12727 |
| p53 signaling pathway | 0.45395 | 0.125 |
| Glucagon signaling pathway | 0.05326 | 0.12048 |
| Cardiac muscle contraction | 0.32416 | 0.11628 |
| Amphetamine addiction | 0.35539 | 0.11538 |
| Carbohydrate digestion and absorption | 0.4645 | 0.11475 |
| Tryptophan metabolism | 1 | 0.112 |
| Vasopressin-regulated water reabsorption | 0.35012 | 0.11111 |
| Nicotine addiction | 0.60391 | 0.10938 |
| Hypertrophic cardiomyopathy (HCM) | 0.015244 | 0.1087 |
| Nitrogen metabolism | 0.45028 | 0.1087 |
| Renin secretion | 0.06935 | 0.10714 |
| Hedgehog signaling pathway | 0.61806 | 0.10714 |
| Aldosterone-regulated sodium reabsorption | 0.85521 | 0.10714 |
| Glycosaminoglycan degradation | 1 | 0.10638 |
| Thiamine metabolism | 0.093625 | 0.10526 |
| Thyroid hormone synthesis | 0.13106 | 0.10526 |
| Antigen processing and presentation | 0.84649 | 0.10526 |
| Viral myocarditis | 1 | 0.096774 |
| Protein processing in endoplasmic reticulum | 0.13024 | 0.09434 |
| RIG-I-like receptor signaling pathway | 0.84649 | 0.09434 |
| African trypanosomiasis | 0.35882 | 0.09375 |
| Glycosylphosphatidylinositol (GPI)-anchor biosynthesis | 0.99187 | 0.093023 |
| Parkinson disease | 0.21422 | 0.089552 |
| Endocrine and other factor-regulated calcium reabsorption | 0.25637 | 0.089286 |
| Non-alcoholic fatty liver disease (NAFLD) | 0.92848 | 0.088235 |
| Ovarian steroidogenesis | 0.93135 | 0.088235 |
| Fatty acid biosynthesis | 0.21257 | 0.087209 |
| Legionellosis | 0.5437 | 0.086957 |
| Systemic lupus erythematosus | 1 | 0.085106 |

| | | |
|---|---|---|
| Intestinal immune network for IgA production | 0.34112 | 0.08 |
| Alcoholism | 0.89743 | 0.08 |
| Phagosome | 0.17496 | 0.078947 |
| Huntington disease | 0.6896 | 0.073529 |
| mRNA surveillance pathway | 1 | 0.071429 |
| Pancreatic secretion | 0.76268 | 0.066667 |
| RNA transport | 0.78941 | 0.065421 |
| Bladder cancer | 0.80618 | 0.064516 |
| MicroRNAs in cancer | 0.49665 | 0.063927 |
| Cell cycle | 0.90397 | 0.063636 |
| Central carbon metabolism in cancer | 3.28E-14 | 0.0625 |
| Cytosolic DNA-sensing pathway | 0.97297 | 0.0625 |
| Various types of N-glycan biosynthesis | 1 | 0.0625 |
| Alzheimer disease | 0.9469 | 0.059701 |
| alpha-Linolenic acid metabolism | 1 | 0.057692 |
| Ferroptosis | 0.017252 | 0.054054 |
| Phosphonate and phosphinate metabolism | 0.56534 | 0.052632 |
| Ribosome biogenesis in eukaryotes | 0.86163 | 0.05 |
| Terpenoid backbone biosynthesis | 0.84649 | 0.048387 |
| Prion diseases | 0.24577 | 0.046875 |
| Porphyrin and chlorophyll metabolism | 1 | 0.04321 |
| Cholesterol metabolism | 0.51597 | 0.042735 |
| Primary bile acid biosynthesis | 0.96919 | 0.042553 |
| Viral protein interaction with cytokine and cytokine receptor | 0.59154 | 0.034247 |
| Chemical carcinogenesis | 1 | 0.033784 |
| Arrhythmogenic right ventricular cardiomyopathy (ARVC) | 0.13024 | 0.031746 |
| Peroxisome | 1 | 0.03125 |
| Biotin metabolism | 0.95138 | 0.026316 |
| Glycosphingolipid biosynthesis - globo and isoglobo series | 0.99187 | 0.025641 |
| Neomycin, kanamycin and gentamicin biosynthesis | 0.95681 | 0.02381 |
| IL-17 signaling pathway | 0.84649 | 0.022727 |
| Rheumatoid arthritis | 0.11008 | 0.020408 |
| Fat digestion and absorption | 0.9137 | 0.015152 |
| Bile secretion | 1 | 0.014085 |
| Mineral absorption | 0.001875 | 0.011765 |
| Viral carcinogenesis | 0.17496 | 0.0072993 |
| Oxidative phosphorylation | 0.17496 | 0.0072993 |
| Transcriptional misregulation in cancer | 0.043076 | 0.0049505 |
| Protein digestion and absorption | 6.40E-13 | 0 |

| | | |
|---|---|---|
| ABC transporters | 4.24E-07 | 0 |
| Proximal tubule bicarbonate reclamation | 0.05326 | 0 |
| Sulfur relay system | 0.17661 | 0 |
| Vitamin digestion and absorption | 0.21973 | 0 |
| Lysosome | 0.23494 | 0 |
| D-Arginine and D-ornithine metabolism | 0.25637 | 0 |
| Antifolate resistance | 0.32491 | 0 |
| Malaria | 0.34112 | 0 |
| Proteasome | 0.35882 | 0 |
| Ubiquitin mediated proteolysis | 0.42183 | 0 |
| Primary immunodeficiency | 0.45028 | 0 |
| Protein export | 0.60266 | 0 |
| Renin-angiotensin system | 0.60391 | 0 |
| Nucleotide excision repair | 0.83433 | 0 |
| Mismatch repair | 0.85631 | 0 |
| Hematopoietic cell lineage | 0.86507 | 0 |
| Ascorbate and aldarate metabolism | 0.90397 | 0 |
| Glycosaminoglycan biosynthesis - heparan sulfate / heparin | 0.90534 | 0 |
| Caffeine metabolism | 0.92702 | 0 |
| RNA polymerase | 0.94522 | 0 |
| Glycosaminoglycan biosynthesis - chondroitin sulfate / dermatan sulfate | 0.95681 | 0 |
| Pentose and glucuronate interconversions | 0.96919 | 0 |
| Base excision repair | 0.96919 | 0 |
| DNA replication | 0.96919 | 0 |
| Phototransduction | 0.96919 | 0 |
| Collecting duct acid secretion | 0.96919 | 0 |
| Homologous recombination | 0.99801 | 0 |
| Basal transcription factors | 1 | 0 |
| Galactose metabolism | 1 | 0 |
| Fanconi anemia pathway | 1 | 0 |
| Spliceosome | 1 | 0 |
| Ribosome | 1 | 0 |
| RNA degradation | 1 | 0 |

**Table S1. Joint Pathway Analysis results upon integration of RNAseq and semi-targeted metabolomics in a batch of PK4A cells cultivated on CDM vs TCP.**

| Term | FDR | Impact |
|---|---|---|
| Central carbon metabolism in cancer | 6.76E-14 | 0.035714 |
| Protein digestion and absorption | 2.81E-11 | 0 |
| Alanine, aspartate and glutamate metabolism | 1.83E-08 | 1.2951 |
| ABC transporters | 1.01E-07 | 0 |
| Aminoacyl-tRNA biosynthesis | 1.50E-06 | 0.25773 |
| Glycolysis or Gluconeogenesis | 3.44E-05 | 1.5692 |
| Citrate cycle (TCA cycle) | 5.71E-05 | 1.5455 |
| Mineral absorption | 5.71E-05 | 0.011765 |
| Pyrimidine metabolism | 0.00016236 | 1.0543 |
| Purine metabolism | 0.00019549 | 1.3333 |
| Butanoate metabolism | 0.00091215 | 0.76364 |
| Glyoxylate and dicarboxylate metabolism | 0.0010108 | 0.40909 |
| Ferroptosis | 0.0011973 | 0.027027 |
| Arginine biosynthesis | 0.0012932 | 0.68571 |
| Valine, leucine and isoleucine biosynthesis | 0.0022745 | 0.5 |
| Pyruvate metabolism | 0.0030298 | 1.283 |
| Proximal tubule bicarbonate reclamation | 0.0038263 | 0 |
| Vitamin digestion and absorption | 0.0055546 | 0.0091743 |
| beta-Alanine metabolism | 0.0057947 | 1.0556 |
| Arginine and proline metabolism | 0.0057947 | 0.49573 |
| Cysteine and methionine metabolism | 0.0057947 | 0.52427 |
| Taurine and hypotaurine metabolism | 0.0057947 | 0.24138 |
| Glucagon signaling pathway | 0.0057947 | 0.24096 |
| Glutathione metabolism | 0.0057947 | 0.85507 |
| Arrhythmogenic right ventricular cardiomyopathy (ARVC) | 0.0065421 | 0.2381 |
| Synthesis and degradation of ketone bodies | 0.0069617 | 2.5 |
| Terpenoid backbone biosynthesis | 0.0069617 | 0.82258 |
| Pantothenate and CoA biosynthesis | 0.0075795 | 0.64286 |
| Peroxisome | 0.0095697 | 0 |
| Lysosome | 0.012118 | 0 |
| Amyotrophic lateral sclerosis (ALS) | 0.012459 | 0.25926 |
| ECM-receptor interaction | 0.013802 | 0.6875 |
| Osteoclast differentiation | 0.013802 | 0.34211 |
| cAMP signaling pathway | 0.014019 | 0.57377 |
| Steroid biosynthesis | 0.014389 | 0.89796 |
| Pentose phosphate pathway | 0.018733 | 0.71429 |
| Focal adhesion | 0.021289 | 0.58065 |
| Legionellosis | 0.036057 | 0.19565 |
| HIF-1 signaling pathway | 0.067306 | 0.21951 |
| Valine, leucine and isoleucine degradation | 0.072713 | 0.61458 |

| | | |
|---|---|---|
| PI3K-Akt signaling pathway | 0.087936 | 0.22772 |
| AMPK signaling pathway | 0.087936 | 0.074468 |
| Rap1 signaling pathway | 0.095781 | 0.6383 |
| Thiamine metabolism | 0.095781 | 0.10526 |
| Fluid shear stress and atherosclerosis | 0.095781 | 0.44762 |
| Synaptic vesicle cycle | 0.095781 | 0.029412 |
| Taste transduction | 0.099396 | 0.31429 |
| Aldosterone synthesis and secretion | 0.10856 | 0.11594 |
| PPAR signaling pathway | 0.10856 | 0.37288 |
| Fatty acid biosynthesis | 0.10856 | 0.098837 |
| MAPK signaling pathway | 0.11053 | 0.35484 |
| Renin secretion | 0.11053 | 0.089286 |
| Estrogen signaling pathway | 0.11335 | 0.22078 |
| Viral carcinogenesis | 0.12368 | 0 |
| Phagosome | 0.13295 | 0.078947 |
| Axon guidance | 0.13295 | 0.27517 |
| Insulin resistance | 0.13295 | 0.095588 |
| Nicotinate and nicotinamide metabolism | 0.13476 | 0.29268 |
| Phenylalanine metabolism | 0.13846 | 0.27397 |
| Fatty acid degradation | 0.14627 | 0.57983 |
| Histidine metabolism | 0.16779 | 0.29032 |
| D-Glutamine and D-glutamate metabolism | 0.17743 | 0.375 |
| Phenylalanine, tyrosine and tryptophan biosynthesis | 0.18263 | 0.43902 |
| Complement and coagulation cascades | 0.18341 | 0.32308 |
| Antifolate resistance | 0.18341 | 0 |
| Leishmaniasis | 0.18572 | 0.26415 |
| Glycine, serine and threonine metabolism | 0.18572 | 0.375 |
| Human papillomavirus infection | 0.18783 | 0.1625 |
| Malaria | 0.19921 | 0 |
| Sulfur relay system | 0.21036 | 0 |
| Thyroid hormone synthesis | 0.21318 | 0.031579 |
| Endocytosis | 0.21318 | 0.087591 |
| Toxoplasmosis | 0.21318 | 0.14286 |
| Amphetamine addiction | 0.21391 | 0.17308 |
| Salmonella infection | 0.22347 | 0.20833 |
| NF-kappa B signaling pathway | 0.22486 | 0.23125 |
| Glycerophospholipid metabolism | 0.22964 | 0.7619 |
| Propanoate metabolism | 0.24004 | 0.5 |
| Glycerolipid metabolism | 0.24909 | 0.58621 |
| Neuroactive ligand-receptor interaction | 0.24909 | 0.14205 |
| Hippo signaling pathway | 0.2625 | 0.35294 |

| | | |
|---|---|---|
| Nitrogen metabolism | 0.2625 | 0.1087 |
| Amoebiasis | 0.27941 | 0.052632 |
| Pancreatic secretion | 0.30792 | 0.011111 |
| Staphylococcus aureus infection | 0.30792 | 0.41026 |
| Th17 cell differentiation | 0.31614 | 0.17143 |
| Fructose and mannose metabolism | 0.32126 | 0.59459 |
| Hepatitis B | 0.32497 | 0.14815 |
| Th1 and Th2 cell differentiation | 0.32497 | 0.29577 |
| Cocaine addiction | 0.32497 | 0.08642 |
| D-Arginine and D-ornithine metabolism | 0.32497 | 0 |
| cGMP-PKG signaling pathway | 0.32749 | 0.23171 |
| Human cytomegalovirus infection | 0.32749 | 0.22086 |
| JAK-STAT signaling pathway | 0.33181 | 1.5278 |
| Maturity onset diabetes of the young | 0.36278 | 0.14706 |
| Hypertrophic cardiomyopathy (HCM) | 0.36278 | 0.1087 |
| TNF signaling pathway | 0.37044 | 0.072165 |
| Riboflavin metabolism | 0.38057 | 0.20833 |
| Galactose metabolism | 0.38057 | 0.3662 |
| Alcoholism | 0.38057 | 0.19 |
| Lysine degradation | 0.38057 | 0.1039 |
| Human T-cell leukemia virus 1 infection | 0.38057 | 0.20769 |
| Insulin secretion | 0.3973 | 0.070175 |
| RNA degradation | 0.43159 | 0.048387 |
| Prolactin signaling pathway | 0.44058 | 0.15152 |
| Ras signaling pathway | 0.44058 | 0.53125 |
| Apoptosis | 0.44058 | 0.13514 |
| Natural killer cell mediated cytotoxicity | 0.44058 | 0.16495 |
| Morphine addiction | 0.44944 | 0.22 |
| Kaposi sarcoma-associated herpesvirus infection | 0.44944 | 0.19084 |
| C-type lectin receptor signaling pathway | 0.47188 | 0.21579 |
| Rheumatoid arthritis | 0.51969 | 0.020408 |
| Colorectal cancer | 0.51969 | 0.14865 |
| Renal cell carcinoma | 0.52869 | 0.10526 |
| Pathways in cancer | 0.52869 | 0.20561 |
| Cellular senescence | 0.52897 | 0.19298 |
| Antigen processing and presentation | 0.52897 | 0.42105 |
| Thermogenesis | 0.52897 | 0.060241 |
| Cell adhesion molecules (CAMs) | 0.52897 | 0.095785 |
| Proteoglycans in cancer | 0.52897 | 0.155 |
| Cytokine-cytokine receptor interaction | 0.52897 | 0.125 |
| Type II diabetes mellitus | 0.52897 | 0.22581 |

| | | |
|---|---|---|
| PD-L1 expression and PD-1 checkpoint pathway in cancer | 0.52897 | 0.1875 |
| Sphingolipid metabolism | 0.52897 | 1.0984 |
| Long-term potentiation | 0.54464 | 0.58065 |
| Tuberculosis | 0.54748 | 0.15441 |
| Hematopoietic cell lineage | 0.54748 | 0 |
| Relaxin signaling pathway | 0.55636 | 0.28889 |
| TGF-beta signaling pathway | 0.55636 | 0.3 |
| Vascular smooth muscle contraction | 0.55837 | 0.32394 |
| Ascorbate and aldarate metabolism | 0.55837 | 0.090909 |
| Parkinson disease | 0.56507 | 0.089552 |
| Parathyroid hormone synthesis, secretion and action | 0.56956 | 0.13415 |
| Dilated cardiomyopathy (DCM) | 0.56956 | 0.15217 |
| Adipocytokine signaling pathway | 0.57816 | 0.30233 |
| GABAergic synapse | 0.57816 | 0.19643 |
| Leukocyte transendothelial migration | 0.58635 | 0.28571 |
| Chemokine signaling pathway | 0.59299 | 0.26562 |
| Inflammatory bowel disease (IBD) | 0.59537 | 0.14516 |
| Biosynthesis of unsaturated fatty acids | 0.61615 | 0.037879 |
| Necroptosis | 0.62835 | 0.095745 |
| Autophagy - animal | 0.62835 | 0.1 |
| Mucin type O-glycan biosynthesis | 0.63421 | 0.625 |
| Mitophagy - animal | 0.63421 | 0.16327 |
| Apelin signaling pathway | 0.63421 | 0.36 |
| Basal transcription factors | 0.63421 | 0 |
| Selenocompound metabolism | 0.63421 | 0.19048 |
| Sulfur metabolism | 0.63421 | 0.21053 |
| Neurotrophin signaling pathway | 0.63421 | 0.26829 |
| RNA transport | 0.63421 | 0.056075 |
| Influenza A | 0.63421 | 0.12389 |
| Melanogenesis | 0.63421 | 0.20455 |
| Vasopressin-regulated water reabsorption | 0.63421 | 0.074074 |
| Oxidative phosphorylation | 0.63421 | 0.0072993 |
| Neomycin, kanamycin and gentamicin biosynthesis | 0.63421 | 0.047619 |
| Pertussis | 0.63421 | 0.13433 |
| B cell receptor signaling pathway | 0.63421 | 0.14286 |
| Circadian entrainment | 0.63421 | 0.62069 |
| Cushing syndrome | 0.64169 | 0.15517 |
| Chagas disease (American trypanosomiasis) | 0.64169 | 0.11111 |
| Graft-versus-host disease | 0.65089 | 0.086957 |
| Prostate cancer | 0.65951 | 0.16393 |

| | | |
|---|---|---|
| Endocrine and other factor-regulated calcium reabsorption | 0.65951 | 0.14286 |
| Pentose and glucuronate interconversions | 0.65951 | 0.088235 |
| One carbon pool by folate | 0.66814 | 0.93333 |
| Fatty acid elongation | 0.66982 | 0.18987 |
| Regulation of actin cytoskeleton | 0.67359 | 0.24096 |
| Starch and sucrose metabolism | 0.68092 | 0.34247 |
| Acute myeloid leukemia | 0.69478 | 0.20833 |
| Calcium signaling pathway | 0.69478 | 0.28358 |
| GnRH secretion | 0.70645 | 0.10638 |
| Ubiquitin mediated proteolysis | 0.70656 | 0 |
| Signaling pathways regulating pluripotency of stem cells | 0.70656 | 0.20408 |
| Bacterial invasion of epithelial cells | 0.70656 | 0.11628 |
| Small cell lung cancer | 0.70656 | 0.13725 |
| mRNA surveillance pathway | 0.70656 | 0.19643 |
| Sphingolipid signaling pathway | 0.70656 | 0.036585 |
| Adrenergic signaling in cardiomyocytes | 0.71282 | 0.22535 |
| Tyrosine metabolism | 0.71283 | 0.41129 |
| Fat digestion and absorption | 0.71283 | 0.030303 |
| Gap junction | 0.71283 | 0.096154 |
| Apoptosis - multiple species | 0.71283 | 0.030303 |
| Circadian rhythm | 0.71283 | 0.5 |
| Inositol phosphate metabolism | 0.71283 | 0.23529 |
| Longevity regulating pathway | 0.71283 | 0.074074 |
| Fc gamma R-mediated phagocytosis | 0.71283 | 0.12963 |
| Oxytocin signaling pathway | 0.71283 | 0.25 |
| Glutamatergic synapse | 0.71926 | 0.31429 |
| Dopaminergic synapse | 0.74696 | 0.25 |
| Cholinergic synapse | 0.7512 | 0.14754 |
| Protein processing in endoplasmic reticulum | 0.75211 | 0.018868 |
| Collecting duct acid secretion | 0.75211 | 0 |
| Phototransduction | 0.75211 | 0.042553 |
| Cortisol synthesis and secretion | 0.76948 | 0.084746 |
| Endometrial cancer | 0.77338 | 0.17391 |
| Drug metabolism - other enzymes | 0.77338 | 0.26667 |
| Prion diseases | 0.77338 | 0.015625 |
| Thyroid cancer | 0.77338 | 0.2 |
| Vitamin B6 metabolism | 0.77338 | 0.38462 |
| Thyroid hormone signaling pathway | 0.77338 | 0.17925 |
| EGFR tyrosine kinase inhibitor resistance | 0.77338 | 0.69697 |
| Tryptophan metabolism | 0.78 | 0.248 |
| Gastric cancer | 0.78505 | 0.20388 |

| | | |
|---|---|---|
| Choline metabolism in cancer | 0.80041 | 0.09434 |
| Phosphonate and phosphinate metabolism | 0.80041 | 0 |
| Amino sugar and nucleotide sugar metabolism | 0.80041 | 0.25466 |
| mTOR signaling pathway | 0.80041 | 0.11905 |
| Basal cell carcinoma | 0.80041 | 0.12 |
| VEGF signaling pathway | 0.80041 | 0.15152 |
| Phospholipase D signaling pathway | 0.80041 | 0.28986 |
| Viral myocarditis | 0.80041 | 0.12903 |
| N-Glycan biosynthesis | 0.80041 | 0.21519 |
| ErbB signaling pathway | 0.80041 | 0.46377 |
| FoxO signaling pathway | 0.80379 | 0.071429 |
| Cytosolic DNA-sensing pathway | 0.80441 | 0.15625 |
| Epstein-Barr virus infection | 0.80726 | 0.24138 |
| Inflammatory mediator regulation of TRP channels | 0.81195 | 0.13483 |
| Hepatitis C | 0.81195 | 0.14433 |
| Platelet activation | 0.81757 | 0.17442 |
| Nucleotide excision repair | 0.82163 | 0 |
| Ribosome biogenesis in eukaryotes | 0.82163 | 0 |
| Intestinal immune network for IgA production | 0.82895 | 0.02 |
| alpha-Linolenic acid metabolism | 0.82895 | 0.28846 |
| Tight junction | 0.82895 | 0.17778 |
| Insulin signaling pathway | 0.82895 | 0.18182 |
| Regulation of lipolysis in adipocytes | 0.82895 | 0.033333 |
| Long-term depression | 0.82895 | 0.15217 |
| Salivary secretion | 0.82895 | 0.061538 |
| Growth hormone synthesis, secretion and action | 0.82895 | 0.15873 |
| Adherens junction | 0.82895 | 0.41333 |
| African trypanosomiasis | 0.82895 | 0 |
| Aldosterone-regulated sodium reabsorption | 0.82895 | 0.14286 |
| GnRH signaling pathway | 0.82895 | 0.12766 |
| Nicotine addiction | 0.83407 | 0.078125 |
| Oocyte meiosis | 0.83407 | 0.15294 |
| Type I diabetes mellitus | 0.83407 | 0 |
| Carbohydrate digestion and absorption | 0.83407 | 0 |
| Bile secretion | 0.83407 | 0.0070423 |
| Serotonergic synapse | 0.83407 | 0.12821 |
| Hedgehog signaling pathway | 0.83407 | 0.10714 |
| Toll-like receptor signaling pathway | 0.83407 | 0.072289 |
| Mismatch repair | 0.83407 | 0 |
| Breast cancer | 0.8375 | 0.19167 |
| Mannose type O-glycan biosynthesis | 0.84393 | 0.1875 |

| | | |
|---|---|---|
| Chronic myeloid leukemia | 0.84393 | 0.090909 |
| Viral protein interaction with cytokine and cytokine receptor | 0.84393 | 0.027397 |
| Endocrine resistance | 0.84393 | 0.34783 |
| Fanconi anemia pathway | 0.84393 | 0.074074 |
| Various types of N-glycan biosynthesis | 0.84393 | 0.1 |
| MicroRNAs in cancer | 0.84393 | 0.031963 |
| Alzheimer disease | 0.84393 | 0.1194 |
| Glioma | 0.84393 | 0.19178 |
| Linoleic acid metabolism | 0.84393 | 0.73171 |
| Transcriptional misregulation in cancer | 0.85682 | 0.009901 |
| T cell receptor signaling pathway | 0.85682 | 0.23944 |
| Notch signaling pathway | 0.85682 | 0.074074 |
| Platinum drug resistance | 0.85682 | 0 |
| Ovarian steroidogenesis | 0.85682 | 0.014706 |
| Hippo signaling pathway - multiple species | 0.85682 | 0.093023 |
| Wnt signaling pathway | 0.87724 | 0.47561 |
| NOD-like receptor signaling pathway | 0.87724 | 0.20606 |
| Caffeine metabolism | 0.87724 | 0 |
| Protein export | 0.87724 | 0 |
| Human immunodeficiency virus 1 infection | 0.88593 | 0.15672 |
| Cholesterol metabolism | 0.89317 | 0.042735 |
| RNA polymerase | 0.89766 | 0 |
| Biotin metabolism | 0.90843 | 0.026316 |
| Measles | 0.91144 | 0.1236 |
| Primary bile acid biosynthesis | 0.91569 | 0.042553 |
| Ribosome | 0.91569 | 0 |
| SNARE interactions in vesicular transport | 0.91569 | 0.026316 |
| Allograft rejection | 0.91569 | 0 |
| Longevity regulating pathway - multiple species | 0.91569 | 0.09434 |
| Cell cycle | 0.92598 | 0.090909 |
| DNA replication | 0.92792 | 0 |
| Autophagy - other | 0.92792 | 0.083333 |
| Yersinia infection | 0.92962 | 0.16842 |
| Primary immunodeficiency | 0.92962 | 0 |
| RIG-I-like receptor signaling pathway | 0.92962 | 0.22642 |
| Phosphatidylinositol signaling system | 0.93049 | 0.51948 |
| Hepatocellular carcinoma | 0.94268 | 0.12712 |
| Glycosylphosphatidylinositol (GPI)-anchor biosynthesis | 0.94268 | 0.11628 |
| Non-small cell lung cancer | 0.94268 | 0.071429 |
| Ether lipid metabolism | 0.94268 | 0.51163 |
| Bladder cancer | 0.95605 | 0 |

| | | |
|---|---|---|
| Herpes simplex virus 1 infection | 0.96657 | 0.20455 |
| Fc epsilon RI signaling pathway | 0.96657 | 0.06 |
| Pancreatic cancer | 0.96657 | 0.070175 |
| AGE-RAGE signaling pathway in diabetic complications | 0.96657 | 0.089552 |
| Proteasome | 0.97309 | 0 |
| Autoimmune thyroid disease | 0.97309 | 0 |
| Renin-angiotensin system | 0.97309 | 0 |
| Folate biosynthesis | 0.97309 | 0.089888 |
| Glycosaminoglycan degradation | 0.97309 | 0.19149 |
| Retinol metabolism | 0.97309 | 0.69492 |
| Gastric acid secretion | 0.98641 | 0.057692 |
| Porphyrin and chlorophyll metabolism | 0.98641 | 0.080247 |
| Cardiac muscle contraction | 0.98972 | 0 |
| Progesterone-mediated oocyte maturation | 1 | 0.1 |
| Spliceosome | 1 | 0 |
| Huntington disease | 1 | 0.029412 |
| Retrograde endocannabinoid signaling | 1 | 0.16667 |
| p53 signaling pathway | 1 | 0.015625 |
| Melanoma | 1 | 0.030303 |
| Non-alcoholic fatty liver disease (NAFLD) | 1 | 0.044118 |
| Glycosphingolipid biosynthesis - lacto and neolacto series | 1 | 0.12281 |
| Arachidonic acid metabolism | 1 | 1.188 |
| IL-17 signaling pathway | 1 | 0.022727 |
| Systemic lupus erythematosus | 1 | 0.12766 |
| Ubiquinone and other terpenoid-quinone biosynthesis | 1 | 0.011628 |
| Drug metabolism - cytochrome P450 | 1 | 0.10377 |
| Metabolism of xenobiotics by cytochrome P450 | 1 | 0.35165 |
| Chemical carcinogenesis | 1 | 0.033784 |
| Steroid hormone biosynthesis | 1 | 0.11168 |
| Olfactory transduction | 1 | 0.23256 |

**Table S2. Joint Pathway Analysis results upon integration of RNAseq and semi-targeted metabolomics in a batch of PK4A cells (different from the one in Table S1) cultivated on CDM vs TCP.**

| | Mean | | |
|---|---|---|---|
| | TCP | CDM | -log10pval |
| IMP | -0.226401547 | 0.457420459 | 2.190796625 |
| Inosine | 0.000458525 | 0.53197057 | 2.003640706 |
| Guanine | -0.189290376 | 0.407560324 | 1.974304971 |
| Guanosine | -0.0737993 | 0.504480785 | 1.875845263 |
| Hypoxanthine | -0.149919098 | -0.11798106 | 1.231610614 |
| ATP | 0.26103179 | 0.091351308 | 1.230673655 |
| Ribose phosphate | -0.266156981 | 0.146469595 | 0.91847204 |
| GDP | -0.027149543 | 0.287546745 | 0.666986369 |
| Xanthine | -0.175084913 | -0.058978471 | 0.655981567 |
| Glutamine | -0.18312295 | -0.177412675 | 0.55499531 |
| GMP | -0.10695879 | 0.146234945 | 0.554801186 |
| AMP | -0.114221145 | -0.02117759 | 0.536578512 |
| Urate | -0.17842375 | -0.151572701 | 0.349426858 |
| ADP | 0.033378498 | 0.16382652 | 0.287413632 |
| Adenine | 0.298279978 | 0.164078662 | 0.186116548 |
| Adenosine | 0.157492325 | 0.06811415 | 0.101585605 |
| GTP | 0.139768028 | 0.179841348 | 0.090120041 |

**Table S3. Purine metabolism compounds in PK4A cells on CDM compared to TCP.**

| | TCP | | | CDM | | | | |
|---|---|---|---|---|---|---|---|---|
| | 1 | 2 | 3 | 1 | 2 | 3 | -LOG10 adj. p val | log2FC |
| Nme1 | 3018 | 2628 | 2555 | 3706 | 4385 | 4806 | 4.75328033 | 0.653163774 |
| Gda | 346 | 293 | 696 | 116 | 30 | 11 | 4.358873854 | -3.088003278 |
| Nme2 | 4254 | 3507 | 3417 | 4677 | 6195 | 5585 | 4.275345424 | 0.558039286 |
| Hprt | 1829 | 1386 | 1635 | 1998 | 1910 | 1949 | 2.393500704 | 0.272177147 |
| Nme3 | 255 | 186 | 210 | 197 | 126 | 163 | 2.038388799 | -0.42170123 |
| Nme7 | 645 | 448 | 568 | 503 | 361 | 454 | 1.710413808 | -0.333701703 |
| Impdh1 | 1317 | 1275 | 1248 | 1529 | 1758 | 1722 | 1.70188637 | 0.383416301 |
| Pnp | 672 | 576 | 645 | 785 | 941 | 892 | 1.484442925 | 0.467790686 |
| Gmpr | 131 | 141 | 89 | 85 | 78 | 71 | 1.354629543 | -0.625490307 |
| Aprt | 3900 | 3290 | 3464 | 5189 | 6411 | 5685 | 10.20753977 | 0.698125418 |
| Pde4a | 1801 | 1110 | 1530 | 1207 | 1039 | 1026 | 5.78330991 | -0.440711823 |
| Nudt5 | 557 | 485 | 523 | 683 | 744 | 677 | 3.523149961 | 0.426972047 |
| Ampd3 | 288 | 196 | 322 | 431 | 1745 | 219 | 1.708819632 | 1.571173912 |
| Pde4b | 625 | 749 | 944 | 1676 | 1506 | 1196 | 1.598966719 | 0.917391388 |
| Enpp1 | 962 | 441 | 1255 | 221 | 165 | 139 | 12.21324838 | -2.339951777 |
| Nt5c | 863 | 578 | 824 | 1190 | 1135 | 1318 | 6.09611753 | 0.685615945 |
| Adcy3 | 825 | 583 | 680 | 357 | 419 | 250 | 5.460343118 | -1.025090981 |
| Adcy8 | 361 | 110 | 215 | 87 | 29 | 111 | 2.044674364 | -1.595516279 |

**Table S3. Purine metabolism gene raw values in PK4A cells on CDM compared to TCP.**

| | TCP | | | CDM | | | | |
|---|---|---|---|---|---|---|---|---|
| | 1 | 2 | 3 | 1 | 2 | 3 | -log10 p adj. | log2FC |
| Upp1 | 1751 | 1932 | 2087 | 3515 | 4794 | 3735 | 7.739071658 | 1.061671 |
| Uckl1 | 570 | 731 | 648 | 931 | 1479 | 828 | 1.649261981 | 0.732369 |
| Uck2 | 4790 | 3299 | 5115 | 5649 | 6274 | 5007 | 0.986900254 | 0.358607 |
| Ctps2 | 1354 | 1161 | 1236 | 1174 | 1340 | 1097 | 0.434762602 | -0.05488 |
| Umps | 5149 | 4355 | 4008 | 4111 | 4307 | 4429 | 0.42920692 | -0.07281 |
| Hddc2 | 407 | 364 | 301 | 421 | 430 | 435 | 0.415917457 | 0.262586 |
| Dctd | 1284 | 1018 | 1387 | 1267 | 1070 | 1092 | 0.397369586 | -0.10544 |
| Cad | 10184 | 9226 | 8024 | 9537 | 10595 | 11943 | 0.376917923 | 0.225484 |
| Tyms | 1501 | 1016 | 1285 | 1253 | 997 | 1355 | 0.168199 | -0.07676 |
| Dtymk | 1454 | 1178 | 1265 | 1361 | 1364 | 1605 | 0.158398933 | 0.152003 |
| Uck1 | 1017 | 692 | 843 | 1007 | 797 | 949 | 0.106937628 | 0.109376 |
| Cmpk1 | 1248 | 913 | 1155 | 1158 | 1066 | 1130 | 0.104388318 | 0.016439 |
| Tk1 | 3739 | 3098 | 2448 | 2937 | 2566 | 3552 | 0.095110384 | -0.03619 |
| Dctpp1 | 1060 | 870 | 831 | 997 | 899 | 1194 | 0.094698173 | 0.162416 |
| Cda | 834 | 815 | 836 | 855 | 966 | 755 | 0.03665178 | 0.051887 |
| Dut | 3497 | 2965 | 2870 | 3487 | 2990 | 3463 | 0.026508141 | 0.09106 |
| Tymp | 1523 | 907 | 1290 | 1329 | 1193 | 1348 | 0.01799685 | 0.057031 |
| Dhodh | 616 | 524 | 508 | 608 | 581 | 557 | 0.01519702 | 0.083337 |
| Tk2 | 423 | 225 | 494 | 496 | 325 | 354 | 0.000810071 | 0.041098 |

**Table S5. Pyrimidine metabolism gene raw values in PK4A cells on CDM compared to TCP.**

| | Mean | | |
|---|---|---|---|
| | TCP | CDM | -log10pval |
| UDP | -0.160903925 | 0.371136425 | 2.902890879 |
| CDP | 0.062855003 | 0.560278208 | 2.459603681 |
| UMP | -0.246809035 | 0.291831103 | 2.21075114 |
| Cytidine | -0.161290038 | 0.408377526 | 1.432032567 |
| CMP | -0.271110088 | 0.250932208 | 1.107942342 |
| UTP | 0.170726153 | 0.311177953 | 0.801571495 |
| L-Dihydroorotic acid | 0.046434651 | 0.377343959 | 0.676753998 |
| beta-Alanine | -0.11717631 | -0.222280671 | 0.590589319 |
| CTP | -0.23451595 | -0.178524743 | 0.553064484 |
| Orotic acid | 0.081224443 | 0.16296641 | 0.09251722 |
| N-carbamoyl-L-aspartic acid | 0.062575423 | 0.072642551 | 0.015222814 |
| Methylmalonic acid | -0.118741303 | -0.118745025 | 0.014565963 |

**Table S6. Pyrimidine metabolism gene raw values in PK4A cells on CDM compared to TCP.**

| GeneSymbol | CDM_1 | CDM_2 | CDM_3 | TCP_1 | TCP_2 | TCP_3 | log2FoldChange | padj. |
|---|---|---|---|---|---|---|---|---|
| Ercc1 | 8400.729903 | 8944.258227 | 6904.610478 | 4122.817782 | 3876.543181 | 3328.785032 | 1.109527176 | 1.99115E-12 |
| Aprt | 5272.482833 | 5867.352859 | 4490.852817 | 3736.658541 | 3110.573449 | 3326.226396 | 0.631460039 | 6.20098E-11 |
| Stx3 | 331.095228 | 309.3378984 | 347.9134385 | 145.3775967 | 184.9821393 | 167.1641984 | 0.998546406 | 9.21373E-08 |
| Nt5c | 1222.362775 | 1038.753002 | 1029.893014 | 656.4707102 | 739.9285571 | 736.0341999 | 0.636569181 | 8.01461E-07 |
| Polr1d | 2121.97726 | 1741.62728 | 1874.578378 | 1493.527654 | 1248.180454 | 1419.189929 | 0.477089502 | 1.88826E-06 |
| Pold4 | 1158.369579 | 1716.916856 | 879.3036157 | 575.8315745 | 608.8247108 | 592.7506013 | 1.088566665 | 3.04026E-06 |
| Nme1 | 4457.265171 | 4013.155871 | 3207.381102 | 2984.783783 | 2294.31731 | 2573.987503 | 0.584892825 | 1.7649E-05 |
| Tsg101 | 1855.802665 | 2269.698189 | 1775.050901 | 1504.885279 | 1352.345154 | 1350.959644 | 0.499471812 | 3.9643E-05 |
| Gtf2f1 | 2976.147302 | 2552.495262 | 2550.499759 | 2199.971913 | 2127.294602 | 2137.313679 | 0.333926488 | 0.00033711 |
| Vps37b | 5006.308238 | 7085.485234 | 5252.454374 | 3096.088505 | 3313.515019 | 2847.761522 | 0.916332753 | 0.000442991 |
| Bola2 | 1147.240328 | 1118.375479 | 988.3511113 | 930.1894666 | 790.2149639 | 844.3497774 | 0.356798349 | 0.000998538 |
| Zwint | 4221.696017 | 4433.233076 | 4086.684718 | 3238.058815 | 3287.473844 | 3068.65707 | 0.420962667 | 0.001663207 |
| Cox17 | 698.360523 | 898.7272668 | 619.6667213 | 624.669361 | 449.8837465 | 526.2260734 | 0.483165928 | 0.002807434 |
| Hprt | 1807.575909 | 1748.033686 | 1729.181717 | 1574.16679 | 1468.183484 | 1559.914892 | 0.211945365 | 0.004041097 |
| Adrm1 | 881.9931705 | 895.9816642 | 719.1941975 | 693.9508719 | 550.4565601 | 676.3327005 | 0.391156839 | 0.009625035 |
| Itpa | 770.7006568 | 730.3303044 | 628.3212844 | 565.6097123 | 519.0275558 | 553.5181874 | 0.390387148 | 0.009766564 |
| Edf1 | 2168.34914 | 2297.154216 | 1748.221756 | 1726.358961 | 1568.756298 | 1589.765642 | 0.358833787 | 0.011726929 |
| Ddb1 | 31447.5546 | 27258.34309 | 27390.8269 | 25571.69211 | 24028.8207 | 26260.98384 | 0.194835503 | 0.015585509 |
| Rala | 2067.258441 | 2679.708185 | 1844.287406 | 1987.58433 | 1603.777188 | 1559.062013 | 0.367513083 | 0.024121484 |
| Pde4b | 1109.215386 | 1378.292529 | 1450.504783 | 850.6860934 | 847.6851431 | 533.0491019 | 0.831165035 | 0.025178699 |
| Polb | 325.5306023 | 344.1155319 | 397.2444484 | 215.7948701 | 314.2900425 | 267.8038688 | 0.426869614 | 0.027765582 |
| Gtf2h1 | 1699.993146 | 1831.316966 | 1761.2036 | 1289.090409 | 1588.511672 | 1422.601443 | 0.310346506 | 0.04696194 |
| Guk1 | 1246.476153 | 1135.764295 | 1066.24218 | 1041.494189 | 839.6033991 | 898.0811269 | 0.324447355 | 0.062414088 |
| Gpx4 | 934.8571144 | 1251.079607 | 872.3799652 | 913.1530295 | 649.2334306 | 825.586449 | 0.36895443 | 0.06495945 |

| | | | | | | | | |
|---|---|---|---|---|---|---|---|---|
| Polr2g | 1226.072525 | 1273.044428 | 1060.183985 | 1218.673135 | 831.5216552 | 898.9340055 | 0.285655694 | 0.065538456 |
| Ercc3 | 1522.852561 | 1524.724671 | 1324.148161 | 1379.951406 | 1214.955507 | 1301.492687 | 0.17866422 | 0.080608298 |
| Aaas | 1648.056639 | 1593.364737 | 1357.900958 | 1453.775967 | 1167.363015 | 1269.93618 | 0.254018166 | 0.095834379 |
| Taf10 | 1598.902446 | 2061.032388 | 1479.064842 | 1560.53764 | 1199.689991 | 1329.63768 | 0.341045744 | 0.097005004 |
| Gmpr2 | 901.4693604 | 756.87113 | 845.5508194 | 594.0037741 | 783.0311915 | 736.0341999 | 0.254408047 | 0.10496352 |
| Ercc2 | 972.8820566 | 917.0312845 | 961.5219656 | 909.745742 | 853.0729724 | 828.9979633 | 0.151469522 | 0.163958317 |
| Tarbp2 | 378.3945463 | 314.8291037 | 326.2770306 | 327.0995926 | 261.309721 | 293.3902257 | 0.224841287 | 0.201062371 |
| Pole4 | 697.4330854 | 779.7511521 | 738.2342364 | 606.4971614 | 706.7036097 | 682.3028504 | 0.159949967 | 0.214998369 |
| Supt5 | 6321.414774 | 5580.894983 | 5971.648571 | 5208.606708 | 5712.895 | 5536.034753 | 0.131000103 | 0.224686565 |
| Rae1 | 2128.469323 | 2254.139774 | 2210.375428 | 1906.945195 | 2009.660329 | 1957.356302 | 0.178113601 | 0.228623877 |
| Sdcbp | 5195.505511 | 5853.624846 | 4552.300215 | 5202.927895 | 4354.264045 | 4492.111392 | 0.162833879 | 0.245628812 |
| Tmed2 | 2134.961386 | 2158.043681 | 1883.232941 | 2026.200254 | 1735.779006 | 1846.482089 | 0.151512628 | 0.257321495 |
| Polr2f | 1161.151892 | 1197.997955 | 897.4781983 | 1111.911462 | 777.6433622 | 939.0192979 | 0.216245824 | 0.266373344 |
| Ada | 57.50113204 | 33.86243266 | 45.86918468 | 26.12253691 | 38.61277664 | 29.85074971 | 0.543295488 | 0.279008251 |
| Vps28 | 788.3219715 | 918.8616862 | 667.2668186 | 797.3052571 | 650.1314021 | 693.3902718 | 0.162256048 | 0.325330691 |
| Ercc4 | 580.5759461 | 548.2053287 | 528.7938082 | 533.808363 | 487.5985516 | 521.9616806 | 0.115947397 | 0.327449584 |
| Dad1 | 1065.625818 | 1108.308269 | 753.81245 | 1068.752488 | 691.4380934 | 810.2346349 | 0.201160556 | 0.340456834 |
| Pde6g | 2.782312841 | 0 | 1.730912629 | 0 | 0.89797155 | 0 | 2.15043376 | 0.345234487 |
| Ccno | 768.8457816 | 1906.363439 | 716.5978285 | 1393.580556 | 300.8204692 | 319.8294611 | 0.761273061 | 0.353037971 |
| Ssrp1 | 12665.08805 | 11699.92808 | 11372.09597 | 11405.32677 | 10770.27077 | 11373.98852 | 0.103186577 | 0.358346168 |
| Polr2h | 721.5464633 | 566.5093464 | 727.8487606 | 539.4871754 | 608.8247108 | 682.3028504 | 0.149079612 | 0.35962841 |
| Ercc5 | 484.1224343 | 419.1620043 | 423.2081379 | 407.7387283 | 414.862856 | 386.3539891 | 0.146119509 | 0.379479955 |
| Taf12 | 719.6915881 | 665.3510417 | 614.4739834 | 618.9905486 | 638.457772 | 606.3966583 | 0.112655191 | 0.400800468 |
| Gtf2h5 | 1085.102008 | 974.6889401 | 958.9255966 | 1009.69284 | 792.010907 | 936.4606622 | 0.153879996 | 0.427090244 |
| Pcna | 4189.2357 | 4209.92406 | 3128.624577 | 4311.354353 | 2352.685461 | 3530.064372 | 0.189645767 | 0.443284922 |

| | | | | | | | | |
|---|---|---|---|---|---|---|---|---|
| Polr2e | 2827.757284 | 3182.153469 | 2599.830769 | 3026.806994 | 2370.644892 | 2595.309467 | 0.119414572 | 0.455050453 |
| Sac3d1 | 673.3197074 | 803.546375 | 647.3613233 | 747.3317082 | 523.5174136 | 585.9275728 | 0.207037369 | 0.4647524 |
| Alyref | 2415.974983 | 2946.946842 | 1820.920086 | 2722.422651 | 1691.7784 | 2020.469316 | 0.17063112 | 0.470136756 |

**Table S7. Hallmark DNA Repair gene set (42) in PK4A cells grown on CDM vs TCP.**

| Term | Adjusted P-value | Combined Score | Genes |
|---|---|---|---|
| Nucleotide-Excision Repair (GO:0006289) | 0.006906542 | 616.5629353 | DDB1;POLB;ERCC1 |
| UV-damage Excision Repair (GO:0070914) | 0.007785371 | 1894.647073 | DDB1;ERCC1 |
| Protein Transport To Vacuole Involved In Ubiquitin-Dependent Protein Catabolic Process Via The Multivesicular Body Sorting Pathway (GO:0043328) | 0.007785371 | 1525.492666 | TSG101;VPS37B |
| DNA Metabolic Process (GO:0006259) | 0.009828962 | 127.7488696 | DDB1;POLB;ERCC1;GTF2H1 |
| DNA Repair (GO:0006281) | 0.009828962 | 125.7920135 | DDB1;POLB;ERCC1;GTF2H1 |
| Positive Regulation Of Viral Life Cycle (GO:1903902) | 0.009828962 | 822.4999723 | TSG101;VPS37B |
| Late Endosome To Vacuole Transport Via Multivesicular Body Sorting Pathway (GO:0032511) | 0.010919503 | 692.6068823 | TSG101;VPS37B |
| Establishment Of Protein Localization To Vacuole (GO:0072666) | 0.011046469 | 624.7099397 | TSG101;VPS37B |
| Multivesicular Body Assembly (GO:0036258) | 0.011046469 | 567.7192041 | TSG101;VPS37B |
| Multivesicular Body Organization (GO:0036257) | 0.011046469 | 542.5477633 | TSG101;VPS37B |
| Ubiquitin-Dependent Protein Catabolic Process Via The Multivesicular Body Sorting Pathway (GO:0043162) | 0.01115115 | 477.6310459 | TSG101;VPS37B |
| Positive Regulation Of DNA Binding (GO:0043388) | 0.01115115 | 477.6310459 | EDF1;NME1 |
| DNA Damage Response (GO:0006974) | 0.01268379 | 82.556904 | DDB1;POLB;ERCC1;GTF2H1 |
| Membrane Fission (GO:0090148) | 0.01268379 | 395.5173965 | TSG101;VPS37B |
| Double-Strand Break Repair Via Nonhomologous End Joining (GO:0006303) | 0.013780891 | 357.2694654 | POLB;ERCC1 |

| | | | |
|---|---|---|---|
| Regulation Of Epidermal Growth Factor Receptor Signaling Pathway (GO:0042058) | 0.015449221 | 306.192709 | RALA;TSG101 |
| Transcription By RNA Polymerase II (GO:0006366) | 0.015449221 | 111.5694469 | ADRM1;GTF2H1;GTF2F1 |
| Regulation Of DNA Binding (GO:0051101) | 0.015724199 | 289.1798989 | EDF1;NME1 |
| Cellular Response To UV (GO:0034644) | 0.039858294 | 147.0511209 | DDB1;ERCC1 |
| Positive Regulation Of Binding (GO:0051099) | 0.047284586 | 120.9673663 | EDF1;NME1 |
| Positive Regulation Of Cellular Component Biogenesis (GO:0044089) | 0.047284586 | 117.370444 | TSG101;ERCC1 |
| Intracellular Protein Transport (GO:0006886) | 0.047284586 | 50.67678819 | TSG101;VPS37B;STX3 |
| Nucleotide-Excision Repair, DNA Gap Filling (GO:0006297) | 0.047284586 | 1237.703279 | POLB |
| Regulation Of Relaxation Of Cardiac Muscle (GO:1901897) | 0.047284586 | 1237.703279 | PDE4B |
| Regulation Of Extracellular Exosome Assembly (GO:1903551) | 0.047284586 | 1237.703279 | TSG101 |
| Negative Regulation Of Relaxation Of Muscle (GO:1901078) | 0.047284586 | 1237.703279 | PDE4B |
| Macroautophagy (GO:0016236) | 0.047956496 | 100.4180375 | TSG101;VPS37B |
| Negative Regulation Of Telomere Capping (GO:1904354) | 0.051544012 | 955.5316922 | ERCC1 |
| Regulation Of Protein Serine/Threonine Kinase Activity (GO:0071900) | 0.051544012 | 90.48373881 | TSG101;GTF2H1 |
| Ubiquitin-Dependent Protein Catabolic Process (GO:0006511) | 0.052101696 | 41.8763589 | DDB1;TSG101;VPS37B |
| Positive Regulation Of Gluconeogenesis (GO:0045722) | 0.052101696 | 771.8855079 | DDB1 |
| Cellular Response To Epinephrine Stimulus (GO:0071872) | 0.052101696 | 771.8855079 | PDE4B |

| | | | |
|---|---|---|---|
| Response To Epinephrine (GO:0071871) | 0.052101696 | 771.8855079 | PDE4B |
| AMP Biosynthetic Process (GO:0006167) | 0.057031714 | 643.5130349 | APRT |
| Synaptic Vesicle Fusion To Presynaptic Active Zone Membrane (GO:0031629) | 0.057031714 | 549.1072571 | STX3 |
| Regulation Of High Voltage-Gated Calcium Channel Activity (GO:1901841) | 0.057031714 | 549.1072571 | PDE4B |
| Regulation Of Actin Filament-Based Movement (GO:1903115) | 0.057031714 | 549.1072571 | PDE4B |
| Neurotransmitter Receptor Transport To Postsynaptic Membrane (GO:0098969) | 0.057031714 | 549.1072571 | STX3 |
| T-Circle Formation (GO:0090656) | 0.057031714 | 476.9942957 | ERCC1 |
| Formation Of Extrachromosomal Circular DNA (GO:0001325) | 0.057031714 | 476.9942957 | ERCC1 |
| Telomere Maintenance Via Telomere Trimming (GO:0090737) | 0.057031714 | 476.9942957 | ERCC1 |
| Copper Ion Transport (GO:0006825) | 0.057031714 | 476.9942957 | COX17 |
| Nucleoside Triphosphate Metabolic Process (GO:0009141) | 0.057031714 | 476.9942957 | ITPA |
| Nucleoside Phosphate Catabolic Process (GO:1901292) | 0.059994969 | 420.2605662 | ITPA |
| Negative Regulation Of Epidermal Growth Factor-Activated Receptor Activity (GO:0007175) | 0.059994969 | 374.5602037 | TSG101 |
| Pyrimidine Deoxyribonucleotide Catabolic Process (GO:0009223) | 0.059994969 | 374.5602037 | NT5C |
| Positive Regulation Of Protein Localization To Cell Surface (GO:2000010) | 0.059994969 | 374.5602037 | STX3 |

| | | | |
|---|---|---|---|
| Base-Excision Repair, Gap-Filling (GO:0006287) | 0.059994969 | 374.5602037 | POLB |
| Endothelium Development (GO:0003158) | 0.059994969 | 374.5602037 | EDF1 |
| Extracellular Transport (GO:0006858) | 0.060221136 | 337.0293803 | TSG101 |
| Endothelial Cell Differentiation (GO:0045446) | 0.060221136 | 337.0293803 | EDF1 |
| Membrane Organization (GO:0061024) | 0.060221136 | 50.96332839 | TSG101;VPS37B |
| Double-Strand Break Repair (GO:0006302) | 0.060221136 | 50.51872529 | POLB;ERCC1 |
| Synaptic Vesicle Membrane Organization (GO:0048499) | 0.060221136 | 305.7071273 | STX3 |
| Mitotic Recombination (GO:0006312) | 0.060221136 | 305.7071273 | ERCC1 |
| Positive Regulation Of Exosomal Secretion (GO:1903543) | 0.060221136 | 305.7071273 | TSG101 |
| Positive Regulation Of Cell Population Proliferation (GO:0008284) | 0.060221136 | 26.79901757 | COX17;STX3;NME1 |
| Positive Regulation Of Glucose Metabolic Process (GO:0010907) | 0.062185083 | 279.2073187 | DDB1 |
| Negative Regulation Of Telomere Maintenance (GO:0032205) | 0.062185083 | 279.2073187 | ERCC1 |
| Protein Localization To Mitochondrion (GO:0070585) | 0.065191082 | 256.5235419 | RALA |
| Transcription By RNA Polymerase I (GO:0006360) | 0.068094345 | 236.9081425 | GTF2H1 |
| Regulation Of Exosomal Secretion (GO:1903541) | 0.06921749 | 219.7945772 | TSG101 |
| Negative Regulation Of ERBB Signaling Pathway (GO:1901185) | 0.06921749 | 219.7945772 | TSG101 |
| DNA-templated Transcription (GO:0006351) | 0.06921749 | 39.29815577 | GTF2H1;GTF2F1 |
| Cellular Response To Catecholamine Stimulus (GO:0071870) | 0.06921749 | 204.7460437 | PDE4B |

| | | | |
|---|---|---|---|
| Positive Regulation Of Mitochondrial Fission (GO:0090141) | 0.06921749 | 204.7460437 | RALA |
| Vesicle Docking (GO:0048278) | 0.06921749 | 204.7460437 | STX3 |
| Mitochondrial Cytochrome C Oxidase Assembly (GO:0033617) | 0.07314193 | 168.906925 | COX17 |
| Negative Regulation Of Epidermal Growth Factor Receptor Signaling Pathway (GO:0042059) | 0.07314193 | 159.3237838 | TSG101 |
| Negative Regulation Of Protein Tyrosine Kinase Activity (GO:0061099) | 0.07314193 | 159.3237838 | TSG101 |
| Positive Regulation Of ERBB Signaling Pathway (GO:1901186) | 0.07314193 | 150.6523119 | RALA |
| Positive Regulation Of Interleukin-2 Production (GO:0032743) | 0.07314193 | 142.7724214 | PDE4B |
| Mitotic Cell Cycle Checkpoint Signaling (GO:0007093) | 0.07314193 | 142.7724214 | ZWINT |
| Negative Regulation Of Signaling Receptor Activity (GO:2000272) | 0.07314193 | 142.7724214 | TSG101 |
| Regulation Of Cell Cycle Phase Transition (GO:1901987) | 0.07314193 | 142.7724214 | DDB1 |
| Spindle Assembly Checkpoint Signaling (GO:0071173) | 0.07314193 | 135.5839309 | ZWINT |
| Organelle Fusion (GO:0048284) | 0.07314193 | 135.5839309 | STX3 |
| Organelle Membrane Fusion (GO:0090174) | 0.07314193 | 135.5839309 | STX3 |
| DNA-templated Transcription Elongation (GO:0006354) | 0.07314193 | 135.5839309 | GTF2F1 |
| Regulation Of Epidermal Growth Factor-Activated Receptor Activity (GO:0007176) | 0.07314193 | 135.5839309 | TSG101 |
| Mitotic Spindle Assembly Checkpoint Signaling (GO:0007094) | 0.07314193 | 135.5839309 | ZWINT |
| Mitotic Spindle Checkpoint Signaling (GO:0071174) | 0.07314193 | 135.5839309 | ZWINT |

| | | | |
|---|---|---|---|
| Cellular Response To Xenobiotic Stimulus (GO:0071466) | 0.07314193 | 135.5839309 | PDE4B |
| Respiratory Chain Complex IV Assembly (GO:0008535) | 0.07314193 | 135.5839309 | COX17 |
| Regulation Of Cardiac Muscle Cell Contraction (GO:0086004) | 0.07314193 | 135.5839309 | PDE4B |
| Metallo-Sulfur Cluster Assembly (GO:0031163) | 0.07314193 | 129.0026369 | BOLA2 |
| Regulation Of Mitochondrial Fission (GO:0090140) | 0.07314193 | 129.0026369 | RALA |
| Iron-Sulfur Cluster Assembly (GO:0016226) | 0.07314193 | 129.0026369 | BOLA2 |
| Positive Regulation Of Filopodium Assembly (GO:0051491) | 0.07314193 | 129.0026369 | RALA |
| Regulation Of Chemotaxis (GO:0050920) | 0.07314193 | 122.9572755 | STX3 |
| Membrane Fusion (GO:0061025) | 0.07314193 | 122.9572755 | STX3 |
| Transcription Elongation By RNA Polymerase II (GO:0006368) | 0.07314193 | 122.9572755 | GTF2F1 |
| Negative Regulation Of Mitotic Metaphase/Anaphase Transition (GO:0045841) | 0.07314193 | 122.9572755 | ZWINT |
| Regulation Of Voltage-Gated Calcium Channel Activity (GO:1901385) | 0.074120539 | 117.3871491 | PDE4B |
| Positive Regulation Of Viral Genome Replication (GO:0045070) | 0.074120539 | 117.3871491 | DDB1 |
| Positive Regulation Of Locomotion (GO:0040017) | 0.075838036 | 112.2402538 | STX3 |
| Regulation Of Protein Localization To Cell Surface (GO:2000008) | 0.076726534 | 107.4717898 | STX3 |
| Regulation Of Gluconeogenesis (GO:0006111) | 0.076726534 | 107.4717898 | DDB1 |
| Transition Metal Ion Transport (GO:0000041) | 0.078360587 | 103.0429662 | COX17 |
| Positive Regulation Of Ion Transmembrane | 0.079167774 | 98.92003497 | COX17 |

| | | | |
|---|---|---|---|
| Transporter Activity (GO:0032414) | | | |
| Positive Regulation Of Epidermal Growth Factor Receptor Signaling Pathway (GO:0045742) | 0.079167774 | 98.92003497 | RALA |
| Protein Targeting To Vacuole (GO:0006623) | 0.079172515 | 95.07350427 | VPS37B |
| Negative Regulation Of Mitotic Cell Cycle (GO:0045930) | 0.079172515 | 95.07350427 | ZWINT |
| Sister Chromatid Segregation (GO:0000819) | 0.079172515 | 95.07350427 | ZWINT |
| Regulation Of Cardiac Muscle Contraction (GO:0055117) | 0.08068274 | 91.47749401 | PDE4B |
| Base-Excision Repair (GO:0006284) | 0.082162112 | 88.10920426 | POLB |
| Positive Regulation Of Exocytosis (GO:0045921) | 0.083611496 | 84.94847445 | TSG101 |
| Regulation Of Telomere Maintenance (GO:0032204) | 0.08503173 | 81.97741569 | ERCC1 |
| Synaptic Vesicle Exocytosis (GO:0016079) | 0.089935656 | 74.05131012 | STX3 |
| Positive Regulation Of Viral Process (GO:0048524) | 0.089935656 | 74.05131012 | DDB1 |
| DNA Recombination (GO:0006310) | 0.091251559 | 71.69565287 | ERCC1 |
| Regulation Of Primary Metabolic Process (GO:0080090) | 0.091380865 | 69.46503882 | EDF1 |
| Regulation Of Filopodium Assembly (GO:0051489) | 0.091380865 | 69.46503882 | RALA |
| Establishment Of Protein Localization To Mitochondrion (GO:0072655) | 0.091380865 | 67.35016648 | RALA |
| Positive Regulation Of Chemotaxis (GO:0050921) | 0.091380865 | 67.35016648 | STX3 |
| Autophagosome Maturation (GO:0097352) | 0.091380865 | 67.35016648 | TSG101 |
| Organelle Assembly (GO:0070925) | 0.091825739 | 18.6155061 | TSG101;VPS37B |
| Mitotic Nuclear Division (GO:0140014) | 0.091825739 | 65.34261802 | ZWINT |

| | | | |
|---|---|---|---|
| Vacuolar Transport (GO:0007034) | 0.093028961 | 63.43475777 | VPS37B |
| Regulation Of Cell Population Proliferation (GO:0042127) | 0.093431473 | 11.90016522 | COX17;STX3;NME1 |
| Regulation Of Interleukin-2 Production (GO:0032663) | 0.093431473 | 61.61964439 | PDE4B |
| Negative Regulation Of DNA Metabolic Process (GO:0051053) | 0.096508124 | 58.24291595 | ERCC1 |
| Intracellular Iron Ion Homeostasis (GO:0006879) | 0.097157044 | 56.67025 | BOLA2 |
| Receptor Internalization (GO:0031623) | 0.097157044 | 55.16811991 | RALA |
| Positive Regulation Of Protein Localization To Cell Periphery (GO:1904377) | 0.097157044 | 55.16811991 | STX3 |
| Positive Regulation Of Protein Localization To Plasma Membrane (GO:1903078) | 0.097157044 | 55.16811991 | STX3 |
| Positive Regulation Of Telomere Maintenance (GO:0032206) | 0.098230839 | 53.73208629 | ERCC1 |
| Regulation Of Cyclin-Dependent Protein Kinase Activity (GO:1904029) | 0.099285935 | 52.35806749 | GTF2H1 |
| Regulation Of Exocytosis (GO:0017157) | 0.099551056 | 51.04230466 | RALA |
| Regulation Of Lipid Metabolic Process (GO:0019216) | 0.099551056 | 51.04230466 | EDF1 |
| Endosome To Lysosome Transport (GO:0008333) | 0.104116633 | 47.4111797 | TSG101 |
| Positive Regulation Of Mitochondrion Organization (GO:0010822) | 0.104295764 | 46.29629512 | RALA |
| Positive Regulation Of Type II Interferon Production (GO:0032729) | 0.104295764 | 46.29629512 | PDE4B |
| DNA-templated Transcription Initiation (GO:0006352) | 0.105247387 | 45.22474372 | GTF2F1 |

| | | | |
|---|---|---|---|
| Protein-Containing Complex Disassembly (GO:0032984) | 0.109608341 | 42.24725438 | TSG101 |
| Regulation Of Secretion By Cell (GO:1903530) | 0.11219529 | 40.4397246 | RALA |
| Cellular Response To Light Stimulus (GO:0071482) | 0.11305775 | 39.58384099 | DDB1 |
| Regulation Of Viral Genome Replication (GO:0045069) | 0.115571656 | 37.96004303 | DDB1 |
| Establishment Of Protein Localization To Membrane (GO:0090150) | 0.116366143 | 36.44426972 | VPS37B |
| DNA-templated DNA Replication (GO:0006261) | 0.116366143 | 36.44426972 | POLB |
| Regulation Of Actin Filament-Based Process (GO:0032970) | 0.116366143 | 35.72373458 | RALA |
| Neutrophil Chemotaxis (GO:0030593) | 0.116366143 | 35.72373458 | PDE4B |
| Vesicle-Mediated Transport (GO:0016192) | 0.116366143 | 12.4390663 | TSG101;STX3 |
| Negative Regulation Of Protein Binding (GO:0032091) | 0.117124569 | 35.02656814 | GTF2F1 |
| Protein Targeting To Membrane (GO:0006612) | 0.117843636 | 34.35170636 | VPS37B |
| DNA Replication (GO:0006260) | 0.117843636 | 33.69814743 | POLB |
| Granulocyte Chemotaxis (GO:0071621) | 0.117843636 | 33.69814743 | PDE4B |
| Transcription Initiation At RNA Polymerase II Promoter (GO:0006367) | 0.121667734 | 31.8561127 | GTF2F1 |
| Neutrophil Migration (GO:1990266) | 0.122033631 | 31.27884428 | PDE4B |
| Proteolysis Involved In Protein Catabolic Process (GO:0051603) | 0.122033631 | 30.71866043 | DDB1 |
| Regulation Of Cyclin-Dependent Protein Serine/Threonine Kinase Activity (GO:0000079) | 0.122033631 | 30.1748515 | GTF2H1 |
| Positive Regulation Of Protein Localization To Membrane (GO:1905477) | 0.122033631 | 30.1748515 | STX3 |

| | | | |
|---|---|---|---|
| Positive Regulation Of Catabolic Process (GO:0009896) | 0.122033631 | 30.1748515 | DDB1 |
| Regulation Of Protein Localization To Plasma Membrane (GO:1903076) | 0.122033631 | 29.64674578 | STX3 |
| Regulation Of Organelle Assembly (GO:1902115) | 0.122033631 | 29.64674578 | TSG101 |
| Establishment Of Protein Localization To Organelle (GO:0072594) | 0.126691431 | 27.2206113 | RALA |
| Positive Regulation Of Protein Localization (GO:1903829) | 0.126691431 | 27.2206113 | STX3 |
| Exocytosis (GO:0006887) | 0.126691431 | 26.77445194 | STX3 |
| Regulation Of Mitotic Cell Cycle Phase Transition (GO:1901990) | 0.126691431 | 26.77445194 | DDB1 |
| Negative Regulation Of Binding (GO:0051100) | 0.126691431 | 26.77445194 | GTF2F1 |
| Regulation Of Type II Interferon Production (GO:0032649) | 0.127119822 | 26.34016926 | PDE4B |
| Positive Regulation Of Plasma Membrane Bounded Cell Projection Assembly (GO:0120034) | 0.127119822 | 25.91731986 | RALA |
| Positive Regulation Of Cell Adhesion (GO:0045785) | 0.127119822 | 25.91731986 | STX3 |
| Regulation Of Vesicle-Mediated Transport (GO:0060627) | 0.127714184 | 25.50548161 | RALA |
| Mitochondrial Respiratory Chain Complex Assembly (GO:0033108) | 0.128105427 | 25.10425243 | COX17 |
| Regulation Of Epithelial Cell Proliferation (GO:0050678) | 0.128105427 | 24.71324913 | NME1 |
| Regulation Of Actin Cytoskeleton Organization (GO:0032956) | 0.128105427 | 24.71324913 | RALA |
| Positive Regulation Of Protein Catabolic Process (GO:0045732) | 0.128675546 | 24.3321063 | DDB1 |
| T Cell Receptor Signaling Pathway (GO:0050852) | 0.131880069 | 23.24443303 | PDE4B |
| Response To UV (GO:0009411) | 0.137647602 | 21.59922754 | DDB1 |

| | | | |
|---|---|---|---|
| Protein Catabolic Process (GO:0030163) | 0.139435059 | 20.99351209 | DDB1 |
| Lysosomal Transport (GO:0007041) | 0.14248021 | 20.13519876 | TSG101 |
| Mitotic Sister Chromatid Segregation (GO:0000070) | 0.149287348 | 18.57990663 | ZWINT |
| Small GTPase Mediated Signal Transduction (GO:0007264) | 0.149689184 | 18.33955918 | RALA |
| Regulation Of Cytoskeleton Organization (GO:0051493) | 0.150475214 | 17.87363593 | RALA |
| Regulation Of MAP Kinase Activity (GO:0043405) | 0.150475214 | 17.87363593 | TSG101 |
| Protein Maturation (GO:0051604) | 0.1524634 | 17.20960125 | BOLA2 |
| Cellular Response To Molecule Of Bacterial Origin (GO:0071219) | 0.1524634 | 17.20960125 | PDE4B |
| Regulation Of Protein Binding (GO:0043393) | 0.152828529 | 16.99703051 | GTF2F1 |
| Positive Regulation Of Nucleic Acid-Templated Transcription (GO:1903508) | 0.154509723 | 7.300904301 | EDF1;GTF2H1 |
| Regulation Of Protein Catabolic Process (GO:0042176) | 0.155084037 | 16.18746001 | DDB1 |
| Receptor-Mediated Endocytosis (GO:0006898) | 0.155084037 | 16.18746001 | RALA |
| Positive Regulation Of Epithelial Cell Proliferation (GO:0050679) | 0.155420592 | 15.99469964 | NME1 |
| Cellular Response To Lipopolysaccharide (GO:0071222) | 0.155752244 | 15.80558703 | PDE4B |
| Regulation Of Mitotic Cell Cycle (GO:0007346) | 0.156079075 | 15.62002575 | DDB1 |
| Protein Targeting (GO:0006605) | 0.159877476 | 14.91147739 | VPS37B |
| Epithelial Cell Differentiation (GO:0030855) | 0.161605089 | 14.41309002 | EDF1 |
| Plasma Membrane Bounded Cell Projection | 0.161605089 | 14.41309002 | STX3 |

| | | | |
|---|---|---|---|
| Organization (GO:0120036) | | | |
| Antigen Receptor-Mediated Signaling Pathway (GO:0050851) | 0.16255185 | 14.09539523 | PDE4B |
| Positive Regulation Of Cellular Process (GO:0048522) | 0.16255185 | 6.488350861 | COX17;STX3 |
| Ras Protein Signal Transduction (GO:0007265) | 0.171560349 | 12.66171067 | RALA |
| Regulation Of Cell Adhesion (GO:0030155) | 0.171560349 | 12.66171067 | STX3 |
| Dephosphorylation (GO:0016311) | 0.173958479 | 12.27626996 | NT5C |
| Neuron Development (GO:0048666) | 0.17632152 | 11.90902253 | STX3 |
| Positive Regulation Of DNA-templated Transcription (GO:0045893) | 0.176376338 | 4.470830597 | EDF1;GTF2H1;GTF2F1 |
| Positive Regulation Of Response To External Stimulus (GO:0032103) | 0.179877057 | 11.33420999 | STX3 |
| Response To Lipopolysaccharide (GO:0032496) | 0.18320576 | 10.905108 | PDE4B |
| Regulation Of Macromolecule Metabolic Process (GO:0060255) | 0.185434039 | 10.59975087 | STX3 |
| Positive Regulation Of Cellular Biosynthetic Process (GO:0031328) | 0.196953824 | 9.502514245 | DDB1 |
| Phosphate-Containing Compound Metabolic Process (GO:0006796) | 0.206173842 | 8.717239265 | NT5C |
| Intracellular Monoatomic Cation Homeostasis (GO:0030003) | 0.21109372 | 8.293245219 | BOLA2 |
| Modification-Dependent Protein Catabolic Process (GO:0019941) | 0.21109372 | 8.158958784 | DDB1 |
| Neuron Projection Development (GO:0031175) | 0.21109372 | 8.158958784 | STX3 |
| Regulation Of Programmed Cell Death (GO:0043067) | 0.212953766 | 7.963719877 | NME1 |

| | | | |
|---|---|---|---|
| Positive Regulation Of Protein Metabolic Process (GO:0051247) | 0.212953766 | 7.900234874 | DDB1 |
| Proteasomal Protein Catabolic Process (GO:0010498) | 0.234978279 | 6.582174803 | DDB1 |
| Cellular Response To Lipid (GO:0071396) | 0.241363029 | 6.216491122 | PDE4B |
| Positive Regulation Of Developmental Process (GO:0051094) | 0.244847774 | 6.003345598 | RALA |
| Protein Transport (GO:0015031) | 0.314393117 | 3.681140559 | STX3 |
| Proteasome-Mediated Ubiquitin-Dependent Protein Catabolic Process (GO:0043161) | 0.317262697 | 3.563535617 | DDB1 |
| Positive Regulation Of Cytokine Production (GO:0001819) | 0.317262697 | 3.544468893 | PDE4B |
| Protein-Containing Complex Assembly (GO:0065003) | 0.322366847 | 3.39714447 | ADRM1 |
| Protein Localization (GO:0008104) | 0.33943144 | 3.01987377 | STX3 |
| Protein Modification By Small Protein Conjugation (GO:0032446) | 0.348097493 | 2.832873084 | DDB1 |
| Negative Regulation Of Cell Population Proliferation (GO:0008285) | 0.358077807 | 2.637069143 | NME1 |
| Regulation Of Gene Expression (GO:0010468) | 0.367271488 | 1.739539663 | EDF1;STX3 |
| Cellular Response To Oxygen-Containing Compound (GO:1901701) | 0.375004379 | 2.330271568 | PDE4B |
| Protein Ubiquitination (GO:0016567) | 0.393557235 | 2.062789055 | DDB1 |
| Regulation Of Nucleic Acid-Templated Transcription (GO:1903506) | 0.404416617 | 1.913081124 | EDF1 |
| Negative Regulation Of Cellular Process (GO:0048523) | 0.458884612 | 1.376580303 | NME1 |

| | | | |
|---|---|---|---|
| Regulation Of Apoptotic Process (GO:0042981) | 0.553531755 | 0.788637622 | NME1 |
| Regulation Of DNA-templated Transcription (GO:0006355) | 0.644273493 | 0.421918243 | EDF1;GTF2H1 |
| Positive Regulation Of Transcription By RNA Polymerase II (GO:0045944) | 0.655550124 | 0.412961224 | GTF2F1 |
| Regulation Of Transcription By RNA Polymerase II (GO:0006357) | 0.904961211 | 0.042113474 | GTF2F1 |

**Table S8. Gene Set Enrichment Analysis on significantly deregulated DNA Repair genes in PK4A cells grown on CDM vs TCP.**

| **W4** | | | **W6** | | | **W9** | | |
|---|---|---|---|---|---|---|---|---|
| | **Symbol** | **logFC** | | **Symbol** | **logFC** | | **Symbol** | **logFC** |
| UP | Pappa | 0.743000819 | UP | Bmper | 0.927574814 | UP | Bmper | 1.156382183 |
| | Serpinf1 | 1.125973328 | | Pappa | 0.95301536 | | Pappa | 2.299483936 |
| | Lamc2 | 2.107126513 | | Serpinf1 | 1.138026338 | | Acan | 0.777061032 |
| | Sema4d | 0.660147728 | | Lamc2 | 2.432821332 | | Serpinf1 | 2.192689193 |
| | Col1a1 | 1.733939679 | | Fcna | 0.774299959 | | Lamc2 | 4.114423335 |
| | Cd109 | 0.834220483 | | Col1a1 | 1.798079964 | | Fcna | 1.174011325 |
| | Clec4d | 1.05203291 | | Cd109 | 1.097371036 | | Col1a1 | 3.007867928 |
| | Bgn | 1.419478408 | | Plxna4 | 0.603955873 | | Cd109 | 2.855908346 |
| | Col3a1 | 1.027220178 | | Clec4d | 1.37525797 | | Tgfb2 | 1.011843352 |
| | Pdgfc | 0.810627357 | | Bgn | 1.064247116 | | Plxna4 | 1.478912111 |
| | Tnc | 4.19755889 | | Col3a1 | 1.152614075 | | Clec4d | 2.311475844 |
| | Mfge8 | 0.842311197 | | Pdgfc | 1.435438903 | | Bgn | 1.947087319 |
| | Inhba | 2.356812107 | | Tnc | 3.994840049 | | Col3a1 | 2.042199243 |
| | Ccl7 | 0.978817823 | | Angpt2 | 0.635168162 | | Pdgfc | 2.632197058 |
| | Il18 | 0.836546453 | | Inhba | 2.880410515 | | Tnc | 6.422815299 |
| | Igsf10 | 0.721462147 | | Ccl7 | 1.71478869 | | Angpt2 | 1.197611119 |
| | Serpinb9b | 0.868943952 | | Il18 | 0.694559032 | | Anxa7 | 0.632309715 |
| | Colec12 | 0.70526922 | | Igsf10 | 0.771258919 | | Sema3f | 1.325367521 |
| | Mmp19 | 1.045578877 | | Serpinb9b | 1.103149899 | | Lgals1 | 0.84185958 |
| | Adam12 | 1.501700051 | | Colec12 | 0.85650314 | | Mfge8 | 0.92288401 |
| | Lgals3 | 1.190136627 | | Col6a1 | 0.658335769 | | Inhba | 4.162243536 |
| | Ndnf | 0.730634166 | | Mmp19 | 1.280046125 | | Ccl7 | 2.598462562 |
| | Timp1 | 4.401990635 | | Adam12 | 1.379831243 | | Il18 | 2.271353414 |

| | | | | | | | | |
|---|---|---|---|---|---|---|---|---|
| | Adam19 | 1.385858662 | | Lgals3 | 1.36908342 | | Igsf10 | 1.299340117 |
| | Sema4b | 0.59167091 | | Ndnf | 0.778081864 | | Serpinb9b | 2.336203748 |
| | Col8a1 | 2.561321222 | | Timp1 | 4.397034107 | | Colec12 | 1.883822412 |
| | Slpi | 0.748943428 | | Adam19 | 0.945530057 | | Col6a1 | 1.571241651 |
| | Csf1 | 0.725814491 | | Sema4b | 0.609171244 | | Adamts7 | 1.136931246 |
| | Slit2 | 1.474009582 | | Col8a1 | 2.714257896 | | Mmp19 | 2.5701386 |
| | Edil3 | 0.660329581 | | Slpi | 0.628171996 | | Adam12 | 2.768188085 |
| | Mgp | 1.449336141 | | Csf1 | 0.684221591 | | Lgals3 | 2.629019533 |
| | Fbn1 | 0.996360129 | | Slit2 | 1.687279879 | | Ndnf | 1.381620012 |
| | Cxcl10 | 1.569904662 | | Vwa5a | 0.73511788 | | Cstb | 0.752534302 |
| | Nid1 | 0.80031577 | | Edil3 | 1.196689734 | | Timp1 | 5.972244897 |
| | Egfl7 | 0.6558699 | | Mgp | 1.333106366 | | Adam19 | 2.447109962 |
| | Masp1 | 0.814670527 | | Fbn1 | 1.137778188 | | Ctsb | 0.930058549 |
| | Serpinb2 | 0.705094231 | | Mmp3 | 0.840612655 | | Sema4b | 1.496427019 |
| | Anxa1 | 1.672235628 | | Nid1 | 0.594291129 | | Col8a1 | 4.141168194 |
| | S100a11 | 1.236599965 | | Lama5 | 0.804004109 | | Slpi | 0.664811186 |
| | Crim1 | 0.636512505 | | Spon2 | 0.894330279 | | Csf1 | 1.370802274 |
| | Fn1 | 2.053001998 | | Masp1 | 1.215881084 | | Slit2 | 3.048517231 |
| | S100a10 | 1.518692006 | | Serpinb2 | 0.77216176 | | Ccbe1 | 0.816732805 |
| | Serping1 | 1.115371661 | | Anxa1 | 2.069143271 | | Vegfc | 1.41478228 |
| | Adamts2 | 1.239958562 | | S100a11 | 1.158319742 | | Vwa5a | 1.321328138 |
| | Sned1 | 0.700078141 | | Crim1 | 0.658243437 | | Edil3 | 2.800522902 |
| | Col6a3 | 0.875658146 | | Fn1 | 1.95369889 | | Mgp | 2.176642269 |
| | Sfrp1 | 3.298101658 | | S100a10 | 1.476366904 | | Fbn1 | 2.211012262 |
| | Col12a1 | 3.313317655 | | Serpinb6a | 0.780083746 | | Mmp3 | 0.753244757 |

| | | | | | | | | |
|---|---|---|---|---|---|---|---|---|
| | Gpc6 | 0.86323235 | | Serping1 | 0.848336609 | | Podnl1 | 0.701520048 |
| | Ctsc | 0.833646036 | | Adamts2 | 1.376679781 | | Cxcl10 | 1.543165306 |
| | Sema6a | 0.635792106 | | Col6a3 | 1.011385661 | | Nid1 | 1.720060678 |
| | Spp1 | 1.379959182 | | Sfrp1 | 2.816548164 | | Lama5 | 2.279795839 |
| | Ctsz | 0.865591215 | | Col12a1 | 3.232067111 | | Lamc1 | 1.221993105 |
| | Mmp23 | 0.793643805 | | Gpc6 | 0.752845785 | | Spon2 | 1.341825031 |
| | P4ha3 | 2.158291793 | | Ctsc | 0.873093313 | | Egfl7 | 0.708135011 |
| | Col15a1 | 0.951950074 | | Adam9 | 0.716140156 | | Masp1 | 3.131314871 |
| | Hspg2 | 0.698434244 | | Spp1 | 1.949200093 | | Serpinb2 | 1.858707396 |
| | Pdgfb | 0.754989016 | | Fgf7 | 0.583947089 | | Anxa1 | 3.756867436 |
| | Sema3d | 0.60250677 | | Ctsz | 0.678820543 | | S100a11 | 1.965337285 |
| | Thbs2 | 2.545778906 | | Mmp23 | 0.75627428 | | Crim1 | 1.51786038 |
| | Aebp1 | 1.323856062 | | P4ha3 | 1.949307037 | | Fn1 | 2.990440792 |
| | Lox | 2.243398028 | | Col15a1 | 0.793331755 | | S100a10 | 2.831486111 |
| | Adamts6 | 1.285780125 | | Sema3d | 0.910637492 | | Crlf1 | 0.920370696 |
| | Cx3cl1 | 0.649299497 | | Thbs2 | 2.481488125 | | Nrg1 | 1.3434766 |
| | Ctsh | 1.660583064 | | Aebp1 | 1.278960089 | | Emilin1 | 0.860133137 |
| | Col1a2 | 1.604864654 | | Lox | 2.253564447 | | Col5a1 | 1.0771754 |
| | Efemp2 | 1.337193474 | | Adamts6 | 1.06408673 | | Serpinb6a | 1.206622885 |
| | Ctso | 0.808572082 | | Cx3cl1 | 0.776635302 | | Serping1 | 1.54169387 |
| | Il34 | 0.665046185 | | Ctsh | 1.603339751 | | Serpinb8 | 0.900560798 |
| | Hpse | 1.93429984 | | Col1a2 | 1.649651663 | | Adamtsl4 | 1.575636517 |
| | Sema5a | 0.719943197 | | Efemp2 | 1.518236114 | | Adamts2 | 2.617824767 |
| | Npnt | 2.328570739 | | Ctso | 0.607283209 | | Sned1 | 0.886821085 |
| | Ltbp1 | 0.600987721 | | Hpse | 0.981240579 | | Col6a3 | 2.252727219 |

| | | | | | | | | |
|---|---|---|---|---|---|---|---|---|
| | Adamts9 | 0.648219706 | | Sema5a | 0.75927194 | | Pik3ip1 | 0.816828176 |
| | Loxl1 | 1.056345126 | | Npnt | 2.773998674 | | Sfrp1 | 3.54861051 |
| | Loxl2 | 0.658886392 | | Ltbp1 | 0.963242706 | | Col12a1 | 5.36385876 |
| | Prl2c3 | 1.837122074 | | Loxl1 | 1.221155883 | | Gpc6 | 1.751454244 |
| | Ltbp2 | 1.25668972 | | Sulf2 | 0.834953068 | | Ctsc | 1.388926972 |
| | Anxa2 | 2.326620666 | | Loxl2 | 1.319021232 | | Adam9 | 1.301115814 |
| | Pcolce | 0.773848474 | | Prl2c3 | 2.932643728 | | Sema6a | 1.061975615 |
| | Timp3 | 1.654316859 | | Ltbp2 | 1.440948907 | | Megf10 | 1.785550216 |
| | Ccl2 | 0.959896056 | | Anxa2 | 2.339176376 | | Gdf11 | 0.913112632 |
| | Sulf1 | 1.469422252 | | Pcolce | 0.756826109 | | Spp1 | 2.419974693 |
| | Slit3 | 0.666621259 | | Timp3 | 1.326884848 | | Ism1 | 1.135201166 |
| | Ctse | 4.704456137 | | Ccl2 | 1.313147832 | | Fgf7 | 0.631691265 |
| | Mmp2 | 1.614131234 | | Ptn | 0.760026895 | | Pdgfd | 0.634193376 |
| | Cthrc1 | 0.927996705 | | Inhbb | 0.736190544 | | Ctsz | 1.375253381 |
| | Ngf | 1.749103576 | | Sulf1 | 1.236384182 | | Mmp23 | 1.456123607 |
| | Lgi2 | 0.778040812 | | Slit3 | 0.866662989 | | P4ha3 | 3.381654126 |
| | C1qtnf1 | 1.044362063 | | Ctse | 3.079185766 | | Col15a1 | 1.862466278 |
| | Loxl3 | 0.668313256 | | Sema3e | 0.719765669 | | Hspg2 | 1.532389452 |
| DOWN | Itih2 | -0.768159391 | | Mmp2 | 1.648523275 | | Pdgfb | 1.344268611 |
| | Angpt1 | -0.743301574 | | Cthrc1 | 1.293097833 | | Sema3d | 1.918293912 |
| | Ntn1 | -0.650037349 | | Ngf | 1.762008485 | | Thbs2 | 4.167583922 |
| | Cela1 | -0.619308782 | | Lgi2 | 0.740850017 | | Aebp1 | 2.126293025 |
| | | | | Esm1 | 0.769102375 | | Lox | 4.191730888 |

| | | | | | | | | |
|---|---|---|---|---|---|---|---|---|
| | | | | C1qtnf1 | 0.854934583 | | Adamts6 | 2.577270759 |
| | | | | Loxl3 | 1.314227125 | | Htra3 | 1.369878879 |
| | | | DOWN | Ntn1 | -0.927966507 | | Cx3cl1 | 2.04777007 |
| | | | | Smoc1 | -0.71745036 | | Ctsh | 2.571024739 |
| | | | | | | | Col1a2 | 2.829207505 |
| | | | | | | | Efemp2 | 2.480675542 |
| | | | | | | | Ctso | 1.255203454 |
| | | | | | | | Sdc4 | 1.188191064 |
| | | | | | | | Il34 | 1.864643563 |
| | | | | | | | Hpse | 1.971103858 |
| | | | | | | | Matn2 | 0.602866251 |
| | | | | | | | Nid2 | 0.803951487 |
| | | | | | | | Sema5a | 1.503163363 |
| | | | | | | | Clec2d | 0.673308467 |
| | | | | | | | Npnt | 4.345711457 |
| | | | | | | | Ltbp1 | 2.095995763 |
| | | | | | | | Adamts9 | 1.003885846 |
| | | | | | | | Adamts14 | 0.852168001 |
| | | | | | | | Loxl1 | 2.221342857 |
| | | | | | | | Sulf2 | 2.109547809 |
| | | | | | | | Loxl2 | 2.415642502 |
| | | | | | | | Sema4f | 0.66429971 |
| | | | | | | | Col6a2 | 1.456460256 |
| | | | | | | | Prl2c3 | 4.43938572 |

| | | | | | | | | |
|---|---|---|---|---|---|---|---|---|
| | | | | | | | Lamb3 | 0.77761496 |
| | | | | | | | Ltbp2 | 3.242791798 |
| | | | | | | | Lamb2 | 0.988400398 |
| | | | | | | | Anxa2 | 3.777114566 |
| | | | | | | | Pcolce | 1.35624643 |
| | | | | | | | Timp3 | 2.053126887 |
| | | | | | | | Ccl2 | 2.370439196 |
| | | | | | | | Inhbb | 1.740975024 |
| | | | | | | | Sulf1 | 2.472666998 |
| | | | | | | | Slit3 | 1.590601064 |
| | | | | | | | Thbs3 | 0.908296454 |
| | | | | | | | Ctse | 6.488579944 |
| | | | | | | | Sema3e | 1.54943102 |
| | | | | | | | Mmp2 | 2.975202362 |
| | | | | | | | Cthrc1 | 1.961446232 |
| | | | | | | | Wnt7a | 1.643361267 |
| | | | | | | | Ngf | 2.592634688 |
| | | | | | | | Lgi2 | 0.648289038 |
| | | | | | | | C1qtnf1 | 2.108950612 |
| | | | | | | | Loxl3 | 2.775835703 |
| | | | | | | DOWN | Ogfod2 | -0.889450036 |
| | | | | | | | Nrtn | -0.702903084 |
| | | | | | | | Itih2 | -2.851238433 |

| | | | | | | | | |
|---|---|---|---|---|---|---|---|---|
| | | | | | | | Muc1 | -1.055836137 |
| | | | | | | | Spock2 | -0.75413411 |
| | | | | | | | Pcsk6 | -0.721629251 |
| | | | | | | | Ctf1 | -1.08843823 |
| | | | | | | | Ngly1 | -0.810148275 |
| | | | | | | | Serpinb1a | -1.084761027 |
| | | | | | | | Ogn | -1.394792099 |
| | | | | | | | Kazald1 | -0.892799881 |
| | | | | | | | Ntn1 | -2.598804314 |
| | | | | | | | Smoc1 | -1.94429811 |
| | | | | | | | Lman1 | -0.885394198 |
| | | | | | | | Cela1 | -4.948909403 |
| | | | | | | | Ntn3 | -0.738671621 |

**Table S9. Significantly deregulated matrisomal genes in PDAC from 4-, 6-, and 9-week-old mice compared to healthy pancreases from healthy age-matched littermates.**

| **W4** | | | **W6** | | | **W9** | | |
|---|---|---|---|---|---|---|---|---|
| | **Symbol** | **logFC** | | **Symbol** | **logFC** | | **Symbol** | **logFC** |
| UP | Lacc1 | 1.436938 | UP | Lacc1 | 0.998629 | UP | Ak8 | 0.661009 |
| | Adcy7 | 1.463565 | | Adcy7 | 0.854905 | | Atic | 0.684515 |
| | Gucy1b1 | 0.599947 | | Pnp2 | 0.677581 | | Pde7a | 0.836188 |
| | Adcy4 | 0.61728 | | Nme7 | 0.639595 | | Lacc1 | 2.727491 |
| | Pnp2 | 0.711431 | | Npr2 | 0.658176 | | Pde1b | 1.037501 |
| | Npr2 | 0.781326 | | Itpa | 0.582458 | | Adcy7 | 2.657372 |
| | Dck | 0.581883 | | Ak1 | 0.692502 | | Ada | 1.201897 |
| | Papss2 | 1.012507 | | Rrm1 | 1.196043 | | Prune1 | 0.898164 |
| | Rrm1 | 1.107791 | | Gda | 0.999815 | | Adcy2 | 0.664026 |
| | Gda | 0.782451 | | Pde7b | 0.688301 | | Ak5 | 0.81981 |
| | Pde7b | 0.977873 | | Ampd3 | 0.612846 | | Adcy3 | 1.008316 |
| | Ampd3 | 0.594932 | | Nt5e | 2.023862 | | Nt5c2 | 1.098785 |
| | Pde4b | 1.056352 | | Pde5a | 0.649135 | | Pgm2 | 0.934379 |
| | Nt5e | 1.537036 | | Pnp | 0.700063 | | Gucy1b1 | 0.81154 |
| | Pde5a | 0.60393 | DOWN | Impdh1 | -0.58308 | | Adcy8 | 1.151237 |
| | Pnp | 0.625795 | | Ada | -0.75401 | | Adcy4 | 0.857548 |
| DOWN | Guk1 | -0.77031 | | Adk | -0.61989 | | Pnp2 | 1.572687 |
| | Urah | -0.58555 | | Pde8b | -0.60142 | | Enpp1 | 1.030523 |
| | Adk | -0.64504 | | | | | Nme7 | 1.392633 |
| | Pde8b | -0.60445 | | | | | Npr2 | 1.603842 |
| | | | | | | | Pde1a | 0.934915 |
| | | | | | | | Nt5c | 0.81377 |
| | | | | | | | Itpa | 0.824185 |
| | | | | | | | Papss2 | 1.656171 |
| | | | | | | | Enpp3 | 1.599032 |
| | | | | | | | Ak1 | 0.651226 |
| | | | | | | | Rrm1 | 1.905265 |
| | | | | | | | Gda | 1.581458 |
| | | | | | | | Pde7b | 1.349121 |
| | | | | | | | Ampd3 | 1.078291 |
| | | | | | | | Xdh | 1.85836 |
| | | | | | | | Nt5c1a | 1.324159 |
| | | | | | | | Pde10a | 0.930684 |
| | | | | | | | Pde3a | 1.053713 |
| | | | | | | | Pde4b | 1.03453 |
| | | | | | | | Nt5e | 4.006326 |
| | | | | | | | Gmpr2 | 0.689213 |
| | | | | | | | Pde5a | 1.213398 |
| | | | | | | | Aprt | 1.020484 |

| | | | | | | | | |
|---|---|---|---|---|---|---|---|---|
| | | | | | | | Ntpcr | 0.755418 |
| | | | | | | | Gucy2c | 0.714955 |
| | | | | | | | Pnp | 1.49676 |
| | | | | | | | Pde4d | 0.590174 |
| | | | | | | DOWN | Entpd1 | -0.83149 |
| | | | | | | | Nme2 | -0.63253 |
| | | | | | | | Pde4a | -0.79946 |
| | | | | | | | Urah | -1.29557 |
| | | | | | | | Impdh1 | -1.33201 |
| | | | | | | | Entpd4 | -1.02945 |
| | | | | | | | Ak9 | -0.86043 |
| | | | | | | | Hddc3 | -0.90565 |
| | | | | | | | Entpd2 | -0.97556 |
| | | | | | | | Prps2 | -0.81096 |
| | | | | | | | Urad | -0.62887 |
| | | | | | | | Ak3 | -1.16856 |
| | | | | | | | Nme3 | -0.78205 |
| | | | | | | | Adk | -1.70643 |
| | | | | | | | Nme6 | -1.15025 |
| | | | | | | | Pde8b | -2.21204 |
| | | | | | | | Adcy9 | -1.31353 |
| | | | | | | | Guk1 | -1.04229 |

**Table S10. Significantly deregulated purine metabolism genes in PDAC from 4-, 6-, and 9-week-old mice compared to healthy pancreases from healthy age-matched littermates.**

| W4 | | | W6 | | | W9 | | |
|---|---|---|---|---|---|---|---|---|
| | Symbol | logFC | | Symbol | logFC | | Symbol | logFC |
| UP | Pcna | 0.633722 | UP | Pcna | 0.599971 | UP | Dctn4 | 0.758609 |
| | Arl6ip1 | 0.926008 | | Arl6ip1 | 0.783727 | | Pcna | 1.008156 |
| | Hcls1 | 1.477891 | | Hcls1 | 0.854406 | | Lig1 | 0.797052 |
| | Zfp36l2 | 0.948928 | | Zwint | 0.785401 | | Arl6ip1 | 1.156947 |
| | Cstf3 | 0.721587 | | Zfp36l2 | 0.584744 | | Hcls1 | 2.138852 |
| | Npr2 | 0.781326 | | Npr2 | 0.658176 | | Zwint | 1.415476 |
| | Usp11 | 0.711616 | | Itpa | 0.582458 | | Ada | 1.201897 |
| | Pde4b | 1.056352 | | Rpa3 | 0.838498 | | Zfp36l2 | 1.622929 |
| | Trp53 | 1.174352 | | Ercc1 | 0.592893 | | Cstf3 | 1.467369 |
| | Pnp | 0.625795 | | Ak1 | 0.692502 | | Pold4 | 0.659703 |
| DOWN | Guk1 | -0.77031 | | Nudt21 | 0.584932 | | Dgcr8 | 1.144591 |
| | | | | Pola1 | 0.774431 | | Smad5 | 0.819995 |
| | | | | Trp53 | 0.743256 | | Npr2 | 1.603842 |
| | | | | Pnp | 0.700063 | | Ddb1 | 0.698559 |
| | | | DOWN | Ada | -0.75401 | | Rfc3 | 0.812458 |
| | | | | | | | Sf3a3 | 0.753309 |
| | | | | | | | Nt5c | 0.81377 |
| | | | | | | | Itpa | 0.824185 |
| | | | | | | | Rpa3 | 1.221217 |
| | | | | | | | Pole4 | 0.781664 |
| | | | | | | | Polr2d | 0.77637 |
| | | | | | | | Ercc1 | 0.726278 |
| | | | | | | | Rnmt | 0.819777 |
| | | | | | | | Ak1 | 0.651226 |
| | | | | | | | Sdcbp | 0.69263 |
| | | | | | | | Tyms | 0.659041 |
| | | | | | | | Stx3 | 0.669689 |
| | | | | | | | Nudt21 | 0.669206 |
| | | | | | | | Bcam | 0.685739 |
| | | | | | | | Pola1 | 1.074382 |
| | | | | | | | Usp11 | 1.068471 |
| | | | | | | | Pde4b | 1.03453 |
| | | | | | | | Trp53 | 1.970581 |
| | | | | | | | Gmpr2 | 0.689213 |
| | | | | | | | Aprt | 1.020484 |
| | | | | | | | Hprt | 0.950862 |
| | | | | | | | Pola2 | 0.627075 |
| | | | | | | | Pnp | 1.49676 |

| | | | | | | | | |
|---|---|---|---|---|---|---|---|---|
| | | | | | | | Alyref | 0.853099 |
| | | | | | | | Cda | 0.818999 |
| | | | | | | DOWN | Umps | -0.87647 |
| | | | | | | | Gtf3c5 | -1.56524 |
| | | | | | | | Ccno | -0.63484 |
| | | | | | | | Vps37d | -0.60743 |
| | | | | | | | Edf1 | -0.85087 |
| | | | | | | | Cmpk2 | -1.17684 |
| | | | | | | | Polr2e | -0.58292 |
| | | | | | | | Ago4 | -0.70223 |
| | | | | | | | Nme3 | -0.78205 |
| | | | | | | | Polr1d | -1.14313 |
| | | | | | | | Rev3l | -0.70986 |
| | | | | | | | Tmed2 | -0.67777 |
| | | | | | | | Polr3gl | -0.82243 |
| | | | | | | | Cetn2 | -0.6929 |
| | | | | | | | Guk1 | -1.04229 |
| | | | | | | | Srsf6 | -0.75994 |
| | | | | | | | Poll | -1.21294 |
| | | | | | | | Polr2k | -1.12907 |

**Table S11. Significantly deregulated DNA Repair genes in PDAC from 4-, 6-, and 9-week-old mice compared to healthy pancreases from healthy age-matched littermates.**

| Term | Adjusted P-value | Combined Score | Genes |
| --- | --- | --- | --- |
| Single-Strand Nucleotide Excision | 9.81E-06 | 1842.167347 | DDB1;PCNA;LIG1;ERCC1 |
| Single Strand Nucleotide Excision DNA Repair Suppression in Cancer | 2.00E-05 | 1118.810683 | DDB1;PCNA;LIG1;ERCC1 |
| CUL4/CRL (SCF4) Complex | 7.49E-04 | 622.6319384 | DDB1;PCNA;LIG1 |
| Uric Acid Synthesis in Gout | 0.006634275 | 685.7447231 | PNP;APRT |
| Single-Strand Mismatch | 0.006634275 | 603.0173367 | PCNA;LIG1 |
| Single Strand Mismatch DNA Repair Suppression in Cancer | 0.006634275 | 481.7596464 | PCNA;LIG1 |
| Single-Strand Base Excision | 0.006634275 | 436.1759087 | PCNA;LIG1 |
| DNA Replication in DNA Machinery | 0.006634275 | 436.1759087 | POLA1;PCNA |
| Single Strand Base Excision DNA Repair Impairment in Cancer | 0.007612316 | 364.6775042 | PCNA;LIG1 |
| Androgens Promote Scalp Dermal Papilla Regression | 0.021435875 | 163.4137007 | PCNA;LIG1 |
| Proteins Involved in Female Infertility | 0.071316629 | 60.57955112 | NPR2;ZFP36L2 |
| Proteins Involved in Diffuse Large-B-Cell Lymphoma | 0.071316629 | 59.13136932 | PDE4B;SMAD5 |
| Vitamin D and Folate in Multiple Sclerosis | 0.086277053 | 318.1473008 | TYMS |
| TGFBR -> SMAD1/5/9 Signaling | 0.086277053 | 255.3400953 | SMAD5 |
| Proteins with Altered Expression in Androgenetic Alopecia | 0.086277053 | 255.3400953 | PCNA |
| Folate Cycle and Homocysteine Overproduction | 0.086277053 | 255.3400953 | TYMS |
| HPRT1 Deficiency in Lesch-Nyhan Syndrome | 0.087532899 | 211.6081512 | GUK1 |
| NPR1/NPR2 -> Fatty Acid Signaling | 0.087532899 | 211.6081512 | NPR2 |
| Ubiquitin Ligase Complex in Prostata Cell | 0.093170735 | 179.5646306 | DDB1 |
| RNA Gene Silencing | 0.098219473 | 155.1716014 | DGCR8 |
| Proteins Involved in Oligodendroglioma | 0.102763332 | 136.0428937 | PCNA |
| MTHFR Mutation, Hyperhomocysteinemia and Folate Deficiency | 0.105991715 | 108.1035636 | TYMS |
| Hyperhomocysteinemia Induced Thrombophilia | 0.105991715 | 108.1035636 | TYMS |

| | | | |
|---|---|---|---|
| Dopamine Mediated Glutamate Release/Uptake Circle in Neuron in Migraine | 0.105991715 | 108.1035636 | STX3 |
| Proteins Involved in Urolithiasis | 0.109437376 | 97.63576151 | APRT |
| Vitamins Insufficiency Causes Homocysteine High Level Synthesis | 0.11259876 | 88.80359996 | TYMS |
| Ethanol Induced Hepatotoxicity | 0.113917731 | 74.75896142 | SMAD5 |
| Sister Chromatid Cohesion | 0.113917731 | 69.09833218 | PCNA |
| Epigenetic Alterations Triggers Genomic Instability | 0.113917731 | 69.09833218 | TYMS |
| Renin-Angiotensin-Aldosterone System in Myocardial ischemia | 0.113917731 | 64.13258687 | NPR2 |
| Dopamine Mediated Glutamate Release and Glutamate Uptake Circle | 0.113917731 | 59.74559829 | STX3 |
| Neurogenic Transcription Factors Role in Hirschsprung Disease | 0.113917731 | 59.74559829 | SMAD5 |
| Vascular Smooth Muscle Cell/Pericyte Differentiation and Proliferation | 0.113917731 | 59.74559829 | SMAD5 |
| BMP Signaling Impairment in Granulosa Cell in POF | 0.113917731 | 55.84535629 | SMAD5 |
| EGFR Nuclear Signaling in Colorectal Cancer | 0.113917731 | 52.35806749 | PCNA |
| DNMT and MBD Families Activation in DNA Methylation in Cancer | 0.113917731 | 52.35806749 | PCNA |
| High Level of Homocystine Effects (Methylation Cycle) | 0.113917731 | 49.2238977 | TYMS |
| Bone Resorption in Hyperparathyroidism | 0.113917731 | 49.2238977 | SMAD5 |
| Werner Syndrome (Adult Progeria) | 0.113917731 | 49.2238977 | PCNA |
| TGFB2 Signaling Impairement in Osteoarthritis | 0.115749574 | 46.39384902 | SMAD5 |
| Hereditary Hemorrhagic Telangiectasia | 0.117480175 | 43.82743487 | SMAD5 |
| Osteoblast Function Decline in Gout | 0.119419745 | 39.35600928 | SMAD5 |
| ActivinR/BMPR -> SMAD1/5/9 Signaling | 0.119419745 | 39.35600928 | SMAD5 |
| Histone Methylation | 0.119419745 | 37.39875501 | PCNA |
| BMP/TGF-beta Signaling Impairment in Pulmonary Hypertension | 0.119419745 | 37.39875501 | SMAD5 |
| Enamel Formation Disruption | 0.122190265 | 33.93874529 | SMAD5 |
| BMPR2 Signaling | 0.122190265 | 33.93874529 | SMAD5 |
| Proteins Involved in Otitis Media | 0.122201539 | 32.40358367 | PDE4B |

| | | | |
|---|---|---|---|
| Replication Stress Triggers Genomic Instability | 0.122201539 | 30.98036032 | PCNA |
| Proteins with Altered Expression in Cancer-Associated Dysregulated DNA Repair | 0.122201539 | 30.98036032 | LIG1 |
| Dentin Formation Disruption | 0.123390735 | 29.65781217 | SMAD5 |
| Genes with Mutation in Cancer-Associated Dysregulated DNA Repair | 0.128023099 | 27.27663103 | LIG1 |
| Hyperparathyroidism, Secondary Effect | 0.129030693 | 26.20179786 | SMAD5 |
| Kinetochore Assembly | 0.139992882 | 22.52576227 | ZWINT |
| Bone Loss in Osteoporosis | 0.163258506 | 17.2828352 | SMAD5 |
| Eosinophil Survival in Asthma | 0.163675651 | 16.27688724 | SDCBP |
| Proteins Involved in Endometrial Cancer | 0.163675651 | 16.27688724 | TYMS |
| Bone Remodeling in Hyperthyroidism | 0.187644865 | 12.72902082 | SMAD5 |
| Cetuximab Resistance in Colorectal Cancer | 0.194628863 | 11.26062558 | PCNA |
| Proteins Involved in Helicobacter Infections | 0.194628863 | 10.99906668 | PCNA |
| Androgens in Sebocyte Maturation | 0.194628863 | 10.99906668 | SMAD5 |
| Neutrophil Recruitment and Priming | 0.194628863 | 10.7469535 | PCNA |
| Eosinophil Survival by Cytokine Signaling | 0.204131417 | 9.408240654 | SDCBP |
| Glioma Stem Cell Program Activation | 0.204131417 | 9.408240654 | SMAD5 |
| TGFB Family in Epithelial to Mesenchymal Transition in Cancer | 0.2210802 | 7.991359336 | SMAD5 |
| Osteoarthritis | 0.222612422 | 7.688672189 | SMAD5 |
| Proteins Involved in Polycystic Kidney Disease | 0.254567234 | 5.889232558 | STX3 |
| Proteins Involved in Colorectal Neoplasms | 0.255355897 | 5.698709605 | TYMS |
| Medulloblastoma | 0.262719892 | 5.261284894 | SMAD5 |
| Proteins Involved in Dilated Cardiomyopathy | 0.384815778 | 2.346506007 | DGCR8 |
| Proteins Involved in Chronic Obstructive Pulmonary Disease | 0.384815778 | 2.272621823 | PDE4B |
| Proteins Involved in Male Infertility | 0.419024495 | 1.801867903 | POLL |
| Proteins Involved in Hearing Loss | 0.439064941 | 1.551483611 | SMAD5 |

| | | | |
|---|---|---|---|
| Proteins Involved in Melanoma | 0.476584664 | 1.206738375 | SDCBP |
| Proteins Involved in Arterial Hypertension | 0.49388174 | 1.051832804 | PDE4B |

**Table S12. Gene Set Enrichment Analysis on significantly deregulated DNA Repair genes in 9-week-old PDAC pancreases compared to pancreases from age-matched healthy littermates.**

| Name/Target | Forward (5->3) | Reverse (5->3) | RefSeq mRNA | Amplicon size |
|---|---|---|---|---|
| *Tbp* | GAAGAACAATCCAGACTAGCAGCA | CCTTATAGGGAACTTCACATCACAG | NM_013684.3 | 129 |
| *Impdh1* | CATGGAGGAACCGCTCTCAC | GGAGGATCAGGAAGTCGTTGT | >NM_001302933.1 | 290 |
| *Impdh2* | TGTGGATGTAGTGGTTTTGGACT | TTGGGGCTGTGGGACTTTATG | >NM_001378921.1 | 249 |
| *Adss* | TGGATTTACTGCGTTGGCCC | TTTGTGCGTTGACAGGTAGC | >NM_007422.3 | 221 |
| *Adsl* | GTCACCTGATGGCCCTTACC | TGCGCCGTTCAATTACTTTGG | >NM_009634.6 | 192 |
| *Ercc1* | AAAGATCCCCAGCAGGCTCTC | AGCTGTTCCAGGGATCCAAATG | NM_007948.3 | 263 |
| *Gtf2h1* | GCCAGCAGTCAAAAGGGCAA | CAGGTGACAGGGCTGTGATG | NM_001360075.1 | 272 |
| *Rad23a* | GGAACCTGACGAGACGGTAA | TGCCAGCATAGATGAGTTTCTG | NM_001378894.1 | 101 |
| *Pold4* | TGGGCCTTGTACAGGTATCACA | TCAGGGTGTGCCTTCAACAC | NM_027196.4 | 105 |
| *Polb* | CCAGGCGATCCACAAGTACA | TGTTCCTACTCCTGGCAGTTTC | NM_011130.2 | 112 |
| *Parp9* | GCATTTGCTAAAGAGCACAAGGA | AAGCACCACTATTACCGCTGA | NM_001405299.1 | 142 |

**Table S13. Primer sequences for qRT-PCR.**

| Antibodies and probes | | | |
|---|---|---|---|
| **Target** | **Name** | **Reference Number** | **Manufacturer** |
| Integrin β1 | BD OptiBuild™ BV421 Mouse Anti-Human CD29 | 743783 | BD Biosciences |
| PDGFRB | BD OptiBuild™ BV786 Mouse Anti-Human CD140b | 743038 | BD Biosciences |
| FAP | Human Fibroblast Activation Protein alpha /FAP Alexa Fluor® 647-conjugated Antibody | FAB3715R | R&D Systems |
| Annexin VI | Annexin VI Antibody (N-19) | sc-1931 | Santa Cruz Biotechnology |
| αSMA | Human alpha-Smooth Muscle Actin APC-conjugated Antibody | IC1420A | R&D Systems |
| GAPDH | Anti-GAPDH antibody - Loading Control | ab9485 | Abcam |
| Fibronectin | BD Transduction Laboratories™ Purified Mouse Anti-Fibronectin | 610078 | BD Biosciences |
| YAP1 | Anti-active YAP1 antibody [EPR19812] | ab205270 | Abcam |
| Gda | Guanine Deaminase Antibody | GTX33233 | GeneTex |
| β-tubulin | Anti-b-Tubulin, clone TUB 2.1 | T4026 | Sigma-Aldrich |
| γH2AX | Recombinant Anti-gamma H2A.X (phospho S139) antibody [EP854(2)Y] | ab81299 | Abcam |
| S6K | p70 S6 Kinase (49D7) Rabbit mAb | #2708 | Cell Signaling Technology |
| p-S6K | Phospho-p70 S6 Kinase (Thr389) (108D2) Rabbit mAb | #9234 | Cell Signaling Technology |
| Akt | Akt Antibody | #9272 | Cell Signaling Technology |
| p-Akt | Phospho-Akt (Ser473) Antibody | #9271 | Cell Signaling Technology |
| Mouse IgG | Goat-anti-Mouse IgG-HRP | 1030-05 | Southern Biotech |
| Rabbit IgG | Goat Anti-Rabbit IgG-HRP | 4030-05 | Southern Biotech |
| Rabbit IgG | Donkey anti-Rabbit IgG (H+L) Highly Cross-Adsorbed Secondary Antibody, Alexa Fluor™ Plus 488 | A32790 | Invitrogen |
| Mouse IgG | Donkey anti-Mouse IgG (H+L) Highly Cross-Adsorbed Secondary Antibody, Alexa Fluor™ 568 | A10037 | Invitrogen |
| Actin | Alexa Fluor™ 647 Phalloidin | A22287 | Invitrogen |

**Table S14. Antibodies and Probes.**

| Reference | Source of data | Technological platform | N° of probe sets/genes | All samples | Primary PDAC samples included in the present analysis |
|---|---|---|---|---|---|
| Badea et al., Hepatogastroenterology 2008 | GEO database, GSE15471 | Affymetrix, array U133 Plus 2.0 | 54K | 78 | 36 |
| van den Broeck et al., J Exp Clin Cancer Res 2012 | GEO database, GSE42952 | Affymetrix, array U133 Plus 2.0 | 54K | 23 | 12 |
| Zhang et al., PLoS One 2012 & Clin Cencer Res 2013 | GEO database, GSE28735 | Affymetrix, array Gene 1.0 ST | 33K | 90 | 45 |
| Lunardi et al., Oncotarget 2014 | GEO database, GSE55643 | Agilent, array 4x44K G4112F (014850) | 44K | 53 | 45 |
| Park et al., Mod Pathol 2014 | GEO database, GSE43795 | Illumina, array Human HT-12 V4.0 | 48K | 31 | 6 |
| Winter et al., PLoS Comput Biol 2012 | Array-Express database, E-MEXP-2780 | Affymetrix, array U133 Plus 2.0 | 54K | 30 | 30 |
| Grutzmann et al., Neoplasia 2004 | Array-Express database, E-MEXP-950 | Affymetrix, array U133 A+B | 22K+22K | 25 | 11 |
| TCGA, PAAD | TCGA portal, https://tcga-data.nci.nih.gov | Illumina, RNA sequencing V2 | 25K | 183 | 150 |
| Straford et al., PLoS Med 2010 | GEO database, GSE21501 | Agilent, array 4x44K G4112F (014850) | 44K | 132 | 132 |
| Monzon et al., Clin Oncol 2009 | GEO database, GSE12630 | Affymetrix, array U133 A | 22K | 24 | 24 |
| Bailey et al., Nature 2016 | European Genome-phenome Archive (EGA), EGAS00001000154 | Illumina, RNA sequencing HiSeq | 18K | 96 | 96 |
| Chen et al., PLoS ONE 2015 | GEO database, GSE57495 | Affymetrix, Rosetta/Merck RSTA Custom 2.0 | 60K | 63 | 63 |

| | | | | | |
|---|---|---|---|---|---|
| Collisson et al., Nat Med. 2011 | GEO database, GSE17891 | Affymetrix, array U133 Plus 2.0 | 54K | 27 | 27 |
| Chaika et al., PLOSOne 2012 | GEO database, GSE34153 | Agilent, array 4x44K G4112F (014850) | 44K | 75 | 15 |
| ICGC, PACA CA (2019) | https://dcc.icgc.org/projects/PACA-CA | Illumina, RNA sequencing HiSeq | 19K | 264 | 195 |
| Moffitt et al., Nat Genet 2015 | GEO database, GSE71729 | Agilent, array 4x44K G4112F (014850) | 44K | 357 | 0 |
| Kirby et al., Mol Oncol. 2016 | GEO database, GSE79670 | Illumina, RNA sequencing HiSeq | 49K | 51 | 51 |
| **TOTAL** | | | | **1602** | **938** |

**Table S15. Human PDAC datasets used in the study.**

| Symbol | Description | Cytoband | Entrez Gene ID |
|---|---|---|---|
| *COL11A1* | collagen type XI alpha 1 chain | 1p21.1 | 1301 |
| *COL12A1* | collagen type XII alpha 1 chain | 6q13-q14.1 | 1303 |
| *COL15A1* | collagen type XV alpha 1 chain | 9q22.33 | 1306 |
| *COL16A1* | collagen type XVI alpha 1 chain | 1p35.2 | 1307 |
| *COL1A1* | collagen type I alpha 1 chain | 17q21.33 | 1277 |
| *COL1A2* | collagen type I alpha 2 chain | 7q21.3 | 1278 |
| *COL27A1* | collagen type XXVII alpha 1 chain | 9q32 | 85301 |
| *COL3A1* | collagen type III alpha 1 chain | 2q32.2 | 1281 |
| *COL4A5* | collagen type IV alpha 5 chain | Xq22.3 | 1287 |
| *COL4A6* | collagen type IV alpha 6 chain | Xq22.3 | 1288 |
| *COL5A1* | collagen type V alpha 1 chain | 9q34.3 | 1289 |
| *COL6A1* | collagen type VI alpha 1 chain | 21q22.3 | 1291 |
| *COL6A2* | collagen type VI alpha 2 chain | 21q22.3 | 1292 |
| *COL6A3* | collagen type VI alpha 3 chain | 2q37.3 | 1293 |
| *COL8A1* | collagen type VIII alpha 1 chain | 3q12.1 | 1295 |
| *AEBP1* | AE binding protein 1 | 7p13 | 165 |
| *BMPER* | BMP binding endothelial regulator | 7p14.3 | 168667 |
| *CRIM1* | cysteine rich transmembrane BMP regulator 1 | 2p22.2 | 51232 |
| *CCN2* | cellular communication network factor 2 | 6q23.2 | 1490 |
| *CTHRC1* | collagen triple helix repeat containing 1 | 8q22.3 | 115908 |
| *CCN1* | cellular communication network factor 1 | 1p22.3 | 3491 |
| *DMP1* | dentin matrix acidic phosphoprotein 1 | 4q22.1 | 1758 |
| *EDIL3* | EGF like repeats and discoidin domains 3 | 5q14.3 | 10085 |
| *EFEMP1* | EGF containing fibulin extracellular matrix protein 1 | 2p16.1 | 2202 |
| *EFEMP2* | EGF containing fibulin extracellular matrix protein 2 | 11q13.1 | 30008 |
| *EMILIN1* | elastin microfibril interfacer 1 | 2p23.3 | 11117 |
| *FBLN5* | fibulin 5 | 14q32.12 | 10516 |
| *FBN1* | fibrillin 1 | 15q21.1 | 2200 |
| *FN1* | fibronectin 1 | 2q35 | 2335 |
| *GLDN* | gliomedin | 15q21.2 | 342035 |
| *IGFBP6* | insulin like growth factor binding protein 6 | 12q13.13 | 3489 |
| *IGSF10* | immunoglobulin superfamily member 10 | 3q25.1 | 285313 |
| *LAMA5* | laminin subunit alpha 5 | 20q13.33 | 3911 |
| *LAMB2* | laminin subunit beta 2 | 3p21.31 | 3913 |
| *LAMB3* | laminin subunit beta 3 | 1q32.2 | 3914 |
| *LAMC1* | laminin subunit gamma 1 | 1q25.3 | 3915 |
| *LAMC2* | laminin subunit gamma 2 | 1q25.3 | 3918 |
| *LAMC3* | laminin subunit gamma 3 | 9q34.12 | 10319 |
| *LGI2* | leucine rich repeat LGI family member 2 | 4p15.2 | 55203 |

| | | | |
|---|---|---|---|
| *LGI4* | leucine rich repeat LGI family member 4 | 19q13.12\|19q13.11 | 163175 |
| *LTBP1* | latent transforming growth factor beta binding protein 1 | 2p22.3 | 4052 |
| *LTBP2* | latent transforming growth factor beta binding protein 2 | 14q24.3 | 4053 |
| *LTBP3* | latent transforming growth factor beta binding protein 3 | 11q13.1 | 4054 |
| *MATN2* | matrilin 2 | 8q22.1-q22.2 | 4147 |
| *MFGE8* | milk fat globule EGF and factor V/VIII domain containing | 15q26.1 | 4240 |
| *MGP* | matrix Gla protein | 12p12.3 | 4256 |
| *NDNF* | neuron derived neurotrophic factor | 4q27 | 79625 |
| *NID1* | nidogen 1 | 1q42.3 | 4811 |
| *NID2* | nidogen 2 | 14q22.1 | 22795 |
| *NPNT* | nephronectin | 4q24 | 255743 |
| *NTN1* | netrin 1 | 17p13.1 | 9423 |
| *NTN3* | netrin 3 | 16p13.3 | 4917 |
| *PCOLCE* | procollagen C-endopeptidase enhancer | 7q22.1 | 5118 |
| *PCOLCE2* | procollagen C-endopeptidase enhancer 2 | 3q23 | 26577 |
| *RSPO2* | R-spondin 2 | 8q23.1 | 340419 |
| *SLIT2* | slit guidance ligand 2 | 4p15.31 | 9353 |
| *SLIT3* | slit guidance ligand 3 | 5q34-q35.1 | 6586 |
| *SMOC1* | SPARC related modular calcium binding 1 | 14q24.2 | 64093 |
| *SNED1* | sushi, nidogen and EGF like domains 1 | 2q37.3 | 25992 |
| *SPON2* | spondin 2 | 4p16.3 | 10417 |
| *SPP1* | secreted phosphoprotein 1 | 4q22.1 | 6696 |
| *THBS2* | thrombospondin 2 | 6q27 | 7058 |
| *THBS3* | thrombospondin 3 | 1q22 | 7059 |
| *TNC* | tenascin C | 9q33.1 | 3371 |
| *VWA1* | von Willebrand factor A domain containing 1 | 1p36.33 | 64856 |
| *VWA5A* | von Willebrand factor A domain containing 5A | 11q24.2 | 4013 |
| *CCN4* | cellular communication network factor 4 | 8q24.22 | 8840 |
| *CCN5* | cellular communication network factor 5 | 20q13.12 | 8839 |
| *ACAN* | aggrecan | 15q26.1 | 176 |
| *BCAN* | brevican | 1q23.1 | 63827 |
| *BGN* | biglycan | Xq28 | 633 |
| *ESM1* | endothelial cell specific molecule 1 | 5q11.2 | 11082 |
| *HSPG2* | heparan sulfate proteoglycan 2 | 1p36.12 | 3339 |
| *OGN* | osteoglycin | 9q22.31 | 4969 |
| *OMD* | osteomodulin | 9q22.31 | 4958 |
| *PODN* | podocan | 1p32.3 | 127435 |
| *PODNL1* | podocan like 1 | 19p13.12 | 79883 |
| *SPOCK2* | SPARC (osteonectin), cwcv and kazal like domains proteoglycan 2 | 10q22.1 | 9806 |
| *ANXA1* | annexin A1 | 9q21.13 | 301 |
| *ANXA11* | annexin A11 | 10q22.3 | 311 |
| *ANXA2* | annexin A2 | 15q22.2 | 302 |

| | | | |
|---|---|---|---|
| *ANXA7* | annexin A7 | 10q22.2 | 310 |
| *C1QTNF 1* | C1q and TNF related 1 | 17q25.3 | 114897 |
| *CLEC2D* | C-type lectin domain family 2 member D | 12p13.31 | 29121 |
| *CLEC4D* | C-type lectin domain family 4 member D | 12p13.31 | 338339 |
| *COLEC1 2* | collectin subfamily member 12 | 18p11.32 | 81035 |
| *GPC6* | glypican 6 | 13q31.3-q32.1 | 10082 |
| *LGALS1* | galectin 1 | 22q13.1 | 3956 |
| *LGALS3* | galectin 3 | 14q22.3 | 3958 |
| *LGALS9* | galectin 9 | 17q11.2 | 3965 |
| *LGALSL* | galectin like | 2p14 | 29094 |
| *LMAN1* | lectin, mannose binding 1 | 18q21.32 | 3998 |
| *MUC1* | mucin 1, cell surface associated | 1q22 | 4582 |
| *PLXNA4* | plexin A4 | 7q32.3 | 91584 |
| *SDC4* | syndecan 4 | 20q13.12 | 6385 |
| *SEMA3C* | semaphorin 3C | 7q21.11 | 10512 |
| *SEMA3D* | semaphorin 3D | 7q21.11 | 223117 |
| *SEMA3E* | semaphorin 3E | 7q21.11 | 9723 |
| *SEMA3F* | semaphorin 3F | 3p21.31 | 6405 |
| *SEMA4B* | semaphorin 4B | 15q26.1 | 10509 |
| *SEMA4D* | semaphorin 4D | 9q22.2 | 10507 |
| *SEMA4F* | ssemaphorin 4F | 2p13.1 | 10505 |
| *SEMA4G* | semaphorin 4G | 10q24.31 | 57715 |
| *SEMA5A* | semaphorin 5A | 5p15.31 | 9037 |
| *SEMA6A* | semaphorin 6A | 5q23.1 | 57556 |
| *SEMA6C* | semaphorin 6C | 1q21.3 | 10500 |
| *ADAM12* | ADAM metallopeptidase domain 12 | 10q26.2 | 8038 |
| *ADAM19* | ADAM metallopeptidase domain 19 | 5q33.3 | 8728 |
| *ADAM22* | ADAM metallopeptidase domain 22 | 7q21.12 | 53616 |
| *ADAM9* | ADAM metallopeptidase domain 9 | 8p11.22 | 8754 |
| *ADAMTS 14* | ADAM metallopeptidase with thrombospondin type 1 motif 14 | 10q22.1 | 140766 |
| *ADAMTS 2* | ADAM metallopeptidase with thrombospondin type 1 motif 2 | 5q35.3 | 9509 |
| *ADAMTS 6* | ADAM metallopeptidase with thrombospondin type 1 motif 6 | 5q12.3 | 11174 |
| *ADAMTS 7* | ADAM metallopeptidase with thrombospondin type 1 motif 7 | 15q25.1 | 11173 |
| *ADAMTS 9* | ADAM metallopeptidase with thrombospondin type 1 motif 9 | 3p14.1 | 56999 |
| *ADAMTS L4* | ADAMTS like 4 | 1q21.2 | 54507 |
| *ADAMTS L5* | ADAMTS like 5 | 19p13.3 | 339366 |
| *CD109* | CD109 molecule | 6q13 | 135228 |
| *CELA1* | chymotrypsin like elastase 1 | 12q13.13 | 1990 |
| *CST6* | cystatin E/M | 11q13.1 | 1474 |
| *CSTB* | cystatin B | 21q22.3 | 1476 |

| | | | |
|---|---|---|---|
| *CTSB* | cathepsin B | 8p23.1 | 1508 |
| *CTSC* | cathepsin C | 11q14.2 | 1075 |
| *CTSE* | cathepsin E | 1q32.1 | 1510 |
| *CTSF* | cathepsin F | 11q13.2 | 8722 |
| *CTSH* | cathepsin H | 15q25.1 | 1512 |
| *CTSO* | cathepsin O | 4q32.1 | 1519 |
| *CTSZ* | cathepsin Z | 20q13.32 | 1522 |
| *HPSE* | heparanase | 4q21.23 | 10855 |
| *HTRA3* | HtrA serine peptidase 3 | 4p16.1 | 94031 |
| *HYAL1* | hyaluronidase 1 | 3p21.31 | 3373 |
| *HYAL2* | hyaluronidase 2 | 3p21.31 | 8692 |
| *ITIH2* | inter-alpha-trypsin inhibitor heavy chain 2 | 10p14 | 3698 |
| *ITIH5* | inter-alpha-trypsin inhibitor heavy chain 5 | 10p14 | 80760 |
| *KAZALD 1* | Kazal type serine peptidase inhibitor domain 1 | 10q24.31 | 81621 |
| *LOX* | lysyl oxidase | 5q23.1 | 4015 |
| *LOXL1* | lysyl oxidase like 1 | 15q24.1 | 4016 |
| *LOXL2* | lysyl oxidase like 2 | 8p21.3 | 4017 |
| *LOXL3* | lysyl oxidase like 3 | 2p13.1 | 84695 |
| *MASP1* | MBL associated serine protease 1 | 3q27.3 | 5648 |
| *MMP16* | matrix metallopeptidase 16 | 8q21.3 | 4325 |
| *MMP19* | matrix metallopeptidase 19 | 12q13.2 | 4327 |
| *MMP2* | matrix metallopeptidase 2 | 16q12.2 | 4313 |
| *MMP3* | matrix metallopeptidase 3 | 11q22.2 | 4314 |
| *NGLY1* | N-glycanase 1 | 3p24.2 | 55768 |
| *OGFOD2* | 2-oxoglutarate and iron dependent oxygenase domain containing 2 | 12q24.31 | 79676 |
| *P4HA3* | prolyl 4-hydroxylase subunit alpha 3 | 11q13.4 | 283208 |
| *P4HTM* | prolyl 4-hydroxylase, transmembrane | 3p21.31\|3p21.3 | 54681 |
| *PAPPA* | pappalysin 1 | 9q33.1 | 5069 |
| *PCSK6* | proprotein convertase subtilisin/kexin type 6 | 15q26.3 | 5046 |
| *SERPIN B2* | serpin family B member 2 | 18q21.33-q22.1 | 5055 |
| *SERPIN B8* | serpin family B member 8 | 18q22.1 | 5271 |
| *SERPINF 1* | serpin family F member 1 | 17p13.3 | 5176 |
| *SERPIN G1* | serpin family G member 1 | 11q12.1 | 710 |
| *SLPI* | secretory leukocyte peptidase inhibitor | 20q13.12 | 6590 |
| *SULF1* | sulfatase 1 | 8q13.2-q13.3 | 23213 |
| *SULF2* | sulfatase 2 | 20q13.12 | 55959 |
| *TIMP1* | TIMP metallopeptidase inhibitor 1 | Xp11.3 | 7076 |
| *TIMP3* | TIMP metallopeptidase inhibitor 3 | 22q12.3 | 7078 |
| *ANGPT1* | angiopoietin 1 | 8q23.1 | 284 |
| *ANGPT2* | angiopoietin 2 | 8p23.1 | 285 |
| *ARTN* | artemin | 1p34.1 | 9048 |
| *BDNF* | brain derived neurotrophic factor | 11p14.1 | 627 |
| *BMP4* | bone morphogenetic protein 4 | 14q22.2 | 652 |

| | | | |
|---|---|---|---|
| *CCBE1* | collagen and calcium binding EGF domains 1 | 18q21.32 | 147372 |
| *CCL2* | C-C motif chemokine ligand 2 | 17q12 | 6347 |
| *CCL20* | C-C motif chemokine ligand 20 | 2q36.3 | 6364 |
| *CCL7* | C-C motif chemokine ligand 7 | 17q12 | 6354 |
| *CHRD* | chordin | 3q27.1 | 8646 |
| *CRLF1* | cytokine receptor like factor 1 | 19p13.11 | 9244 |
| *CSF1* | colony stimulating factor 1 | 1p13.3 | 1435 |
| *CTF1* | cardiotrophin 1 | 16p11.2 | 1489 |
| *CX3CL1* | C-X3-C motif chemokine ligand 1 | 16q21 | 6376 |
| *CXCL10* | C-X-C motif chemokine ligand 10 | 4q21.1 | 3627 |
| *CXCL12* | C-X-C motif chemokine ligand 12 | 10q11.21 | 6387 |
| *EGFL7* | EGF like domain multiple 7 | 9q34.3 | 51162 |
| *C1QTNF 12* | C1q and TNF related 12 | 1p36.33 | 388581 |
| *ERFE* | erythroferrone | 2q37.3 | 151176 |
| *FGF18* | fibroblast growth factor 18 | 5q35.1 | 8817 |
| *FGF7* | fibroblast growth factor 7 | 15q21.2 | 2252 |
| *FGFBP3* | fibroblast growth factor binding protein 3 | 10q23.32 | 143282 |
| *GDF11* | growth differentiation factor 11 | 12q13.2 | 10220 |
| *IL16* | interleukin 16 | 15q25.1 | 3603 |
| *IL18* | interleukin 18 | 11q23.1 | 3606 |
| *IL34* | interleukin 34 | 16q22.1 | 146433 |
| *INHBA* | inhibin subunit beta A | 7p14.1 | 3624 |
| *INHBB* | inhibin subunit beta B | 2q14.2 | 3625 |
| *ISM1* | isthmin 1 | 20p12.1 | 140862 |
| *MEGF10* | multiple EGF like domains 10 | 5q23.2 | 84466 |
| *MEGF6* | multiple EGF like domains 6 | 1p36.32 | 1953 |
| *MST1* | macrophage stimulating 1 | 3p21.31 | 4485 |
| *NGF* | nerve growth factor | 1p13.2 | 4803 |
| *NRG1* | neuregulin 1 | 8p12 | 3084 |
| *NRTN* | neurturin | 19p13.3 | 4902 |
| *PDGFB* | platelet derived growth factor subunit B | 22q13.1 | 5155 |
| *PDGFC* | platelet derived growth factor C | 4q32.1 | 56034 |
| *PDGFD* | platelet derived growth factor D | 11q22.3 | 80310 |
| *PIK3IP1* | phosphoinositide-3-kinase interacting protein 1 | 22q12.2 | 113791 |
| *PTN* | pleiotrophin | 7q33 | 5764 |
| *S100A10* | S100 calcium binding protein A10 | 1q21.3 | 6281 |
| *S100A11* | S100 calcium binding protein A11 | 1q21.3 | 6282 |
| *S100A3* | S100 calcium binding protein A3 | 1q21.3 | 6274 |
| *S100A7A* | S100 calcium binding protein A7A | 1q21.3 | 338324 |
| *S100B* | S100 calcium binding protein B | 21q22.3 | 6285 |
| *SCUBE3* | signal peptide, CUB domain and EGF like domain containing 3 | 6p21.31 | 222663 |
| *SFRP1* | secreted frizzled related protein 1 | 8p11.21 | 6422 |
| *TGFB2* | transforming growth factor beta 2 | 1q41 | 7042 |
| *THPO* | thrombopoietin | 3q27.1 | 7066 |
| *TNFSF12* | TNF superfamily member 12 | 17p13.1 | 8742 |
| *VEGFC* | vascular endothelial growth factor C | 4q34.3 | 7424 |

| | | | |
|---|---|---|---|
| *WNT10B* | Wnt family member 10B | 12q13.12 | 7480 |
| *WNT5B* | Wnt family member 5B | 12p13.33 | 81029 |
| *WNT7A* | Wnt family member 7A | 3p25.1 | 7476 |
| *WNT9A* | Wnt family member 9A | 1q42.13 | 7483 |
| *FCN1* | ficolin 1 | 9q34.3 | 2219 |
| *FCN2* | ficolin 2 | 9q34.3 | 2220 |
| *MMP23B* | matrix metallopeptidase 23B | 1p36.33 | 8510 |
| *SERPIN B1* | serpin family B member 1 | 6p25.2 | 1992 |
| *HMSD* | histocompatibility minor serpin domain containing | 18q22.1 | 284293 |
| *SERPIN B9* | serpin family B member 9 | 6p25.2 | 5272 |
| *PRL* | prolactin | 6p22.3 | 5617 |

**Table S16. List of human matrisomal genes extracted from public human PDAC data sets.**

| Symbol | Description | Cytoband | Entrez Gene ID |
|---|---|---|---|
| ADA | adenosine deaminase | 20q13.12 | 100 |
| ADCY6 | adenylate cyclase 6 | 12q13.12 | 112 |
| ADRM1 | ADRM1 26S proteasome ubiquitin receptor | 20q13.33 | 11047 |
| AGO4 | argonaute RISC component 4 | 1p34.3 | 192670 |
| AK1 | adenylate kinase 1 | 9q34.11 | 203 |
| AK4P3 | adenylate kinase 4 pseudogene 3 | 12p11.21 | 645619 |
| ALYREF | Aly/REF export factor | 17q25.3 | 10189 |
| APRT | adenine phosphoribosyltransferase | 16q24.3 | 353 |
| ARL6IP1 | ADP ribosylation factor like GTPase 6 interacting protein 1 | 16p12.3 | 23204 |
| BCAM | basal cell adhesion molecule (Lutheran blood group) | 19q13.32 | 4059 |
| BCAP31 | B cell receptor associated protein 31 | Xq28 | 10134 |
| BOLA2B | bolA family member 2B | 16p11.2 | 654483 |
| CANT1 | calcium activated nucleotidase 1 | 17q25.3 | 124583 |
| CBX4 | chromobox 4 | 17q25.3 | 8535 |
| CCNO | cyclin O | 5q11.2 | 10309 |
| CDA | cytidine deaminase | 1p36.12 | 978 |
| CETN2 | centrin 2 | Xq28 | 1069 |
| CMPK2 | cytidine/uridine monophosphate kinase 2 | 2p25.2 | 129607 |
| COX17 | cytochrome c oxidase copper chaperone COX17 | 3q13.33 | 10063 |
| CSTF3 | cleavage stimulation factor subunit 3 | 11p13 | 1479 |
| DAD1 | defender against cell death 1 | 14q11.2 | 1603 |
| DCTN4 | dynactin subunit 4 | 5q33.1 | 51164 |
| DDB1 | damage specific DNA binding protein 1 | 11q12.2 | 1642 |
| DDB2 | damage specific DNA binding protein 2 | 11p11.2 | 1643 |
| DGCR8 | DGCR8 microprocessor complex subunit | 22q11.21 | 54487 |
| DGUOK | deoxyguanosine kinase | 2p13.1 | 1716 |
| DUT | deoxyuridine triphosphatase | 15q21.1 | 1854 |
| EDF1 | endothelial differentiation related factor 1 | 9q34.3 | 8721 |
| EIF1B | eukaryotic translation initiation factor 1B | 3p22.1 | 10289 |
| ELL | elongation factor for RNA polymerase II | 19p13.11 | 8178 |
| ELOA | elongin A | 1p36.11 | 6924 |
| ERCC1 | ERCC excision repair 1, endonuclease non-catalytic subunit | 19q13.32 | 2067 |
| ERCC2 | ERCC excision repair 2, TFIIH core complex helicase subunit | 19q13.32 | 2068 |
| ERCC3 | ERCC excision repair 3, TFIIH core complex helicase subunit | 2q14.3 | 2071 |
| ERCC4 | ERCC excision repair 4, endonuclease catalytic subunit | 16p13.12 | 2072 |
| ERCC5 | ERCC excision repair 5, endonuclease | 13q33.1 | 2073 |
| ERCC8 | ERCC excision repair 8, CSA ubiquitin ligase complex subunit | 5q12.1 | 1161 |
| FEN1 | flap structure-specific endonuclease 1 | 11q12.2 | 2237 |
| GMPR2 | guanosine monophosphate reductase 2 | 14q12 | 51292 |
| GPX4 | glutathione peroxidase 4 | 19p13.3 | 2879 |
| GSDME | gasdermin E | 7p15.3 | 1687 |

| | | | |
|---|---|---|---|
| GTF2A2 | general transcription factor IIA subunit 2 | 15q22.2 | 2958 |
| GTF2B | general transcription factor IIB | 1p22.2 | 2959 |
| GTF2F1 | general transcription factor IIF subunit 1 | 19p13.3 | 2962 |
| GTF2H1 | general transcription factor IIH subunit 1 | 11p15.1 | 2965 |
| GTF2H3 | general transcription factor IIH subunit 3 | 12q24.31 | 2967 |
| GTF2H5 | general transcription factor IIH subunit 5 | 6q25.3 | 404672 |
| GTF3C5 | general transcription factor IIIC subunit 5 | 9q34.13 | 9328 |
| GUK1 | guanylate kinase 1 | 1q42.13 | 2987 |
| HCLS1 | hematopoietic cell-specific Lyn substrate 1 | 3q13.33 | 3059 |
| HEXIM1 | HEXIM P-TEFb complex subunit 1 | 17q21.31 | 10614 |
| HPRT1 | hypoxanthine phosphoribosyltransferase 1 | Xq26.2-q26.3 | 3251 |
| IMPDH2 | inosine monophosphate dehydrogenase 2 | 3p21.31 | 3615 |
| ITPA | inosine triphosphatase | 20p13 | 3704 |
| LIG1 | DNA ligase 1 | 19q13.33 | 3978 |
| MPG | N-methylpurine DNA glycosylase | 16p13.3 | 4350 |
| MRPL40 | mitochondrial ribosomal protein L40 | 22q11.21 | 64976 |
| NCBP2 | nuclear cap binding protein subunit 2 | 3q29 | 22916 |
| NELFB | negative elongation factor complex member B | 9q34.3 | 25920 |
| NELFCD | negative elongation factor complex member C/D | 20q13.32 | 51497 |
| NELFE | negative elongation factor complex member E | 6p21.33 | 7936 |
| NFX1 | nuclear transcription factor, X-box binding 1 | 9p13.3 | 4799 |
| NME1 | NME/NM23 nucleoside diphosphate kinase 1 | 17q21.33 | 4830 |
| NME3 | NME/NM23 nucleoside diphosphate kinase 3 | 16p13.3 | 4832 |
| NME4 | NME/NM23 nucleoside diphosphate kinase 4 | 16p13.3 | 4833 |
| NPR2 | natriuretic peptide receptor 2 | 9p13.3 | 4882 |
| NT5C | 5', 3'-nucleotidase, cytosolic | 17q25.1 | 30833 |
| NT5C3A | 5'-nucleotidase, cytosolic IIIA | 7p14.3 | 51251 |
| NUDT21 | nudix hydrolase 21 | 16q13 | 11051 |
| NUDT9 | nudix hydrolase 9 | 4q22.1 | 53343 |
| PCNA | proliferating cell nuclear antigen | 20p12.3 | 5111 |
| PDE4B | phosphodiesterase 4B | 1p31.3 | 5142 |
| PDE6G | phosphodiesterase 6G | 17q25.3 | 5148 |
| PNP | purine nucleoside phosphorylase | 14q11.2 | 4860 |
| POLA1 | DNA polymerase alpha 1, catalytic subunit | Xp22.11-p21.3 | 5422 |
| POLA2 | DNA polymerase alpha 2, accessory subunit | 11q13.1 | 23649 |
| POLB | DNA polymerase beta | 8p11.21 | 5423 |
| POLD1 | DNA polymerase delta 1, catalytic subunit | 19q13.33 | 5424 |
| POLD3 | DNA polymerase delta 3, accessory subunit | 11q13.4 | 10714 |
| POLD4 | DNA polymerase delta 4, accessory subunit | 11q13.2 | 57804 |
| POLE4 | DNA polymerase epsilon 4, accessory subunit | 2p12 | 56655 |
| POLH | DNA polymerase eta | 6p21.1 | 5429 |
| POLL | DNA polymerase lambda | 10q24.32 | 27343 |
| POLR1B | RNA polymerase I subunit B | 2q14.1 | 84172 |
| POLR1C | RNA polymerase I and III subunit C | 6p21.1 | 9533 |

| | | | |
|---|---|---|---|
| POLR1D | RNA polymerase I and III subunit D | 13q12.2 | 51082 |
| POLR1H | RNA polymerase I subunit H | 6p22.1 | 30834 |
| POLR2A | RNA polymerase II subunit A | 17p13.1 | 5430 |
| POLR2C | RNA polymerase II subunit C | 16q21 | 5432 |
| POLR2D | RNA polymerase II subunit D | 2q14.3 | 5433 |
| POLR2E | RNA polymerase II, I and III subunit E | 19p13.3 | 5434 |
| POLR2F | RNA polymerase II, I and III subunit F | 22q13.1 | 5435 |
| POLR2G | RNA polymerase II subunit G | 11q12.3 | 5436 |
| POLR2H | RNA polymerase II, I and III subunit H | 3q27.1 | 5437 |
| POLR2J3 | RNA polymerase II subunit J3 | 7q22.1 | 548644 |
| POLR2K | RNA polymerase II, I and III subunit K | 8q22.2 | 5440 |
| POLR3C | RNA polymerase III subunit C | 1q21.1 | 10623 |
| POLR3GL | RNA polymerase III subunit GL | 1q21.1 | 84265 |
| POM121C | POM121 transmembrane nucleoporin C | 7q11.23 | 100101267 |
| PRIM1 | DNA primase subunit 1 | 12q13.3 | 5557 |
| RAD52 | RAD52 homolog, DNA repair protein | 12p13.33 | 5893 |
| RAE1 | ribonucleic acid export 1 | 20q13.31 | 8480 |
| RALA | RAS like proto-oncogene A | 7p14.1 | 5898 |
| RBX1 | ring-box 1 | 22q13.2 | 9978 |
| REV3L | REV3 like, DNA directed polymerase zeta catalytic subunit | 6q21 | 5980 |
| RFC2 | replication factor C subunit 2 | 7q11.23 | 5982 |
| RFC3 | replication factor C subunit 3 | 13q13.2 | 5983 |
| RFC4 | replication factor C subunit 4 | 3q27.3 | 5984 |
| RFC5 | replication factor C subunit 5 | 12q24.23 | 5985 |
| RNMT | RNA guanine-7 methyltransferase | 18p11.21 | 8731 |
| RPA3 | replication protein A3 | 7p21.3 | 6119 |
| RRM2B | ribonucleotide reductase regulatory TP53 inducible subunit M2B | 8q22.3 | 50484 |
| SAC3D1 | SAC3 domain containing 1 | 11q13.1 | 29901 |
| SDCBP | syndecan binding protein | 8q12.1 | 6386 |
| SEC61A1 | SEC61 translocon subunit alpha 1 | 3q21.3 | 29927 |
| SF3A3 | splicing factor 3a subunit 3 | 1p34.3 | 10946 |
| SMAD5 | SMAD family member 5 | 5q31.1 | 4090 |
| SNAPC4 | small nuclear RNA activating complex polypeptide 4 | 9q34.3 | 6621 |
| SNAPC5 | small nuclear RNA activating complex polypeptide 5 | 15q22.31 | 10302 |

| | | | |
|---|---|---|---|
| SRSF6 | serine and arginine rich splicing factor 6 | 20q13.11 | 6431 |
| SSRP1 | structure specific recognition protein 1 | 11q12.1 | 6749 |
| STX3 | syntaxin 3 | 11q12.1 | 6809 |
| SUPT4H1 | SPT4 homolog, DSIF elongation factor subunit | 17q22 | 6827 |
| SUPT5H | SPT5 homolog, DSIF elongation factor subunit | 19q13.2 | 6829 |
| SURF1 | SURF1 cytochrome c oxidase assembly factor | 9q34.2 | 6834 |
| TAF10 | TATA-box binding protein associated factor 10 | 11p15.4 | 6881 |
| TAF12 | TATA-box binding protein associated factor 12 | 1p35.3 | 6883 |
| TAF13 | TATA-box binding protein associated factor 13 | 1p13.3 | 6884 |
| TAF1C | TATA-box binding protein associated factor, RNA polymerase I subunit C | 16q24.1 | 9013 |
| TAF6 | TATA-box binding protein associated factor 6 | 7q22.1 | 6878 |
| TAF9 | TATA-box binding protein associated factor 9 | 5q13.2 | 6880 |
| TARBP2 | TARBP2 subunit of RISC loading complex | 12q13.13 | 6895 |
| TK2 | thymidine kinase 2 | 16q21 | 7084 |
| TMED2 | transmembrane p24 trafficking protein 2 | 12q24.31 | 10959 |
| TP53 | tumor protein p53 | 17p13.1 | 7157 |
| TSG101 | tumor susceptibility 101 | 11p15.1 | 7251 |
| TYMS | thymidylate synthetase | 18p11.32 | 7298 |
| UMPS | uridine monophosphate synthetase | 3q21.2 | 7372 |
| UPF3B | UPF3B regulator of nonsense mediated mRNA decay | Xq24 | 65109 |
| USP11 | ubiquitin specific peptidase 11 | Xp11.3 | 8237 |
| VPS28 | VPS28 subunit of ESCRT-I | 8q24.3 | 51160 |
| VPS37B | VPS37B subunit of ESCRT-I | 12q24.31 | 79720 |
| VPS37D | VPS37D subunit of ESCRT-I | 7q11.23 | 155382 |
| XPC | XPC complex subunit, DNA damage recognition and repair factor | 3p25.1 | 7508 |
| XRCC2 | X-ray repair cross complementing 2 | 7q36.1 | 7516 |
| ZFP36L2 | ZFP36 ring finger protein like 2 | 2p21 | 678 |
| ZWINT | ZW10 interacting kinetochore protein | 10q21.1 | 11130 |

**Table S17. List of human DNA repair genes extracted from public human PDAC data sets.**
